# Omics analysis of MRSA under antibiotic stress identifies conserved adaptive modules and candidate adjuvant targets

**DOI:** 10.64898/2026.05.15.725322

**Authors:** Pedro C. Rosado, Pedro F. Pinheiro, M. Matilde Marques, Gonçalo C. Justino

## Abstract

Methicillin-resistant *Staphylococcus aureus* (MRSA) survives antibiotic exposure through coordinated physiological adaptation that extends beyond canonical resistance determinants. Here, we used an untargeted multi-omics strategy integrating proteomics, post-translational modifications, and metabolomics to characterize MRSA ATCC 43300 responses to five mechanistically distinct antibiotics: ampicillin, methicillin, vancomycin, chloramphenicol, and ciprofloxacin. Cells were exposed to 0.5×, 1×, and 2× IC_50_, enabling comparison of graded stress responses across antibiotic classes.

Across treatments, MRSA did not deploy fully distinct drug-specific programs. Instead, antibiotic exposure repeatedly converged on a limited set of conserved adaptive modules detectable across independent molecular layers. These included coupling of envelope stress with genome maintenance, recurrent remodeling of metal/cofactor and redox homeostasis, sustained pressure on nucleotide and folate metabolism, and reprogramming of transport and surface-associated functions. A particularly robust cross-antibiotic signature was the accumulation of MoO_2_-molybdopterin cofactor. Ciprofloxacin additionally induced compensatory envelope reinforcement, supporting tight coupling between DNA damage responses and cell-envelope maintenance.

Overall, the data support a unified model in which MRSA buffers mechanistically distinct antibiotic stress through a compact set of conserved stress-response functions rather than through entirely separate adaptive programs. These recurrent modules highlight candidate adjuvant vulnerabilities, particularly in metal/cofactor handling, nucleotide supply and repair, and transport/envelope compensation pathways. As this was an exploratory design intended to identify candidate adaptive patterns, these vulnerabilities now require validation in biologically replicated cultures and targeted functional studies.

## 1 Introduction

Antimicrobial resistance (AMR) represents one of the major alarming threats and is considered a critical global challenge. *Staphylococcus aureus* are Gram-positive bacteria responsible for several serious infections in humans, affecting mostly the skin and the respiratory system. The mortality rate of patients infected with these bacteria in the pre-antibiotic era was extremely high. Besides the desired therapeutic effect, the introduction of antibiotics against *S. aureus*, namely penicillin G and methicillin, have led to the emergence of strains resistant to these compounds. However, they are still effective against methicillin-susceptible *S. aureus* (MSSA) strains.

The emergence of methicillin-resistant *S. aureus* (MRSA) started in the 1960s, when methicillin-resistant isolates were detected worldwide, including in countries where methicillin was not available (1, 2).

Since then, MRSA has become one of the most clinically significant antibiotic-resistant pathogens and is included in the ESKAPE group (*Enterococcus faecium*, *S. aureus*, *Klebsiella pneumoniae*, *Acinetobacter baumannii*, *Pseudomonas aeruginosa* and *Enterobacter spp*.), a set of organisms responsible for the majority of life-threatening nosocomial infections and characterized by remarkable multidrug resistance mechanisms (3). Its inclusion in this group highlights the relevance of MRSA and its ability to evade both antimicrobial therapies and host defenses.

Nowadays, MRSA is a major cause of hospital- and community-acquired infections worldwide (4, 5). In Europe, MRSA accounts for more than 10 % of *S. aureus* invasive isolates in Portugal and in several southern European countries, and these strains are typically resistant to multiple antibiotics (4–6). Although methicillin is no longer clinically used, the term MRSA is still used to describe resistance to virtually all β-lactam antibiotics, including penicillins, cephalosporins, and carbapenems (1, 4). MRSA infections are often treated with glycopeptides such as vancomycin, since it is one of the most effective forms of treatment; nevertheless, vancomycin-resistant *S. aureus* (VRSA) strains have already been isolated, reinforcing the urgency to understand resistance mechanisms and develop innovative therapies (5, 7). Beyond its resistance phenotype, the epidemiological success of MRSA is also linked to the emergence of distinct evolutionary strains adapted to different environments. Traditionally, these are grouped into hospital-associated MRSA (HA-MRSA), community-associated MRSA (CA-MRSA) and livestock-associated MRSA (LA-MRSA). HA-MRSA strains are predominantly responsible for severe invasive infections such as bloodstream infections, endocarditis, and ventilator associated pneumonia, particularly affecting elderly and immunocompromised patients. In contrast, CA-MRSA strains frequently infect healthy individuals and are associated with skin and soft tissue infections. These strains often display enhanced virulence, driven by the expression of virulent factors that increase their pathogenic potential. LA-MRSA strains have emerged within intensive livestock production systems, and although primarily adapted to animal hosts, they have been increasingly identified as zoonotic agents capable of colonizing or infecting humans. The boundaries between these groups are becoming increasingly blurred as strains circulate across hospital, community, and livestock reservoirs (3, 7, 8). Clinically, MRSA infections are associated with high mortality rates, longer hospital stays and increased healthcare costs when compared with MSSA infections. Therapeutic options are significantly limited, and the emergence of strains with reduced susceptibility to glycopeptides or to last line antibiotics such as daptomycin and linezolid further undermines treatment efficacy (3). The spread of highly virulent and multidrug resistant strains increases the challenge, allowing MRSA to persist both in healthcare settings and in the community.

Although β-lactam resistance is a defining feature of MRSA, its adaptive capacity cannot be explained by *mecA* alone. Survival under antibiotic stress depends on coordinated regulatory and physiological remodeling involving cell envelope stress responses, metabolic adaptation, redox and biosynthetic balance, and global regulatory networks. Because different antibiotic classes impose distinct cellular constraints while also engaging overlapping survival programs, comparative systems-level analyses are needed to distinguish drug-specific effects from convergent adaptive responses and to identify vulnerabilities with potential therapeutic relevance (9–13).

Understanding antimicrobial resistance in MRSA requires an integrated view of bacterial physiology that extends beyond individual genes or pathways to the coordinated networks underlying adaptive survival. Multi-omics approaches, including genomics, transcriptomics, metabolomics, lipidomics, and proteomics, provide this system-level perspective by capturing the global molecular responses elicited by antibiotic exposure. Such strategies can reveal regulatory cascades, metabolic rewiring, stress-response programs, and compensatory adaptations that collectively shape resistance phenotypes. Integration across omics layers may therefore uncover overlooked resistance determinants, identify physiological vulnerabilities, and generate mechanistic hypotheses linking cellular state to antibiotic response (14–16). Ultimately, this framework supports a more rational basis for antimicrobial discovery by highlighting pathways and proteins with potential therapeutic relevance.

Here, we investigated the adaptive physiology of MRSA under antibiotic stress using a comparative multi-omics framework. Rather than focusing on a single resistance determinant or drug class, we aimed to define both antibiotic-specific and convergent responses across distinct antibacterial challenges, in order to identify metabolic and regulatory processes repeatedly associated with survival. In this way, the study was conceived not only to describe the molecular consequences of antibiotic exposure, but also to prioritize physiological vulnerabilities with potential therapeutic relevance.

For this purpose, MRSA was exposed to antibiotics representing different mechanistic classes and cellular constraints, namely the β-lactams methicillin and ampicillin, ciprofloxacin, chloramphenicol, and vancomycin. Together, these treatments provide a comparative framework to distinguish shared adaptive programs from drug-dependent effects. By integrating the corresponding omics signatures, we sought to uncover pathways and processes that are consistently mobilized during antibiotic adaptation and that may offer a rational basis for the future development of anti-MRSA intervention strategies. Accordingly, an 8 h exposure time was selected to prioritise the identification of adaptive physiological remodelling programs emerging under antibiotic pressure, rather than transient, early, or antibiotic-specific molecular responses.”

## 2 Experimental Procedures

Methicillin-resistant *Staphylococcus aureus* ATCC 43300 (MRSA) was used in this study. Bacteria were routinely cultivated at 37 °C with aeration in Mueller-Hinton broth or agar (MHB and MHA, Merck Millipore). Overnight pre-cultures were prepared by inoculating a single colony into liquid MHB and incubating for approximately 16 h at 37 °C with orbital shaking at 250 rpm. Antibiotic stock and work solutions were prepared in solvents appropriate for each compound: ampicillin (NZYTech) and vancomycin (ThermoScientific) in water, chloramphenicol (PanReac) and methicillin (ThermoScientific) in DMSO, and ciprofloxacin (Chemcruz) in 0.1 N HCl.

To establish the growth profile of MRSA, 50 mL of MHB in a 250 mL flask were inoculated with an overnight culture to an initial optical density at 640 nm (OD_640_) of approximately 0.02. The inoculated culture was incubated at 37 °C with orbital shaking at 250 rpm, and OD_640_ was monitored over time. Viable cell counts were determined by colony-forming unit (CFU) enumeration through serially dilution of culture aliquots; 10^-4^ to 10^-7^ dilutions were spread onto MHA plates. Plates were incubated at 37 °C for 24 h, after which colonies were counted, and viable cell concentrations were expressed as CFU/mL. The antibacterial activity of test compounds against MRSA was evaluated by determining minimum inhibitory concentration (MIC) and half-maximal inhibitory concentration (IC_50_) values using an adapted broth microdilution assay. Overnight cultures grown in MHB were diluted in fresh MHB to OD_640_ = 0.02; 500 µL of the diluted bacterial suspension were added to 495 µL of fresh MHB and 5 µL of compound solution in 15 mL tubes. Compounds were tested across concentrations ranging from 1 mM to 1 nM using serial dilutions. In some experiments, the assay was performed in 96-well plates by combining 1 µL of compound solution, 99 µL of fresh MHB, and 100 µL of diluted bacterial suspension per well. Cultures were incubated at 37 °C for 24 h.

### 2.1 Antibiotic treatment of MRSA cultures for omics experiments

For omics experiments evaluating antibiotic-induced responses in MRSA, overnight cultures were diluted in fresh MHB to OD_640_ = 0.02 and used to inoculate 5 mL cultures prepared in 50 mL tubes. Each culture contained 4.95 mL of MHB and 50 µL of antibiotic solution, corresponding to final concentrations of 0.5×, 1×, or 2× the previously determined IC_50_ value. Cultures were incubated at 37 °C with orbital shaking for 8 h. Growth was monitored by OD_640_, and culture biomass was normalized using the established conversion in which OD_640_ = 1.0 corresponded to approximately 6 × 10^9^ viable cells/mL. Aliquots corresponding to 10^8^ cells were collected, and cells were pelleted by centrifugation at 16 000 × *g* for 10 min for downstream omics workflows.

### 2.2 Sample preparation

At each time point, growth was monitored by OD_640_, biomass was normalized using the established OD-to-viable cell number relationship, and cell pellets and supernatants were collected for downstream omics analyses. Samples were analyzed by ultra-high performance liquid chromatography coupled to high resolution mass spectrometry (UHPLC/HRMS)-based metabolomic, and proteomic workflows, and resulting datasets were analyzed.

#### 2.2.1 Metabolomics

A metabolomics approach was employed to characterize the intracellular and extracellular metabolomes (endometabolome and exometabolome, respectively) of bacterial cells, reflecting both intracellular activity and secreted metabolic products. For this purpose, different extraction methods were employed to selectively isolate polar and non-polar metabolites from culture media and cell pellets. These protocols were adapted and optimized based on published methodologies (17, 18). To analyze the exometabolome, two types of extractions were performed. In the first method, 400 μL of cold methanol was added to 100 μL of culture medium, followed by vortexing and centrifugation at 24000 × *g*. The supernatants were stored at -20 °C for further LC-MS analysis. In the second approach, 1 mL of ethyl acetate was added to 1 mL of culture medium, followed by vortexing. After phase separation, the organic phase was collected and dried using a speed vacuum concentrator. The samples were then resuspended in 100 μL of methanol:water (3:2, v/v) containing 100 μM of L -Tryptophan-(indole-*d_5_*) 98 % (Cambridge Isotope Laboratories, Inc), 2 μM butadiene sulfone 98 % (Aldrich), and 20 μM 1,2-dipalmitoyl-*rac*-glycerol (Sigma-Aldrich). The endometabolome was extracted by resuspending cell pellets in 250 μL of methanol:water (3:2, v/v) containing 100 μM of L -Tryptophan-(indole-*d_5_*) 98 %, 2 μM butadiene sulfone 98 %, and 20 μM 1,2-dipalmitoyl-*rac*-glycerol. Three freeze-thaw cycles were performed using liquid nitrogen. Samples were vortexed, placed in an ultrasonic bath for 5 minutes, and then transferred to ice. This step was repeated twice, followed by the addition of 250 μL of chloroform. Samples were centrifuged at 14000 × *g* for 10 minutes at 4 °C. The polar phase (top layer) was collected and stored at -20 °C for LC-MS analysis. An additional apolar-oriented metabolomic extraction was performed to extend molecular coverage towards hydrophobic and lipid-associated metabolites.

#### 2.2.2 Proteomics

Proteomic profiling enables comprehensive study of protein expression and post-translational modifications. Both the endoproteome (intracellular proteins) and exoproteome (secreted proteins) were extracted using optimized lysis and precipitation protocols to ensure high protein yield and integrity. Protein samples were processed for downstream LC-MS analysis using standardized workflows (18, 19). To cover the exoproteome, the culture medium total protein content was extracted by adding four volumes of cold acetone (- 20 °C), followed by vortexing. Samples were incubated at - 20 °C for 16 hours and centrifuged at 15000 X *g* for 10 minutes (19). The supernatant was carefully discarded. Samples were left open at room temperature for 30 minutes to allow residual acetone to evaporate. Further processing was carried out following an optimized filter-aided sample preparation (FASP) protocol (20–23). For endoproteomics, a 250 μL volume of lysis buffer (0.1 M Tris-HCl pH 7.8, 0.05 M DTT, 2 %SDS, 0.1 mg/mL lysozyme) was added to each cell pellet or protein precipitate (from exoproteome extraction). Three freeze-thaw cycles with liquid nitrogen were performed. Samples were sonicated for 2 minutes and then incubated at 95 °C for 5 minutes. After cooling to room temperature, samples were centrifuged at 16000 X *g* for 10 minutes. The supernatant was quantified using the Bradford assay, and 50 μg of protein from each sample were used for further processing employing an optimized FASP protocol (20–23).

*Filter-aided sample preparation:* Samples were loaded into 3 kDa MWCO regenerated cellulose spin concentrators (amicon-ultracel 3, Sigma-Aldrich). In all steps, except the final elution steps, the eluates were discarded. Samples were centrifuged at 12000 X *g* for 30 minutes at 20 °C. Then, 200 μL of buffer containing 25 mM ammonium bicarbonate and 8 M urea (Merck) were added, followed by another centrifugation under same conditions. A volume of 100 μL of 25 mM ammonium bicarbonate, 8 M urea, and 50 mM iodoacetamide (Sigma) was then added. Samples were incubated in the dark for 20 minutes, and centrifuged at 12000 X *g* for 30 minutes, at 20 °C. Samples were washed twice with 200 μL of 25 mM ammonium bicarbonate. Membranes were transferred to clean collection tubes, and 100 μL of a trypsin solution (1:30 ratio (w/w), trypsin sequencing grade from bovine pancreas, Roche) in 12.5 mM ammonium bicarbonate were added. Samples were incubated for 16 hours at 37 °C. After incubation, samples were sonicated for 2 minutes and centrifuged at 12000 X *g* for 30 minutes at 20 °C. The eluate was retained. Then, 50 μL of 3 % acetonitrile and 0.1 % formic acid were added, followed by another centrifugation under the same conditions. The eluate was retained and combined with the previous one. This step was repeated once more. The final eluate (200 μL) was stored at - 20 °C for LC-MS analysis.

### 2.3 UHPLC/HRMS analysis

HRMS analysis was carried out for each sample in triplicate using an Elute UHPLC system coupled to an Impact II QqTOF mass spectrometer with an electrospray ionization (ESI) source (Bruker Daltonics GmbH & Co) (UHPLC-ESI-HRMS). Data acquisition was performed using optimized pre-established methods (17, 18). Metabolomic samples were analyzed in both positive and negative modes, coupled to reversed-phase (RP) chromatography and hydrophilic interaction liquid chromatography (HILIC).

For all studies, initial calibration of the mass spectrometer was performed though direct infusion with the respective calibrant introduced in the ion source via a 20 μL loop using the high-precision calibration mode (HPC), with a search range of ± 0.1 ppm relative to each expected ion *m/z* and with a standard deviation below 0.08 ppm for the set of identified ions with an intensity threshold of 1000, ensuring a score above 99.95%, except where mentioned. For metabolomics studies the calibrant was1 mM sodium formate/acetate solution (125:125:375:125:25 H_2_O:IPA:AcOH:HCOOH:NaOH 1 M, v/v/v/v/v); for all other studies, calibrant 0.1 % sodium trifluoracetic solution (in water) titrated to pH 3.5 with 1 M NaOH and introduced to the ion source via a 20 μL loop using the high-precision calibration mode. For proteomics, calibrant was ESI-L Low Concentration Tuning-Mix (Agilent, USA), and calibration was performed using the enhanced quadratic calibration (EQC) mode. For internal calibration in each sample, an initial 0.5 s calibration segment was acquired,

#### 2.3.1 Metabolomics

Metabolites were separated by reverse-phase UHPLC on a Luna 2.5 µm C18(2)-HST column (100 Å, 150 x 2 mm, Phenomenex) at a constant temperature of 40 °C, using a gradient elution at a flow rate of 250 μL/min (mobile phase A: 0.1 % (v/v) formic acid in water; mobile phase B: 0.1 % (v/v) formic acid in acetonitrile): 0.0-0.5 min, 0 % B; 0.5-1.5 min, 0 to 20 % B; 1.5-4.0 min, 20 to 60 % B; 4.0-6.0 min, 60 to 100 % B; 6.0-9.0 min, 100 % B; 9.0-10.0 min, 100 to 0 % B, followed by a 5 min column re-equilibration step. For hydrophilic interaction liquid chromatography (HILIC), an XBridge BEH Amide XP Column (130 Å, 2.5 µm, 150 x 2.1 mm, Waters) was used at a constant temperature of 40 °C. With a flow rate of 250 μL/min, a gradient elution of 10 mM ammonium acetate in water containing 0.1 % (v/v) acetic acid (A) and 10 mM ammonium acetate in acetonitrile:water:acetic acid 980:20:1 (v/v/v) (B) was applied: 0-2 min, 90 % B; 2-6 min, 90 to 70 % B; 6-9 min, 70 to 30 % B; 9-13 min, 30 % B; 13-18 min, 30 to 90 % B, followed by a 4 min column re-equilibration step.

High resolution mass spectra were acquired in both electrospray ionization modes, with the following acquisition parameters: capillary voltage, 4.5 kV (ESI+) or 3 kV (ESI-); end plate offset, 500 V; nebulizer, 2.0 bar; dry gas (N_2_) flow, 8.0 L/min; dry heater temperature, 220 °C. The tune parameters were set according to: transfer funnel 1/2 RF power (150/200 Vpp), hexapole RF power (50 Vpp), ion energy (4.0 eV), low mass (90 *m/z*), collision energy (7.0 eV), collision RF power (650 Vpp), transfer time (80 μs), pre-pulse storage (5 μs). Spectra acquisition was performed with an absolute threshold of 25 counts per 1000. For auto MS/MS mode, spectra were acquired with a threshold of 20 counts per 1000, cycle time of 3.0 seconds with exclusion after 3 spectra and release after 1.00 min. All acquisitions were performed with an *m/z* range from 70 to 1000 and with a 3 Hz spectra rate. Three full scans and 1 auto MS/MS scan were performed for each sample using both positive and negative ionisation modes.

#### 2.3.2 Untargeted proteomics

Peptides were separated on a reverse-phase bioZen™ 2.6 µm Peptide XB-C18 column (100 Å, 100 x 2.1 mm, Phenomenex), at a constant temperature of 45 °C, using a gradient elution at a flow rate of 300 μL/min (mobile phase A: 0.1 % (v/v) formic acid in water; mobile phase B: 0.1 % (v/v) formic acid in acetonitrile): 0-2 min, 0 % B; 2-5 min, 1 % B; 5-60 min, 1 to 50 % B; 60-65 min, 50 % B; 65-70 min, 50 to 95 % B; 70-78 min, 95 % B; 78-85 min, 95 to 1 % B, followed by a 5 min column re-equilibration step. After sample loading, a simple on-line desalting step was implemented in the first 2 minutes of elution (0 % B): using the mass spectrometer six-port valve, the flow was sent to waste, avoiding contamination of the MS instrument.

MS acquisition parameters were set as follows: capillary voltage of 4.5 kV (ESI+), with an end plate offset of 500 V, a nebulizer pressure of 2.5 bar, a dry gas (N_2_) flow of 8.0 L/min and a heater temperature of 200 °C. The tune parameters were set according to: transfer funnel 1/2 RF power (400/600 Vpp), hexapole RF power (400 Vpp), ion energy (5.0 eV), low mass (200 *m/z*), collision energy (7.0 eV), pre pulse storage (5 μs), stepping (on, basic), collision RF power (200-1200 Vpp), transfer time (50-110 μs), timing (50 %), collision energy (100-120 %). Spectra acquisition was performed in auto MS/MS mode with a threshold of 28 counts per 1000, cycle time of 3.0 seconds with exclusion after 1 spectrum and release after 0.50 min. All acquisitions were performed with an *m/z* range from 150 to 2200 and a 2 Hz spectra rate. After every 20 injections, a bovine serum albumin tryptic digest was analyzed to monitor retention time variation and base peak chromatogram intensity. These were maintained below 10 s and 10%, respectively, across all runs to ensure the consistency of the analyses.

### 2.4 Experimental Design and Statistical Rational

Each condition comprised one bacterial sample analyzed in triplicate by LC–MS in each omics layer. For the proteomics workflow, 37 samples were analyzed in total, including 19 exoproteome samples and 18 cellular proteome samples; these comprised solvent/control conditions and antibiotic-treated samples across the tested concentrations. Given the controlled nature of the bacterial culture system, this work was structured as a first-pass discovery experiment prioritizing depth of molecular characterization and analytical reproducibility within each condition. The resulting omics changes are interpreted as candidate condition-associated responses requiring confirmation in biologically replicated experiments. For each sample, control growths were performed replacing the antibiotic with the solvent used to prepare antibiotic solutions (water for ampicillin and vancomycin, DMSO for chloramphenicol and methicillin, 0.1 M HCl for ciprofloxacin). For exoproteomics and exometabolomics, culture media (MHB) was used as background control. No acquired data was excluded from downstream analyses. Statistical comparisons between each treated sample and its corresponding control were performed on log_2_-transformed feature intensities using fold-change (FC) analysis combined with two-sided Welch’s *t*-tests across the three technical replicate measurements. Welch’s *t*-test was selected because it does not assume equal variance between groups and is commonly used for small-sample pairwise comparisons in quantitative omics workflows. Differential abundance was defined using a stringent effect-size threshold of |log_2_FC| ≥ 1.5 together with *p* < 0.05. This cutoff was selected as a conservative filter to prioritize large-magnitude changes in this exploratory dataset. Because the replicate structure was technical rather than biological, these *p*-values are used here as measures of analytical reproducibility and signal consistency, not as population-level estimates of biological significance. Functional enrichment analyses in STRING (24) and XCMS (25, 26) were interpreted using false discovery rate (FDR)-adjusted significance values as provided by the respective software platforms, with a 0.05 FDR cut-off.

### 2.5 Data Analysis

Metabolomics data was processed, analyzed, and interpreted using the XCMS platform (25–29) applying the parameters presented in Table S1 for UHPLC/QqTOF and SAUR282458 as the biosource. Pairwise analysis of both LC columns and ionization modes were integrated through a multimodal analysis (30) using a 0.05 *p*-value cutoff.

Proteomics data analysis was performed using MaxQuant v2.4.2.0 followed by statistical analysis in Perseus v1.6.15.0 (31, 32). Acquired raw data files were processed with MaxQuant using the internal search engine Andromeda (33), and protein sequences from the Uniprot database (34). The whole proteome was used for *Staphylococcus aureus* ATCC 43300 (UP000244076, 2737 proteins), the primary reference due to its clinical relevance and resistance phenotype), *Staphylococcus aureus* strain NCTC 8325/PS47 (UP000249159, 2614 proteins), a widely studied laboratory strain with extensive genomic and proteomic annotations), and *Staphylococcus aureus subsp. aureus* 71193 (UP000005003, 2620 proteins), included to broaden coverage and account for potential strain-specific protein variation.

A curated panel of MRSA proteins implicated in virulence regulation, antibiotic resistance, and stress response was built (1396 sequences, including canonical sequences and known isoforms), collecting the sequences of *agrA/B/C/D, arlRS, blaZ, ccpA, clpP, cvfA, femA, fhs, hld, hlyA, mgrA, psmA (all sequences), msa, pbp* (all sequences), *qnr, sarA/U/X, spoVG, spsAB, and ssrAB*, as well as all sequences corresponding to proteins annotated as “shikimate kinase” and as “sigma factor” (35–58). This strategy enables evaluating whether established resistance and virulence determinants were differentially expressed under antibiotic stress, providing complementary biological validation of the proteomics results.

Database searches were performed against the selected *S. aureus* proteomes; all proteins from other taxonomies were not considered. During downstream filtering, protein groups annotated by MaxQuant as ‘Potential contaminant’ (standard MaxQuant database), ‘Reverse’, or ‘Only identified by site’ were removed before statistical analysis, network analysis, and biological interpretation. Database searching in MaxQuant/Andromeda used a first-search precursor tolerance of 20 ppm, a main-search precursor tolerance of 10 ppm, and a fragment ion tolerance of 40 ppm under the TOF setting. Peptide-spectrum matches were accepted by target-decoy filtering at 1% peptide-level false discovery rate (FDR), while protein groups and modification sites were filtered at 1% FDR. No additional manual Andromeda score or expectation-value cutoff was applied, because identification confidence was controlled through FDR-based filtering. This approach was selected as a standard, statistically controlled acceptance criterion for discovery proteomics.

For trypsin digestion experiments, carbamidomethylation of cysteine (+57.0215 Da) was set as a fixed modification, whereas oxidation of methionine (+15.9949 Da), protein *N*-terminal acetylation (+42.0106 Da), phosphorylation of serine, threonine, tyrosine, and histidine (+80.9736 Da), deamidation of asparagine (+0.9840 Da), succinylation of lysine (+100.0160 Da), and ubiquitination of lysine were defined as variable modifications. Bruker Q-TOF was selected as instrument and its default parameters were used. Enzyme specificity was set to trypsin/P, allowing cleavages at the carboxyl side of arginine and lysine residues and before proline residues, with a maximum of 2 missed cleavages allowed. The false discovery rate for peptides and proteins was set to 1 %, and the minimum score and delta score for modified peptides were set to 0. Match between runs was enabled with a 0.7 min match time window and a 20 min alignment window. Dependent peptides were enabled with a fold discovery rate of 1 %. Normalised spectral protein label-free quantification (LFQ) intensities were calculated using the MaxLFQ algorithm (59), with a minimum ratio count of 1. For experiments without digestion, the fixed modification carbamidomethylation of cysteine was removed. After filtering, proteins classified as only identified by site, contaminants, and reverse hits were removed. Protein group LFQ intensities were log-2-transformed, and the quantitative profiles were filtered from missing values with a minimum valid number of “(number of replicates/2)+1” in at least one group. Missing values were replaced using a normal distribution (width 0.3 and down shift 1.8). Log_2_ fold changes were calculated from the difference in mean log_2_(LFQ) intensities between the compared conditions. Statistical testing was performed in Perseus using a two-sided Student’s *t*-test with *p* < 0.05 and S0 = 0. Significantly altered proteins were used for network enrichment in STRING (24) using the reference database for *Staphylococcus aureus* subsp. *aureus* NCTC 8325, at an FDR cut-off of 0.05.

## 3 Results and discussion

To characterize the molecular consequences of antibiotic pressure in MRSA, a comprehensive untargeted mass spectrometry-based multi-omics strategy was implemented. This integrative workflow, combining proteomics, and metabolomics across intracellular and extracellular fractions, was designed to resolve antibiotic-induced perturbations at multiple layers of cellular physiology. By mapping changes in metabolic pathways and protein networks, this approach provides a systems-level view of MRSA adaptation to ampicillin, chloramphenicol, ciprofloxacin, methicillin, and vancomycin. Beyond a descriptive profiling, the integration of these datasets enables the identification of mechanistically relevant pathways and molecular nodes that may constitute actionable vulnerabilities for overcoming antibiotic resistance.

Antibiotic susceptibility was assessed in MRSA by determining IC_50_ values for each compound: 17.2 µM for ampicillin, 12.5 µM for chloramphenicol, 0.38 µM for ciprofloxacin, 12.85 µM for methicillin, and 0.75 µM for vancomycin. MRSA was exposed at 0.5, 1, and 2xIC_50_ concentrations, at 8 h of growth.

### 3.1 Proteomics changes associated to antibiotic action and resistance

A proteomic analysis was conducted as one of the layers of a multi-omics approach to gain insight into protein-level adaptation of MRSA to antibiotic stress.

#### 3.1.1 Ampicillin

Ampicillin is a β-lactam antibiotic that targets bacterial cell wall biosynthesis by binding to PBPs and inhibiting their activity (2). However, *S. aureus* has developed resistance to this antibiotic through the production of penicillinase, an enzyme that hydrolyzes the β-lactam ring (2, 60). The proteomic profiling of MRSA treated with ampicillin revealed a set of significantly dysregulated proteins across the three tested concentrations (Figure S1).

Ampicillin significantly dysregulated several cell wall biosynthesis proteins, including penicillin-binding protein 3 (*pbp3* or *pbpC*, log_2_FC = +1.80 at 0.5× IC_50_), a direct β-lactam target that catalyzes late peptidoglycan synthesis (61); lipid II:glycine glycyltransferase (*femX*, log_2_FC = +1.73 at 0.5× IC_50_), required for glycine incorporation into the pentaglycine interpeptide bridge and linked to β-lactam resistance (62–64); teichoic acid D-alanine hydrolase (*fmtA*, log_2_FC = -1.59 and -1.61 at 0.5× and 1× IC_50_, respectively), which modifies wall teichoic acids and contributes to methicillin resistance (65); and peptidoglycan hydrolase (Q2FWV4, log_2_FC = -1.71 at 2× IC_50_), involved in peptidoglycan remodeling, turnover, and recycling (66).

DNA repair and genome stability proteins were also up-regulated, including DNA ligase (*ligA*, log_2_FC = +2.80 and +3.04 at 0.5× and 2× IC_50_, respectively) which seals nicks during DNA replication and repair (67); RadA (log_2_FC = +1.94 at 0.5× IC_50_), involved in homologous recombination and repair (68, 69); and . ribonucleoside-diphosphate reductase (Q2G078, log_2_FC = +3.32 at 0.5× IC_50_), which supplies deoxyribonucleotides and supports survival under stress (70). These findings suggest that ampicillin affects not only cell envelope processes but also DNA integrity, requiring repair responses to support persistence. Phage-associated proteins, including HK97 portal protein family members, were also up-regulated, consistent with prophage mobilization, horizontal gene transfer, and virulence regulation in *S. aureus* (71).

Several uncharacterized proteins were also dysregulated and probed by BLASTp (72) Q2FW89 (log_2_FC = +6.10 and 6.11 at 1× and 2× IC_50_, respectively) showed high similarity to the teicoplanin resistance-associated HRH-type transcriptional regulator TcaR (COV = 59%, ID = 100%), a transcription factor linked to antibiotic responses and biofilm regulation in *S. epidermidis* (73, 74). Q2G057 (log_2_FC = -1.55 at 0.5× IC_50_) was highly similar to a DEAD/DEAH box helicase (COV = 100%, ID = 98.7%), a component of the *S. aureus* RNA degradosome (75, 76). Since efficient RNA degradation supports rapid *agr* adaptation, its down-regulation may favor biofilm formation and persistence, important survival strategies under antibiotic stress (77, 78). Q2G0R7, down-regulated at all concentrations (log2FC = -2.74, -2.99 and -2.79 at 0.5×, 1× and 2× IC50, respectively) was similar to a polysaccharide biosynthesis protein (COV = 100%, ID = 99.2%) and was consistently down-regulated across all ampicillin concentrations, suggesting altered cell-envelope or biofilm-associated functions (79, 80). By contrast, Q2FXC1 (log_2_FC = +1.75 at 1× IC_50_) and Q2G0Z3 (log_2_FC = +2.36 at 2xIC_50_) proteins could not be annotated.

Five proteins were up-regulated at least two ampicillin concentrations. DNA ligase, which supports DNA replication and repair (81), was induced (log_2_FC =+2.80 and +3.04 at 0.5× and 2× IC_50_, respectively), underscoring the importance of genomic stability under β-lactam stress. Ribonuclease P (*rnpA*), essential for tRNA maturation, was also induced (log_2_FC = +3.16 and +1.93 at 0.5× and 1× IC_50_, respectively), indicating the need to sustain translation under antibiotic stress; notably, it has been proposed as an antimicrobial target (82). The phage portal protein (HK97) was also consistently up-regulated (log_2_FC = +1.82 and +1.75 at 0.5× and 1× IC_50_, respectively).

Interestingly, two poorly annotated proteins were up-regulated in multiple ampicillin conditions: Q2FW89 (log_2_FC = +6.10 and +6.11 at 1× and 2× IC_50_, respectively) and FKLRK (log_2_FC = +1.61 and 1.66 at 0.5× and 1× IC_50_, respectively). FKLRK is described as a cell surface protein whose induction has been observed under biofilm-promoting conditions in MRSA. Together, these patterns suggest that both proteins may contribute to ampicillin adaptation, potentially through biofilm-associated modulation of cell surface functions in *S. aureus* (83–85).

To place these individual protein responses within a broader biological context, a protein-protein interaction (PPI) analysis (Figure S2) was carried out using STRING v12.0 (24).

At 0.5× IC50, enrichment involved transcriptional/regulatory proteins, AAA-domain proteins, HNHc nucleases, and DNA replication-associated factors, including DNA polymerase A and PD-(D/E)XK nucleases, indicating early adaptive responses centered on transcriptional modulation, DNA repair, protein quality control, and stress sensing that help preserve genomic and proteomic stability. Enrichment of ComK- and sigma-70-related functions further supports activation of stress-responsive regulatory circuits that may promote rapid adaptation (70, 86, 87).

At 1× IC50, STRING enrichment identified nucleotide excision repair, dUTPase, DNA polymerase A, and PD-(D/E)XK nuclease-associated functions, indicating a stronger genome-maintenance response. Because nucleotide excision repair removes bulky DNA lesions and dUTPase prevents uracil incorporation into DNA, these data support a more defined DNA damage-management program under intermediate ampicillin stress that may promote MRSA persistence (81, 88–90).

At 2× IC50, enrichment shifted toward siderophore-dependent iron import and glycerophospholipid metabolism, indicating broader metabolic and envelope remodeling. Consistently, the proteome showed up-regulation of an inositol monophosphatase family protein (Q2FVV7) and down-regulation of an Fe/B12 periplasmic-binding domain-containing protein (Q2FWN6), supporting roles for metal homeostasis, phospholipid remodeling, and broader metabolic adaptation in response to high ampicillin stress (91–97).

Proteins dysregulated in at least two conditions were also analyzed as a separate network (Figure 1). This network resolved into local clusters associated with ester-bond hydrolase activity, phosphodiesterase activity, and glycerophospholipid metabolism.

**Figure 1.**
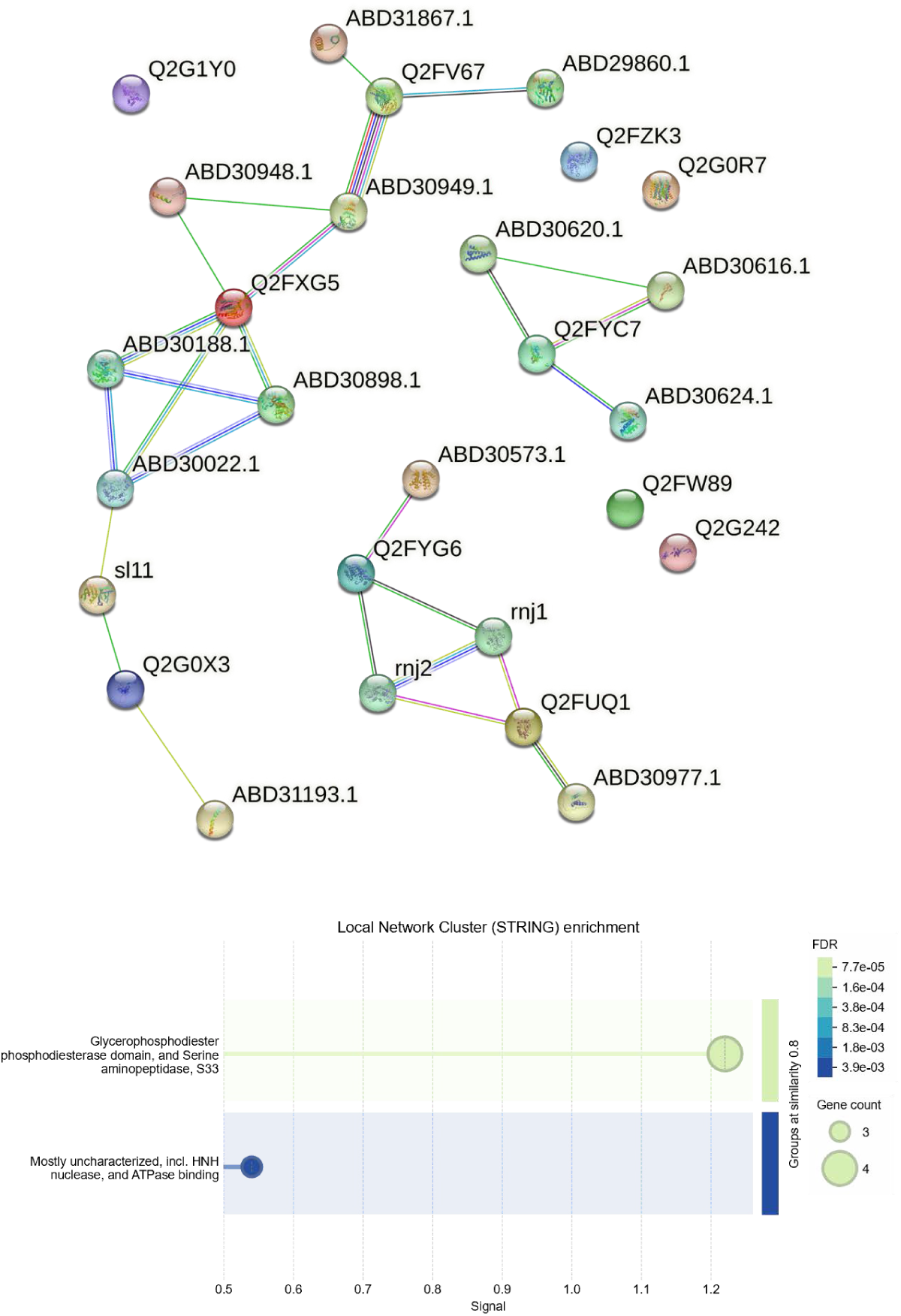
STRING analysis of proteins dysregulated in at least two ampicillin conditions. Upper panel: protein–protein interaction network generated from proteins significantly altered in at least two of the three ampicillin concentrations tested (0.5×, 1×, and 2× IC50), with first-shell interactors added for network contextualization. Lower panel: functional enrichment summary of the same network showing significantly enriched at FDR ≤ 0.05.

Recurrent down-regulation of the serine aminopeptidase S33 domain-containing protein Q2FXG5 and the uncharacterized protein Q2G0R7 across all three concentrations indicates conserved suppression under ampicillin stress and suggests that these proteins may represent recurrently affected adaptive nodes. Overall, the data support a concentration-dependent proteomic response to ampicillin.

While at 0.5× IC_50_, MRSA appears to engage early and potentially reversible stress programs centered on regulatory adjustment and genome surveillance, at 1× IC_50_, the response shifts toward active DNA repair and nucleotide homeostasis, and at 2× IC_50_ the adaptive program becomes more metabolic, involving lipid remodeling, metal homeostasis, and alternative catabolic pathways. Across concentrations, the data also suggest participation of biofilm-associated survival strategies, indicating that MRSA combines shared persistence mechanisms with dose-specific adaptations to withstand ampicillin stress.

In addition to the network-based analyses, the proteomic dataset was cross-referenced against a curated panel of MRSA proteins associated with virulence regulation, antibiotic resistance, and stress response. For ampicillin, none of the proteins in this panel were consistently dysregulated across multiple concentrations, suggesting that the adaptive response primarily involves other processes.

#### 3.1.2 Chloramphenicol

Chloramphenicol targets bacterial protein biosynthesis by binding to the 50S ribosomal subunit of the 70S ribosomes, inhibiting peptide chain elongation (98). Resistant strains of *S. aureus* to chloramphenicol have been reported due to the expression of chloramphenicol acetyltransferases (CATs), or by reducing intracellular concentration of the drug which is related to efflux pumps.

Exposure to chloramphenicol significantly altered protein expression of 33 proteins (Figure S3) across sub-inhibitory to inhibitory concentrations (0.5×, 1×, and 2× IC_50_). DNA repair and replication modules were strongly affected. Both a DNA repair/chromosome segregation ATPase (Q2FVZ7, log_2_FC = +1.95 and +2.54 at 0.5× and 1× IC_50_, respectively) and the chromosome partitioning protein Smc (*smc*, Q2FZ49, log_2_FC = +2.02 at 2× IC_50_) were induced, together with NrdI (*nrdI*, Q2G079, log_2_FC = +2.74 and 2.57 at 1× and 2× IC_50_, respectively). In parallel, NrdI supports ribonucleotide reductase activity, sustaining ribonucleotide reduction and dNTP supply (99).

Proteins related to translation and RNA metabolism were also dysregulated. Most notably, the *argS* gene (Q2G0F8), which encodes arginine-tRNA ligase, involved in arginine-tRNA aminoacylation (100), was down-regulated (log_2_FC = -2.63 at 2× IC_50_) at the highest antibiotic concentration, consistent with chloramphenicol’s primary action on the ribosome (98). Taken together, these changes suggest that impaired translation under chloramphenicol imposes collateral replication stress, necessitating stabilization of chromosome architecture and nucleotide metabolism to ensure survival.

Central metabolic enzymes showed a coordinated remodeling. Fatty acid catabolism was inhibited, with acyl-CoA dehydrogenase (Q2FVW7, log_2_FC = -5.22, -4.42, -5.56 at 0.5×, 1× and 2× IC_50_, respectively) strongly down-regulated across all conditions. In contrast, compensatory changes included induction of dihydrolipoamide acetyltransferase of the pyruvate dehydrogenase complex (Q2FY54, log_2_FC = +1.72 at 1× IC_50_) and enoyl-CoA (Q2G1C9, log_2_FC = +2.23 at 1× IC_50_), suggesting a metabolic rerouting from β-oxidation toward pyruvate metabolism and alternative energy sources. These changes are accompanied by the down-regulation of dihydrolipoyllysine-residue succinyltransferase component of 2-oxoglutarate dehydrogenase complex (*odhB*, Q2FYM2, log_2_FC = -2.68 at 2× IC_50_), indicating a remodeling of the TCA cycle. This is notable because *S. aureus* relies on amino acids such as alanine, aspartate, glutamate, and proline to fuel pyruvate oxaloacetate, and TCA intermediates during host infection (96). Disruption of 2-oxoglutarate dehydrogenase activity (*sucA*/*sucB*) has also been linked to reduced membrane potential and ATP levels, which in turn promotes persister formation and antimicrobial tolerance (101). Thus, the observed down-regulation of the dihydrolipoyllysine-residue succinyltransferase component of the 2-oxoglutarate dehydrogenase complex suggests a possible shift toward persistence-like physiology under chloramphenicol stress.

Increased abundance of the glutathione hydrolase proenzyme (Q2G1F4, log_2_FC = +1.86 at 0.5× IC_50_) and the CobW C-terminal domain containing protein (Q2G0W4, log_2_FC = +2.02 at 0.5× IC_50_) further implies redox stress management under chloramphenicol stress. Glutathione and related cysteine-containing molecules such as thioredoxin are central to *S. aureus* oxidative stress defenses (9) . Perturbation of glutathione levels has also been associated with increased oxidative DNA damage and activation of SOS responses, which can contribute to resistance evolution (102).

Transport and membrane associated proteins were also impacted. Several uptake systems, including an AEC family transporter (Q2FVZ8, log_2_FC = +2.59 and +2.83 at 1× and 2× IC_50_, respectively), an amino acid ABC transporter (Q2FX87, log_2_FC = +1.57 and +2.66 at 1× and 2× IC_50_, respectively), and the EIICB component of the lactose-specific phosphotransferase system (PTS) system (Q2G2D4, log_2_FC = +2.36 at 2× IC_50_), were induced, reflecting increased demand for external nutrients under translational inhibition. Amino acid transporters are among the most abundant membrane proteins in *S. aureus* and are known to play critical roles not only in nutrient acquisition but also in virulence and antibiotic resistance, with previous studies suggesting that interfering with amino acid permeases can impair metabolic homeostasis and pathogenicity (103, 104). The induction of the PTS system lactose-specific EIICB component aligns with previous findings that PTS transporters couple sugar uptake to phosphorylation, directly integrating nutrient sensing with central metabolism (105, 106) Collectively, the observed induction of nutrient transport systems under chloramphenicol stress supports the notion that *S. aureus* relies on enhanced environmental nutrient acquisition to sustain metabolism when protein synthesis is impaired.

Conversely, a major facilitator superfamily (MFS) transporter (Q2G0C4, log_2_FC = -1.66 at 1× IC_50_) was repressed. MFS proteins constitute one of the largest classes of bacterial transporters, with broad substrate specificity, ranging from sugars and amino acids to antimicrobial agents (107). Importantly, several MFS members in *S. aureus* act as multidrug efflux pumps that contribute to antibiotic resistance and are, therefore, considered promising targets for efflux pump inhibitors aimed at restoring antimicrobial efficacy (108). The down-regulation observed in this protein can be potentially part of the strategy to conserve energy and redirect membrane transport. Together, the induction of multiple uptake systems and the repression of an MFS efflux transporter suggest a strategic remodeling of membrane transport under chloramphenicol stress favoring an energy-conserving adaptation.

Proteins linked to the cell envelope response were also altered: surface protein F (*sasF*, Q2FUW9, log_2_FC = +2.23 at 2× IC_50_), lipoprotein signal peptidase (*lspA*, Q2FZ79, log_2_FC = +1.92 at 2× IC_50_), and a LiaF-like sensor (Q2FX07, log_2_FC = +1.52 at 1× IC_50_) were up-regulated, suggesting remodeling of the staphylococcal surfaceome.

While SasF remains functionally uncharacterized, it belongs to the LPXTG-anchored family of surface proteins that serve as essential virulence factors and vaccine targets in *S. aureus* (109, 110). LspA, required for lipoprotein maturation and full virulence in Gram-positive bacteria, is also considered a promising anti virulence drug target (111). The LiaF-like protein is associated with the VraTSR regulatory system, which mediates resistance to cell wall-active antibiotics, suggesting stress-induced reinforcement of envelope defense pathways (112, 113). In contrast, down-regulation of the transcriptional regulator HptR (Q2G1E1, log_2_FC = -1.71 at 0.5× IC_50_), normally involved in hexose-phosphate uptake and envelope stress signaling, points to a reorganization of metabolic control under chloramphenicol stress (114). The induction of the FKLRK surface protein (Q2G0X3, log_2_FC = +2.29 and +2.01 at 1× and 2× IC_50_, respectively), previously associated with biofilm induction (115), further suggests that exposure to chloramphenicol may promote biofilm-associated adaptation.

Phage-associated proteins were also induced, including a hypothetical phage protein (Q2FY90, log_2_FC = +1.74 and +2.89 at 1× and 2× IC_50_, respectively), and a conserved phage protein (Q2FY93, log_2_FC = +3.46 at 0.5× IC_50_). This parallels observations under ampicillin, suggesting that activation of prophage elements is a general stress response that enhances genetic plasticity (116, 117).

Finally, several poorly annotated or uncharacterized proteins, were clarified through BLASTp (72) searches, revealing potential roles in chloramphenicol adaptation. The Q2FW76 protein (log_2_FC = +1.56 at 0.5× IC_50_) was identified as a FecCD-family ABC transporter permease, a subfamily associated with nutrient and ion uptake in *S. aureus* (55). The Q2G2C3 protein (log_2_FC = +2.67 at 1× IC_50_), belonging to the YlaN (UPF0358) family, is an Fe(II)-binding protein that relieves Fur-mediated repression in *S. aureus*, thereby enabling release from repression of iron acquisition genes under iron-limited conditions(118). Its induction suggests that chloramphenicol stress may trigger secondary iron homeostasis responses. Similarly, Q2GW9 protein (log_2_FC = +2.22 at 2× IC_50_), containing a PSP1 C- terminal domain, was induced at high concentrations. Phage shock protein (PSP) systems are known to stabilize bacterial membranes and protect against envelope stress, which is consistent with the cell surface remodeling observed here (119).

Two AraC/XylS-type transcriptional regulators (Q2G0D2 and Q2FY65) were also up-regulated (log_2_FC = +1.73 and log_2_FC = +1.99 at 2× IC_50_, respectively), both showing high sequence identity to canonical AraC regulators. Members of this family, including Rbf and Rsp, are established regulators of biofilm formation in *S: aureus*, acting by modulating *icaR* expression and surface protein production (120). Their induction under chloramphenicol suggests a potential link between translation inhibition and biofilm-promoting regulatory circuits. By contrast, Q2FYE6 (log_2_FC = -1.74 at 2× IC_50_) and Q2G209 (log_2_FC = -2.55 at 0.5× IC_50_) proteins remain without functional annotation, while Q2G1G1 (Sce7726 family, log_2_FC = +1.80 at 2× IC_50_) also remains functionally unknown, though its induction at high chloramphenicol concentrations indicates stress-related activity.

Functional enrichment of PPI networks further contextualized chloramphenicol-induced proteomic responses (Figure S4). At 0.5× IC50, enriched terms were mainly related to DNA defense, restriction-modification systems, adenine methylation, general stress responses, and glutathione metabolism, indicating early activation of genome protection and redox homeostasis under translational stress. Arachidonic metabolism was also enriched, likely reflecting fatty acid/lipid remodeling rather than true arachidonic acid biosynthesis, which does not occur in staphylococci (121–124). Overall, this profile supports a defensive stress response rather than broad central metabolic reprogramming.

At 1× IC50, enrichment shifted toward metabolic reorganization, including fatty acid β-oxidation, fatty acid metabolism/degradation, and pathways involving XRE-family and BRCT-domain proteins, suggesting rewiring of energy and carbon metabolism.

Consistently, acyl-CoA dehydrogenase (Q2FVW7) was repressed, whereas enoyl-CoA hydratase (Q2G1C9) and dihydrolipoamide acetyltransferase (Q2FY54) were induced, supporting alternative energy-generating routes. Enrichment of amino acid and aromatic compound degradation pathways, together with induction of nutrient transporters (Q2FVZ8 and Q2FX87), further supports broader substrate utilization under stress (125, 126). Together, these changes suggest that MRSA compensates for translational inhibition by expanding metabolic inputs to sustain energy and nucleotide homeostasis.

At 2× IC50, enrichment was more restricted, dominated by nickel cation binding, with weaker persistence of DNA-restriction pathways and XRE-family/BRCT-domain proteins. This agrees with up-regulation of urease subunit γ (ureA, Q2FVW5), a Ni^2+^-dependent protein, and supports a role for metal cofactor homeostasis and urease-associated nitrogen metabolism in the high-dose response. This is consistent with the known roles of urease and metal homeostasis in *S. aureus* stress adaptation and persistence (127–129).

Proteins dysregulated in at least two conditions were also analyzed as a separate network (Figure 2). Although this set was not enriched on its own, expansion with first-shell interactors highlighted local network clusters linked to ribonucleoside-diphosphate reductase, BRCT-domain and XRE-family proteins, periplasmic transport systems and oxidoreductase/redox-associated enzymes, supporting conserved adaptive modules centered on DNA metabolism, transcriptional control, nutrient uptake, and redox balance under chloramphenicol stress (104, 130–135).

**Figure 2.**
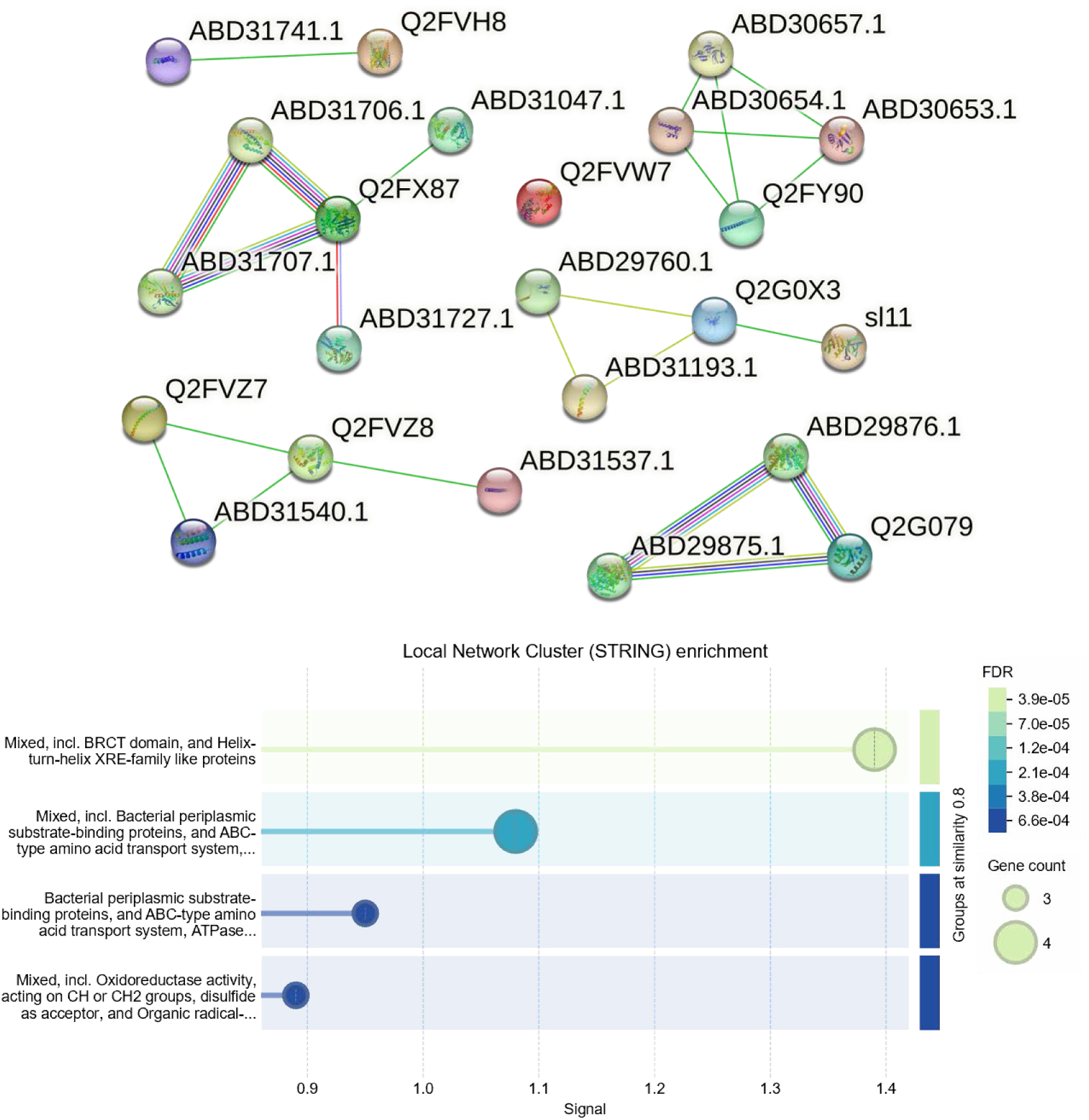
STRING analysis of proteins dysregulated in at least two chloramphenicol conditions. Upper panel: protein–protein interaction network generated from proteins significantly altered in at least two of the three chloramphenicol concentrations tested (0.5×, 1×, and 2× IC50), with first-shell interactors added for network contextualization. Lower panel: functional enrichment summary of the same network showing significantly enriched terms passing the threshold of FDR ≤ 0.05.

Comparison with a curated panel of MRSA virulence-, resistance-, and stress-related proteins showed that only the RNA polymerase sigma factor was recurrently dysregulated, being repressed at 0.5 and 2× IC_50_. This suggests attenuation of transcriptional stress-adaptation pathways, including functions associated with adhesins, biofilm formation, and methicillin resistance (136, 137).

#### 3.1.3 Ciprofloxacin

Ciprofloxacin is a fluoroquinolone that targets bacterial topoisomerase II (DNA gyrase) and topoisomerase IV enzymes, leading to DNA replication inhibition (138–140).

Mutations in the genes encoding these enzymes result in proteins with reduced susceptibility or insensitivity to fluoroquinolones (141). Moreover, active efflux of the drug by membrane transporters is also associated with resistance. Exposure to ciprofloxacin altered significantly protein expression of 39 proteins (Figure S5) across sub-inhibitory to inhibitory concentrations.

Dysregulated proteins could be broadly grouped into categories related to DNA maintenance and repair, transcriptional regulation, metabolism transport, envelope remodeling, prophage elements, and uncharacterized stress associated factors. The proteomic changes align with ciprofloxacin’s primary action of DNA gyrase and topoisomerase, which includes DNA double-strand breaks and triggers replication stress responses (138–140).

The replication initiation control protein YabA (Q2G2W8, log_2_FC = +2.24 at 1× IC_50_, Figure S5) was induced, possibly to restrain replication initiation under DNA damage. YabA is a negative regulator of the DnaA replication initiator (142, 143). In parallel, a site-specific adenine DNA methyltransferase (Q2FXD0, +1.71 at 0.5× IC_50_) was up-regulated, supporting the activation of restriction-modification defense systems. Beyond DNA protection, adenine methylation in bacteria can contribute broadly to pathogenicity, gene expression regulation, and adaptive plasticity, including the capacity of *S. aureus* to acquire antibiotic resistance and persist (144, 145). In contrast, the chromosome segregation protein FtsK (Q2FXI9, log_2_FC = - 4.03 at 2× IC_50_) was repressed. Ftsk, together with SpoIIIE, ensures faithful chromosome partitioning in *S. aureus*, and disruption of these systems leads to severe nucleoid segregation defects (146). Its down-regulation under ciprofloxacin stress therefore suggest that while MRSA activates protective mechanisms to stabilize DNA, the drug simultaneously compromises chromosome segregation machinery, intensifying replication stress.

Transcriptional reprogramming emerged as a major adaptive feature. The RNA polymerase sigma factor SigS (Q2FXF2, log_2_FC = +1.90 at 2× IC_50_) was induced, consistent with its established role in stress adaptation, virulence, and immune invasion in *S. aureus* (147). The core DNA-directed RNA polymerase subunit β’ (*rpoC,* Q2G0N5, log_2_FC = +1.73 at 2xIC_50_) was also up-regulated, in line with reports that *rpoB*/*rpoC* mutations are central to transcriptional adaptation and antibiotic resistance in MRSA (10). As shown in Figure S5, multiple HTH regulators were dysregulated, including gntR-type (Q2G1B1, log_2_FC = +4.17 and +3.07 at 0.5 and 2× IC_50_, respectively), marR-type (Q2G141, log_2_FC = +1.64 and +1.75 at 1× and 2× IC_50_, respectively), lacI-type (Q2G1A3, log_2_FC = +1.58 at 1× IC_50_), and SarU (Q2G1T7, log_2_FC = -2.08 and -1.57 at 0.5× and 1× IC_50_, respectively), alongside the persulfide-sensing repressor CstR (Q2G1R8, log_2_FC = +2.55 at 1× IC_50_). These transcriptional regulators collectively link DNA damage responses to redox regulation and virulence control (148–153).

Metabolic remodeling was also evident. The pentose phosphate pathway enzyme glucose-6-phoshpate 1-dehydrogenase (*zwf*, Q2FY66, log_2_FC = +1.52 at 1× IC_50_) was induced, indicating a shift toward NADPH generation of antioxidant defense (154). Aspartokinase (Q2FYP1, log_2_FC = -1.96, -1.53 and -2.25 at 0.5×, 1× and 2× IC_50_, respectively), a key enzyme in lysine biosynthesis, was down-regulated in all experimental conditions, consistent with previous reports of lysine pathway control in *S. aureus* (155). Additionally, the phosphoenolpyruvate-dependent phosphotransferase system (PTS) was represented by induction of HPr kinase (Q2G045, log_2_FC = +4.43 at 2× IC_50_) and ABC transporter permease Q2G0D7 (log_2_FC = +2.58 at 2× IC_50_), pointing to increased carbohydrate uptake and energy metabolism during ciprofloxacin stress (156). BLASTp analysis (72) further identified Q2G0D7 protein as VraG, an ABC transporter permease that is part of the GraSR two-component system involved in drug antimicrobial resistance (157).

Transport and envelope associated proteins showed variable regulation. The accessory Sec protein Asp1 (Q2FUW3, log_2_FC = +2.41 at 1× IC_50_) was altered, supporting its critical role in the export of the serine-rich adhesin SraP through the accessory Sec system which is related to virulence (158). Multiple ABC transporters (Q2G0D7 induced (log_2_FC = +2.58 at 2× IC_50_); Q2FW75 (log_2_FC = -2.52 at 0.5× IC_50_) and Q2FYW3 (log_2_FC = -2.13 and -2.00 at 0.5× and 2×IC_50_, respectively) repressed) were also modulated, consistent with their importance in antimicrobial resistance (104). The MFS protein (Q2G0C4, log_2_FC = +2.14 at 1 IC_50_) was up-regulated, which may support drug efflux or nutrient flux, in line with the reported role of MFS transporters in antibiotic resistance and stress adaptation (107, 108). The immunoglobulin-binding protein Sbi (Q2FVK5, log_2_FC = +2.96 at 0.5× IC_50_) was strongly induced, suggesting enhanced immune evasion and virulence (159). Moreover, the envelope regulator MsrR (Q7BHL7, log_2_FC = +1.68 at 1× IC_50_) was also induced, consistent with its role in maintaining cell wall integrity, mediating antibiotic resistance, and responding to cell wall-active stressors (160).

Prophage-associated proteins were also dysregulated, including a phage terminase large subunit (Q2FWT4, log_2_FC = -2.25, -1.94, and -2.99 at 0.5×, 1× and 2× IC_50_, respectively) and a phage structural protein (Q2FZ07, log_2_FC = +1.98 and +2.11 at 0.5× and 1× IC_50_, respectively). This points to prophage activation as part of the ciprofloxacin stress response, consistent with previous reports of fluoroquinolone-induced SOS response (117, 119, 149).

Five uncharacterized proteins (Q2FUZ5, Q2FVY8, Q2G018, Q2G215, and Q2G2V8) remain annotated as hypothetical proteins with no clear homologs, though their consistent induction or repression suggest ciprofloxacin stress-linked roles. Q2FXA9 (log_2_FC = -1.62, -2.03 and -1.68 at 0.5×, 1x and 2× IC_50_, respectively) showed strong BLASTp similarity to the DUF6978 family, predicted to adopt an α/β fold with conserved histidine residues forming a putative metal binding site, hinting at roles in metal ion regulation or redox processes (161). Q2G0C5 (log_2_FC = -1.63 at 1× IC_50_) aligned with a sugar efflux transporter, part of the DUF5080 family that has also been associated with the type VII secretion system (T7SS), including membrane proteins linked to toxin-immunity interactions (161, 162). Q2FX03 (log_2_FC = +3.35 at 2× IC_50_), annotated as an UPF0421 protein, matched the FUSC family (also called FusC-like proteins, whose members are typically membrane-associated and often linked to drug resistance. FUSC proteins are implicated in fusidic acid resistance, either by contributing to efflux activity or supporting *fus* gene function (163). Its strong up-regulation suggests possible involvement in membrane remodeling or metabolite/drug transport.

Two proteins were up-regulated in at least two concentrations, suggesting their role as conserved adaptive factors. The first was a phage-associated protein (Q2FZ07), which was strongly up-regulated at both 1× and 2× IC_50_. Phage proteins have been repeatedly observed to increase under antibiotic stress and are associated with prophage mobilization, horizontal gene transfer, and virulence modulation in *S. aureus* (117). Their induction under ciprofloxacin parallels observations from ampicillin and chloramphenicol, supporting the view that prophage activation is a generalized MRSA strategy to enhance genetic plasticity during stress.

The second recurring factor was a HTH gntR-type transcriptional regulator (Q2G1B1), which was up-regulated at both 1× and 2× IC_50_ conditions. GntR regulators form a large family of bacterial transcription factors that integrate environmental sensing with metabolic control. In *S. aureus*, GntR-like proteins have been linked to carbon metabolism, oxidative stress resistance, and virulence regulation (153). Its consistent induction under ciprofloxacin suggests a central role in reprogramming metabolism and gene expression to support survival during DNA damage stress.

At 0.5× IC50, STRING analysis (Figure S6) showed enrichment of DNA restriction-modification functions, consistent with detection of a site-specific adenine DNA methyltransferase (Q2FXD0), indicating activation of genome defense and DNA integrity mechanisms under fluoroquinolone stress. Because DNA methylation influences replication, mismatch repair, and transcriptional regulation, and has been linked to adaptive states in *S. aureus*, these data support early coordination of genome protection and regulatory adaptation. Additional enrichment of siderophore-mediated iron import and SarA/Rot-centered regulatory networks, together with phage-associated and stress-linked nodes in the interaction network, further suggests concurrent remodeling of metal homeostasis and transcriptional control during early ciprofloxacin adaptation (144, 145).

At 1× IC50, network enrichment shifted toward carbohydrate metabolism, particularly the phosphoenolpyruvate-dependent phosphotransferase system (PTS) and carbohydrate transport, indicating reorganization of carbon uptake and flux under ciprofloxacin stress. As PTS components integrate nutrient uptake with metabolic and transcriptional control, including catabolite and stress regulation (106, 156, 164), these results suggest that MRSA mobilizes carbohydrate import and PTS-linked regulatory functions to support adaptation. Phage portal proteins and HNH nuclease networks further indicate concurrent activation of phage-related and DNA-processing modules.

At 2× IC50, enrichment was more limited, with lysine biosynthesis reaching significance. This agrees with repression of aspartokinase and indicates modulation of amino acid biosynthesis under high ciprofloxacin stress. In the broader network context, this response occurred alongside additional stress-associated and regulatory clusters, suggesting that severe fluoroquinolone exposure is accompanied by selective metabolic narrowing rather than broad biosynthetic activation. Together, these data support reallocation of cellular resources away from anabolic growth-related functions and toward stress adaptation and survival (155)

Proteins dysregulated in at least two conditions (Figure 3) were also analyzed as a separate network, revealing clusters linked to HNH nucleases and phage-associated proteins, consistent with genome-plasticity and phage-related responses under ciprofloxacin stress (117, 147). Additional modules involved transport- and membrane-associated functions, including ABC transporter- and cardiolipin synthase-related clusters, suggesting membrane and transport remodeling (104, 157, 160, 165). Toxin/transposase-related clusters further supported involvement of mobile genetic element-associated functions (147, 153).

**Figure 3.**
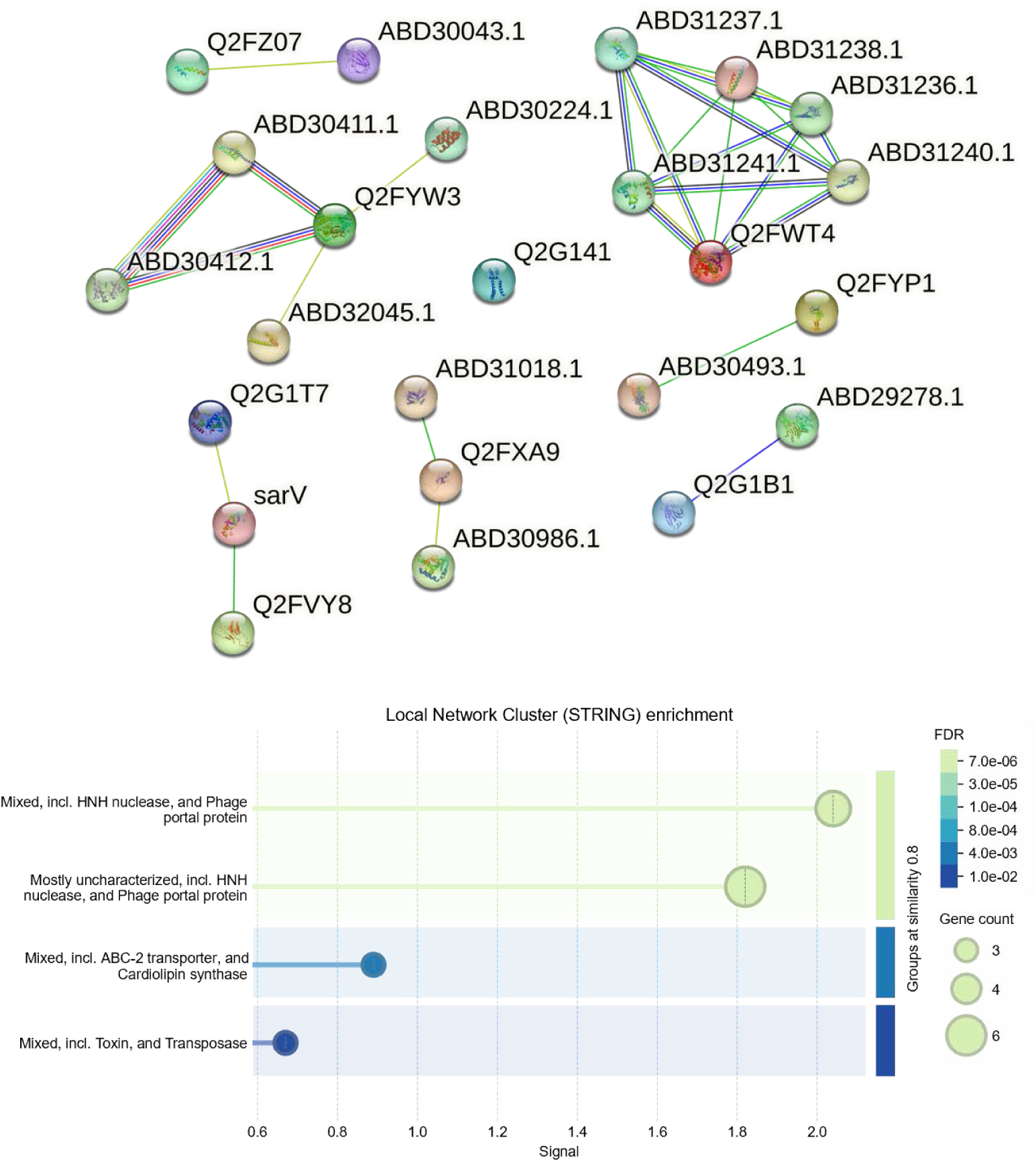
STRING analysis of proteins dysregulated in at least two ciprofloxacin conditions. Upper panel: protein–protein interaction network generated from proteins significantly altered in at least two of the three ciprofloxacin concentrations tested (0.5×, 1×, and 2× IC50), with first-shell interactors added for network contextualization. Lower panel: functional enrichment summary of the same network showing significantly enriched terms passing the threshold of FDR ≤ 0.05.

#### 3.1.4 Methicillin

Methicillin is a β-lactam antibiotic that inhibits PBPs, which are involved in peptidoglycan biosynthesis and contribute to cell wall stability (50). *S. aureus* was able to develop resistance to this drug by acquiring the *mecA* gene which codes for a different PBP, PBP2a. This enzyme can continue peptidoglycan crosslinking in the presence of methicillin, maintaining cell wall integrity (1, 166). Exposure to methicillin significantly altered the proteomic profile of MRSA, with 35 proteins found dysregulated across different concentrations (Figure S7).

Methicillin stress triggered broad remodeling of envelope- and metabolism-linked functions. The acyl-CoA dehydrogenase Q2FW7 was strongly repressed in all conditions (log_2_FC = -5.64, -4.90, -3.74 at 0.5×, 1× and 2× IC_50_, respectively), pointing to reduced fatty acid catabolism and altered energy flow (125). Induction of ABC transporter permease Q2FVL3 (log_2_FC = +2.17 and +1.97 at 1× and 0.5×IC_50_, respectively) and the PTS glucose transporter Q2FXJ8 (log_2_FC = +1.51 at 1× IC_50_) indicates that MRSA reallocates membrane transport capacity toward external nutrient salvage under methicillin stress. Amino acid import provides building blocks and nitrogen for peptidoglycan and teichoic-acids synthesis (103, 104), while PTS-mediated sugar uptake directly supplies phosphorylated hexoses for glycolysis and UDP-sugar pools required for cell wall glycan assembly (164, 167, 168). Together these changes are expected to sustain envelope repair and cellular energy/redox balance when de novo biosynthesis is constrained.

Multiple DNA repair proteins were induced under methicillin stress. DNA-3-methyladenine glycosylase I (Q2FXR7, log_2_FC = +1.73 at 0.5× IC_50_) participates in base excision repair, recognizing and excising alkylated bases from DNA (169). RecF (Q2G275, log_2_FC = +2.52 at 0.5× IC_50_) supports homologous recombination repair of single-strand gaps (170). At higher concentrations, ATP-dependent helicase/deoxyribonuclease subunit B (Q2FZ76), part of the AddAB complex essential for double-strand break repair, was strongly up-regulated (log_2_FC = +4.02, +3.98 at 1× and 2× IC_50_, respectively) (170, 171). These responses highlight activation of multiple repair pathways to counter methicillin induced DNA damage.

A gnt-R type transcriptional regulator (Q2FZ17, log_2_FC = +2.48 at 1× IC_50_) was strongly induced under methicillin stress, suggesting transcriptional reprogramming to modulate metabolism, stress responses, and potentially virulence. GntR-family regulators, as mentioned before, are widely recognized for integrating environmental sensing with metabolic control, and in *S. aureus*, they have been linked to carbon metabolism, oxidative stress resistance, and adaptive responses to antibiotic-induced DNA damage (153).

Methicillin also triggered induction of prophage-associated proteins, including the phage head protein Q2FX56 (log_2_FC = +5.16 at 2× IC_50_), consistent with mobilization of prophage elements at higher stress. This mirrors observations under ampicillin and chloramphenicol, where phage-related proteins were similarly up-regulated, suggesting that activation of prophage elements represents a general stress response that enhances genetic plasticity and potentially facilitates horizontal gene transfer (116, 117, 172). *S. aureus* pathogenicity islands (SaPIs) rely on such phage proteins for excision, replication, and packing into phage-like particles, highlighting their role in adaptive mobilization under stress (172).

In parallel, the T7SS ATPase EssC (log_2_FC = +2.07 at 0.5× IC_50_) was also up-regulated, implicating activation of secretion-linked stress adaptation. The T7SS is conserved in *S. aureus* and mediates the secretion of antibacterial toxins, which can act as protection (162). Up-regulation of EssC suggest that MRSA may modulate interbacterial interactions and toxin export under methicillin stress, potentially as part of a broader survival strategy that complements prophage activation.

Moreover, Q2FUX6 protein was up-regulated in all conditions (log_2_FC = +1.58, 1.60 and 1.72 at 0.5×, 1× and 2× IC_50_, respectively), which remains without defined function but may represent a stress-specific regulator or effector.

To refine the interpretation of methicillin responses, BLASTp/tBLASTn searches (72) were conducted. Four proteins previously annotated as hypothetical proteins or poorly defined were re-annotated. Q2FUX6 protein remains a hypothetical protein with no clear homologs, though its consistent induction suggests stress-responsive function.

Q2FVG1 aligned with an ABC transporter permease (ID = 98.8 %, COV = 100 %), supporting enhanced amino acid and nutrient uptake, consistent with a broader strategy of metabolic remodeling under stress. Q2G1N4 aligned with SirA, a staphyloferrin B ABC transporter substrate-binding protein, and was down-regulated (log_2_FC = -1.62 to - 1.71 across conditions), suggesting modulation of iron acquisition to balance resource allocation under methicillin pressure. SirA is part the SirABC transporter, which specifically imports iron complexed with staphylobactin, an endogenously produced siderophore. Loss or repression of SirA has been shown to alter iron uptake, increase resistance to iron-dependent stressors, and influence transcription of siderophore biosynthesis genes, highlighting its critical role in iron homeostasis and bacterial fitness under iron-limited conditions (173). Finally, Q2FY63 protein matched a LacI-family transcriptional regulator (ID = 97.4 %, COV = 100 %), indicating selective silencing of non-essential carbon metabolism pathways. Members of the LacI family predominantly act as repressors of sugar metabolism genes, responding to intracellular effectors via HTH DNA-binding motifs (150). In contrast, the gnt-R type transcriptional regulator (Q2FZ17, log_2_FC = +2.48 at 1× IC_50_) was induced, reflecting activation of adaptative metabolic programs. Together, these opposing regulatory trends highlight that MRSA selectively tunes transcriptional networks under methicillin stress, simultaneously promoting nutrient acquisition and energy generation while down-regulating non-essential carbon metabolism to optimize survival.

To place these responses in context, STRING PPI analysis was performed (Figure S8). At 0.5× IC50, enrichment highlighted siderophore-dependent iron uptake and iron homeostasis, indicating that disruption of metal acquisition is an early feature of methicillin stress (103, 104, 173). Beyond iron-related functions, the network also included clusters associated with transport- and surface-linked processes, including secretion-related nodes and additional regulatory or stress-associated proteins, suggesting that early methicillin adaptation couples metal homeostasis with broader remodeling of the cell interface and stress-response functions.

At 1× IC50, enrichment shifted toward siderophore-dependent iron import, transition metal ion transport, and metabolic pathways linked to fatty acid, aromatic compound, and branched-chain amino acid degradation, consistent with broader metabolic rewiring under β-lactam pressure (125, 129, 167, 168, 174). Networks involving phage tail and portal proteins were also identified, suggesting engagement of phage-associated functions and increased genome plasticity during methicillin stress.

At 2× IC50, enrichment was centered on urease-associated, nickel-binding, and ammonium/urea-related networks, consistent with induction of urease-linked functions and supporting a role for metal homeostasis and nitrogen metabolism in the high-dose methicillin response (128, 175). Phage tail- and portal protein-associated networks were also detected, indicating additional engagement of phage-related functions, while sugar transferase- and anti-repressor-associated networks were present with weaker signal.

Proteins dysregulated in at least two methicillin conditions were also analyzed as a separate STRING network (Figure 4). This network contained modules linked to siderophore-dependent iron import and iron homeostasis, amino acid/ABC transporters, and phage tail- and portal-associated proteins, reinforcing iron and metal management, transport remodeling, and phage-related functions as recurrent components of methicillin adaptation. Overall, these data suggest that methicillin responses combine early perturbation of iron homeostasis, intermediate metabolic and transport reprogramming, and high-dose urease-linked stress tolerance.

**Figure 4.**
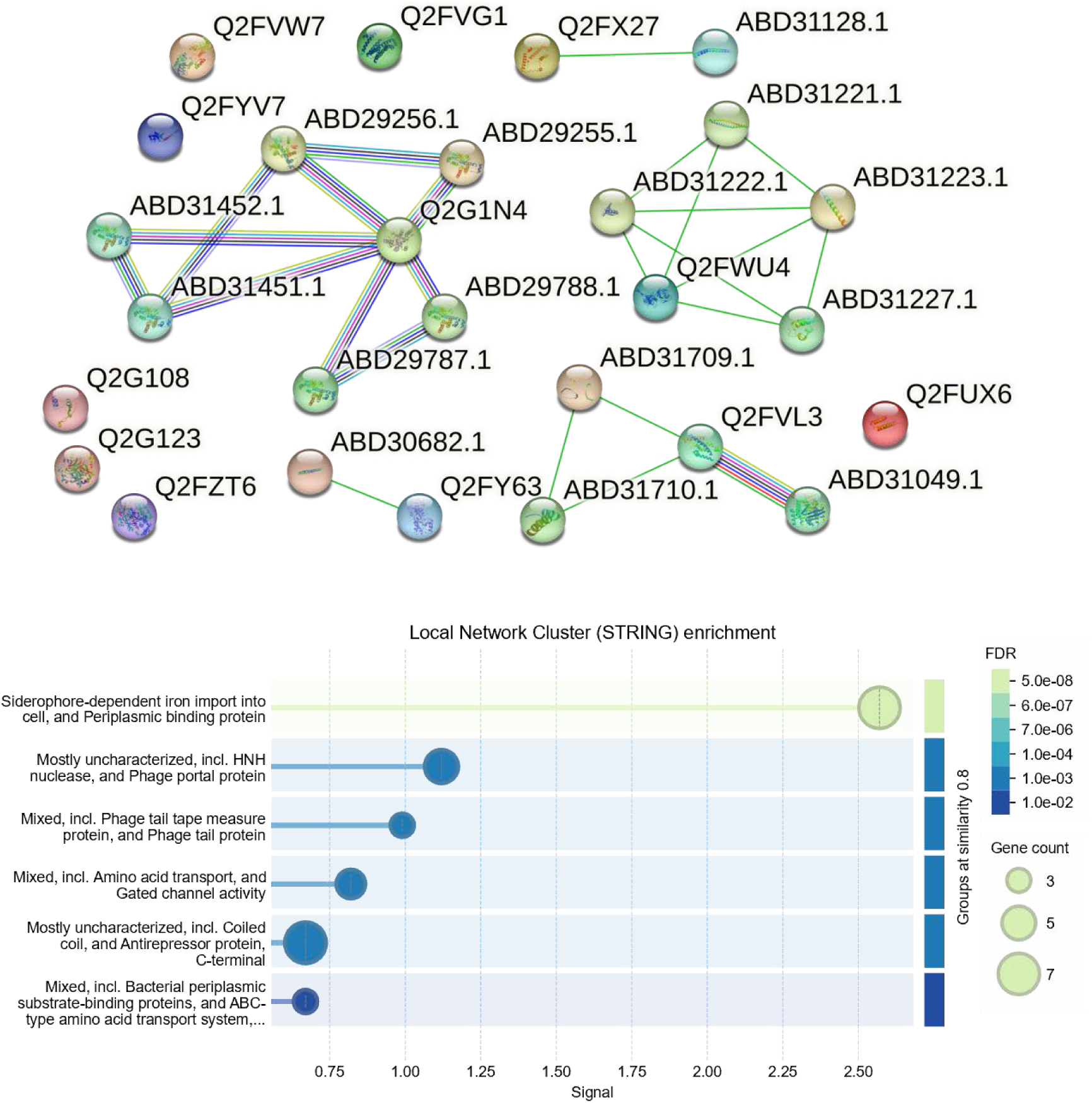
STRING analysis of proteins dysregulated in at least two methicillin conditions. Upper panel: protein–protein interaction network generated from proteins significantly altered in at least two of the three methicillin concentrations tested (0.5×, 1×, and 2× IC50), with first-shell interactors added for network contextualization. Lower panel: functional enrichment summary of the same network showing significantly enriched terms passing the threshold of FDR ≤ 0.05.

Cross-referencing with the curated MRSA panel identified recurrent dysregulation of three proteins. AgrB was down-regulated at all concentrations (log_2_FC = -2.23, -2.05 and -2.23 at 0.5×, 1× and 2× IC_50_, respectively), suggesting repression of *agr*-mediated (176) quorum sensing and virulence regulation. Sortase A was also consistently down-regulated (log_2_FC = -1.90, -1.84 and -1.81 at 0.5×, 1× and 2× IC_50_, respectively), indicating reduced surface display of adhesins and related virulence factors (177, 178). In contrast, the HTH-type regulator NorG was up-regulated at 0.5× and 1× IC_50_ (log_2_FC = +1.57 and +1.51 at 0.5× and 1× IC_50_, respectively), consistent with activation of resistance-associated regulatory pathways (153). Together, these changes suggest that methicillin stress suppresses quorum sensing and surface virulence functions while promoting adaptive resistance regulation.

#### 3.1.5 Vancomycin

Vancomycin is a glycopeptide antibiotic that binds to the D-Ala-D-Ala terminus of the peptidoglycan precursor, compromising the integrity of the bacterial cell wall (179). Vancomycin resistant *S. aureus* (VRSA) have already been isolated and this resistance is characterized mainly by modification of the terminal D-amino acid in Lipid II, which reduces vancomycin binding affinity (65, 179). Vancomycin exposure elicited a multifaceted adaptive response in MRSA, encompassing stress-responsive proteins, transport systems, cell wall and metabolic enzymes, and prophage-related elements. A total of 43 dysregulated proteins (Figure S9) were identified across the tested antibiotic concentrations, with distinct, dose-dependent shifts in the adaptive trajectory.

Consistent with vancomycin’s inhibition of peptidoglycan precursor utilization, the essential cell wall enzyme UDP-*N*-acetylmuramate-L-alanine-ligase (*murC*, Q2FXJ0) was strongly repressed at 1× IC_50_ (log_2_FC = -2.99). MurC catalyzes an early step in the peptidoglycan biosynthesis pathway (180). Functional studies have shown that staphylococcal MurC accepts L-Ala, L-Ser, or Gly as substrates, with a strong preference for L-Ala (181), and together with MurE and MurF, dictates the unique lysine-containing peptide cross-bridges that define *S. aureus* peptidoglycan. Despite being essential, Mur ligases remain underexploited as drug targets, with no clinical approved inhibitors for MurC (180).

Additional perturbations were observed in lysine biosynthesis and membrane lipid metabolism. Aspartokinase (Q2FYP1), a key enzyme in lysine pathway, was repressed (log_2_FC = -2.19 at 2× IC_50_), consistent with earlier observations that lysine biosynthesis is tightly controlled during β-lactam and glycopeptide exposure (155). Lysine is indispensable for *S. aureus* peptidoglycan crosslinking, and restriction of this pathway may represent a bottleneck in sustaining wall synthesis under vancomycin pressure.

Conversely, glycerol-3-phosphate acyltransferase PlsY (Q2FYS6) was induced (log_2_FC = +1.72 at 2× IC_50_). PlsY catalyzes the first step in membrane phospholipid biosynthesis (182), thereby supporting envelope maintenance when peptidoglycan synthesis is compromised. Together, these adjustments indicate a reorientation of envelope metabolism under vancomycin pressure.

The chaperonin GroEL (Q2FWN4) was observed up-regulated at 0.5× and 1× IC_50_ (log_2_FC = +3.06 and +1.89, respectively), consistent with its role in protein folding under stress. GroEL induction is a hallmark of antibiotic-triggered proteotoxic stress, as the GroEK/ES complex safeguards nascent and misfolded proteins during stress.

Previous studies have highlighted GroEL/ES as indispensable for bacterial viability and increasingly attractive as a therapeutic target, with several small molecule inhibitors already demonstrating antibacterial activity. Importantly, despite its moderate evolutionary conservation with human mitochondrial HSP60, is still a viable bacterial-selective drug target (183, 184).

Similarly, peptide deformylase (Q2FZG6) was induced at 2× IC_50_ (log_2_FC = +2.85), pointing to enhanced protein maturation and translational stress adaptation. Peptide deformylase is essential in Gram-positive bacteria, catalyzing the removal of *N*-formyl groups from nascent polypeptides as a prerequisite for normal protein function (185). In *S. aureus*, two homologs (DefA and DefB) have been described, with *defB* proven essential for viability (186). Induction of deformylase under vancomycin stress therefore reflects increased pressure on protein quality control systems, reinforcing the tight link between translation-associated pathways and antibiotic resistance. Together with GroEL up-regulation, these data suggest that vancomycin tolerance is supported not only by envelope remodeling but also by intensified protein folding and maturation machinery.

Some proteins linked to DNA metabolism were also modulated. DNA polymerase β (Q2FZD4) was induced at 1× IC_50_ (log_2_FC = +2.89), and DNA-directed DNA polymerase (Q2G0T5) at 0.5× IC_50_ (log_2_FC = +1.60), suggesting increased DNA repair activity (170). Conversely, the GTPase Der (Q2FYG0, log_2_FC = -3.49 at 2× IC_50_), essential for ribosome assembly and stability, was repressed, which may compromise translation under vancomycin stress (187).

The TelA-like protein (Q2FYM7) was observed up-regulated at 0.5× IC_50_ (log_2_FC = +3.39), suggesting a role in vancomycin adaptation. TelA homologs were originally linked to resistance against toxic anions such as tellurite, selenite, and arsenite (188). More recently, TelA was shown to contribute to resistance against multiple envelope-targeting antimicrobials, including nisin, bacitracin, and β-lactams, in *Listeria monocytogenes* (189). Its induction under vancomycin stress is therefore likely part of a general envelope protective response.

Several metabolic and envelope-associated enzymes were differentially expressed. Mannitol-1-phosphatase dehydrogenase (Q2FW96) was induced at 0.5× and 2× IC_50_ (log_2_FC = +1.79 and +2.67, respectively). This enzyme is central to mannitol catabolism in *S. aureus*. Beyond its metabolic role, mannitol metabolism has been linked to survival against membrane-active fatty acids and oxidative stress (190), and in other bacteria it functions as an osmoprotectant under high salt stress (191). Its induction under vancomycin suggests a role in redox buffering and osmotic protection during envelope damage. A α/β hydrolase (Q2FYZ3, log_2_FC = +2.59 at 1× IC_50_) was also up-regulated. Members of this family include peptidoglycan hydrolases, a large family of enzymes essential for cell wall turnover and division in *S. aureus* (192, 193). Their modulation under antibiotic stress likely supports compensatory remodeling of the cell wall, a critical strategy to withstand vancomycin-induced envelope disruption.

Vancomycin exposure also triggered broad changes in transport systems. At 2× IC_50_, a cation/H^+^ exchanger (Q2FVI0, log_2_FC = +1.95) was up-regulated. *S. aureus* encodes multiple Na^+^/H^+^exchangers of the Mnh type, which maintain cytoplasmic pH and ion homeostasis under alkaline and high-salt conditions, and deletion of *mnhA1* in particular reduces both growth at elevated pH and virulence *in vivo* (194, 195). Their induction under vancomycin likely reflects a protective response to envelope stress, which perturbs ionic balance. Efflux-related systems were also induced, including an ABC transporter ATP-binding protein (Q2FVJ2, log_2_FC = +1.51) and MFS transporter (Q2G2B3, log_2_FC = +3.54). ABC transporters in these bacteria not only contribute to resistance and solute uptake but are also emerging as potential antistaphylococcal drug targets (104). The MFS family includes multidrug efflux pumps such as NorA, which export structurally diverse antibiotics and are linked to multidrug resistance. Inhibitors of these pumps are being actively explored to restore susceptibility (107, 196).

Metal transport systems also shifted. The putative hemin transport permease HrtB (Q2G168) was induced at 1× and 2× IC_50_ (log_2_FC = +2.64 and +1.90, respectively), consistent with the need to balance iron acquisition with protection against heme toxicity. The HrtAB system not only limits heme-mediated damage but also modulates host immune responses by regulating the expression of neutrophil-interfering proteins (197, 198). In contrast, nickel-binding protein NikA (Q2G2P5) was repressed (log_2_FC = -2.25 at 2× IC_50_). The *S. aureus* Nik system provides nickel for urease, which neutralizes acidic environment (199). Its repression may reflect trade-offs in metal homeostasis under vancomycin stress. Finally, the regulatory protein YycH (Q2G2U3) was observed down-regulated in all conditions (log_2_FC = -1.73, -1.85 and -1.64 at 0.5×, 1×, and 2× IC_50_, respectively). YycH, together with Yycl, positively regulates the essential WalRK two component system, stimulating WalK phosphorylation and thereby driving cell wall hydrolase expression (200, 201). Loss or mutation of YycH is associated with vancomycin-intermediate *S. aureus*. Its observed repression at all conditions may represent an adaptive shift toward cell wall remodeling to tolerate vancomycin. Collectively, these transporter changes reflect broad reprogramming of ion, metal and drug transport pathways under vancomycin stress.

Transcriptional reprogramming was also observed. The Rrf2 family transcriptional regulator (Q2G0I3) was induced (log_2_FC = +2.51 at 0.5× IC_50_), consistent with the role of Rrf2 regulators as redox sensors and metabolic switches in *S. aureus* (202). In parallel, an AraC/XylS-type regulator (Q2FY65, log_2_FC =+2.90 at 2× IC_50_) was up-regulated. Members of this family, such as Rbf and Rsp, are established regulators of biofilm formation and virulence gene expression in *S. aureus* (120, 149). These findings suggest that vancomycin stress activates transcriptional programs tied to oxidative defense and biofilm-associated adaptation. By contrast, a prophage-associated protein (Q2FWS9) was observed down-regulated (log_2_FC = -1.68 and -1.57 at 0.5× and 1× IC_50_, respectively), contrasting with the strong prophage mobilization observed under methicillin or ciprofloxacin, suggesting that vancomycin stress does not strongly activate prophage elements.

Several hypothetical proteins were refined through BLASTp/tBLASTn (72). Q2FVC6, annotated as cytosolic protein, aligned with an insertion sequence ATP-binding protein (ID = 83.3 %, COV = 100 %), suggesting a potential link to mobile element activity and genomic plasticity under stress (203). Q2FVH0 aligned with an osmoprotectant ABC transporter substrate-binding protein (ID = 98.4 %, COV = 100 %). Given that osmoprotectant ABC transporters are inducible by osmotic stress and controlled by stress-regulated genes, its induction under vancomycin is consistent with envelope/osmotic adaptation (204). By contrast, Q2G0R7 matched a polysaccharide biosynthesis protein (ID = 99.2 %, COV = 100 %) but was consistently repressed (log_2_FC = -1.90, -2.19 and -2.66 at 0.5×, 1× and 2× IC_50_, respectively). This protein can contribute to modifications of the cell envelope of biofilm-associated tolerance (79, 80). Several proteins (Q2FUX6, Q2FVD3, Q2FWS9, Q2G018, Q2G0Z3, Q2G1Q7 and Q2G209) remained annotated as hypothetical, though their dysregulation indicates potential antibiotic stress-linked functions.

To place these responses in context, STRING PPI analysis was performed (Figure S10). At 0.5× IC50, enrichment was mainly related to translation, gene expression, and biosynthetic functions, consistent with induction of GroEL and ribosomal protein bL32 and repression of proteins linked to aminoacyl-tRNA metabolism and RNA handling (184, 205–207). n parallel, the network also suggested an additional envelope-associated module, indicating that early vancomycin stress affects both information-processing functions and cell-surface remodeling.

At 1× IC50, enrichment was detected only after expansion with 65 first-shell interactors and highlighted lactose metabolic processes. Induction of hydrolase Q2FYZ3 (log_2_FC = +2.59), together with increased abundance of transporters including the ABC transporter Q2FVJ2 (log_2_FC = +1.51) and the MFS transporter Q2G2B3 (log_2_FC = +3.54), supports coordinated enhancement of carbohydrate metabolism and sugar uptake, consistent with STRING indications of altered sugar utilization, including glucosidase-, carbohydrate metabolism-, and phosphotransferase system-related functions (190, 191). Additional enrichment of cell death/CidB-LrgB family and Phe-tRNA-binding networks suggests that this shift in substrate utilization was accompanied by modulation of cell death-related functions and translational control.

At 2× IC50, significant enrichment was detected only after expansion with first-shell interactors. Enriched biological processes included nickel cation transport, metal ion transport, cation transport, peptide transport, and transmembrane transport. Induced proteins included HrtB (log_2_FC = +1.90), the cation/H^+^ exchanger Q2FVI0 (log_2_FC = +1.95), and additional ABC/MFS transporters, highlighting the centrality of metal homeostasis and transport remodeling in the vancomycin envelope-stress response (107, 195, 197–199).Proteins dysregulated in at least two conditions also formed a separate network that, after expansion, was enriched for PTS, and carbohydrate metabolism, supporting conserved carbohydrate-centered metabolic remodeling across vancomycin conditions (Figure 5) (190, 191, 208). Overall, vancomycin adaptation appears to involve early translational and proteostasis responses, intermediate carbohydrate remodeling, and high-dose reprogramming of transport, ion homeostasis, and signaling.

**Figure 5.**
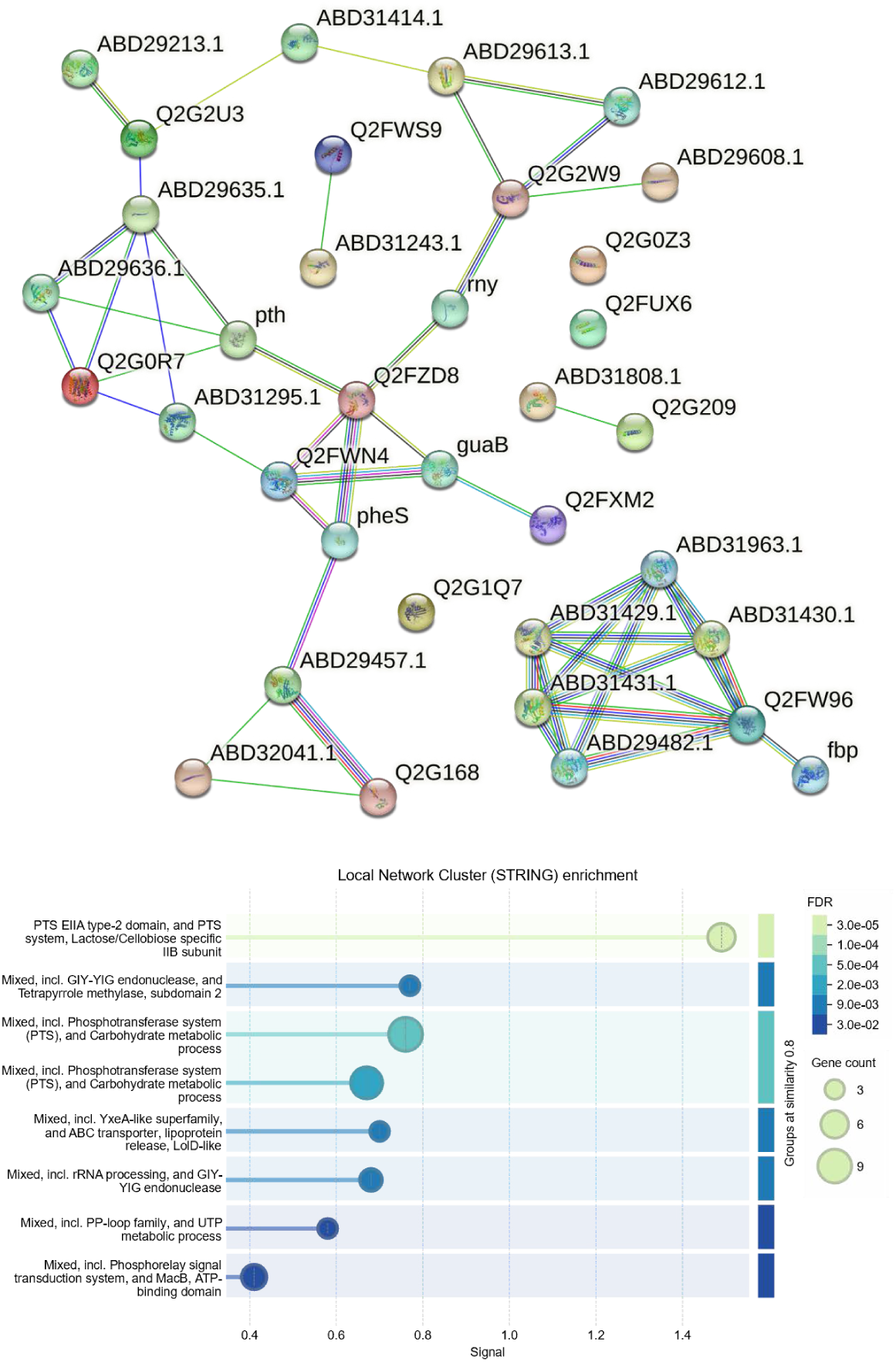
STRING analysis of proteins dysregulated in at least two vancomycin conditions. Upper panel: protein–protein interaction network generated from proteins significantly altered in at least two of the three vancomycin concentrations tested (0.5×, 1×, and 2× IC50), with first-shell interactors added for network contextualization. Lower panel: functional enrichment summary of the same network showing significantly enriched terms passing the threshold of FDR ≤ 0.05.

Cross-referencing with the curated MRSA panel identified recurrent dysregulation of several virulence- and stress-related proteins. AgrB was consistently repressed (log_2_FC = -1.93, -2.31 and -2.14 at 0.5×, 1× and 2× IC_50_, respectively), indicating down-regulation of *agr*-dependent quorum sensing (176). Similarly, the HTH-type transcriptional regulator SarU Q2G1T7 (log_2_FC = -1.69, -1.50 and -1.59 at 0.5×, 1× and 2× IC_50_, respectively), and RNA polymerase sigma factor SigS Q2G0P4 (log_2_FC = -2.31 and -2.06 at 0.5× and 2× IC_50_, respectively), were down-regulated, supporting broader repression of virulence-associated and stress-responsive transcriptional programs (137, 152).

Additional decreases were observed for the staphylococcal complement inhibitor Q2FWV6 (log_2_FC = - 1.89 and -2.00 at 1× and 2× IC_50_, respectively) and the serine protease SplF (log_2_FC = -1.95, -2.39 and - 2.47 at 0.5×, 1× and 2× IC_50_, respectively), suggesting reduced immune-evasion and extracellular virulence functions (209, 210). In contrast, the up-regulation of the global transcriptional regulator CodY Q2FZ27 (log_2_FC = +1.69, +1.50 and +1.63 at 0.5×, 1× and 2× IC_50_, respectively) indicates adaptive stress-response pathway. CodY is a regulator that can repress *agr* under nutrient-rich conditions but can also directly control *agr*-regulated genes in an *agr*-independent background (211). This function suggests that under vancomycin stress, CodY may not only reduce quorum sensing signaling but also substitute for *agr* in regulating key virulence determinants (212, 213). Collectively, these data suggest that vancomycin triggers a coordinated adjustment of quorum sensing, virulence, and resistance determinants, reflecting both conserved and adaptive pathways in MRSA under glycopeptide stress.

Proteomic profiling revealed that MRSA responds to antibiotic stress through a combination of shared and drug-specific adaptations. Across treatments, recurrently affected functions included cell envelope remodeling, transport, metabolic reprogramming, DNA maintenance, and stress-associated regulation. At the same time, each antibiotic imposed a distinct proteomic signature, with ciprofloxacin more strongly associated with DNA-centered responses, chloramphenicol with translational and metabolic adaptation, and the cell wall-active agents with envelope remodeling and transport-associated changes. Together, these findings indicate that MRSA proteomic adaptation is structured around a conserved stress-response core overlaid by antibiotic-specific remodeling patterns, providing a basis for subsequent integrative prioritization of candidate vulnerabilities.

### 3.2 Post-translational modifications of interest

While differential protein abundance provides critical insight into MRSA adaptation to antibiotic stress, it does not capture the full spectrum of protein regulation. Post-translational modifications (PTMs) provide an additional, rapid mechanism by which *S. aureus* adapts to environmental stressors such as antibiotic exposure. PTMs involve covalent modification of specific amino acid residues, which can modulate protein activity, stability, interactions, or subcellular localization without requiring new protein synthesis. Such modifications include acetylation, glycosylation, oxidation, phosphorylation, and succinylation, many of which have been shown to affect bacterial virulence, biofilm formation, immune evasion, and antibiotic resistance (214). Although once thought to be largely restricted to eukaryotic systems, it is now clear that PTMs are widespread in bacteria and central to the regulation of metabolic and stress-response pathways (215). PTMs enable bacteria to fine-tune metabolic fluxes and regulatory networks on short timescales, supporting adaptation under fluctuating environmental and host-associated conditions.

PTMs identified in the proteomic dataset are examined to highlight modifications that may contribute to MRSA adaptation under antibiotic stress. Particular attention is given to recurrent, statistically significant PTM changes detected for ampicillin, ciprofloxacin, and vancomycin (Figures S11–S13), whereas methicillin and chloramphenicol did not induce detectable PTM dysregulation under the tested conditions.

Genome maintenance and DNA damage control represented a shared PTM-sensitive axis with antibiotic-specific signatures. Under ampicillin, acetylation increased at Lys570 of the 3′–5′ exonuclease DinG (dinG, Q2FYH5) at 1× and 2× IC_50_ (log_2_FC = +1.55 and +1.94; Figure S11), consistent with PTM-level tuning of nuclease function during stress (216–218). Under ciprofloxacin, acetylation increased at Lys66 of a nuclease/exonuclease family protein (Q2FY88) at 0.5× and 1× IC_50_ (log_2_FC = +1.91 and +1.67; Figure S12), aligning with genotoxic stress adaptation (138–140, 216, 217) In contrast, phosphorylation of UvrABC system protein A (uvrA, Q2G046) decreased at His14 at 1× and 2× IC_50_ (log_2_FC = −2.46 and −2.04; Figure S12), indicating selective PTM remodeling within DNA repair pathways (219, 220).

Envelope and cell-wall-associated functions were most prominently remodeled under vancomycin. Protein Q2G0R7 (BLAST-supported polysaccharide biosynthesis protein; COV = 100 %, ID = 99.21 %) showed reduced acetylation at Lys212 across concentrations (log_2_FC = −2.15 to −2.85; Figure S13), reduced oxidation at Met206 (log_2_FC = −2.29 to −2.84; Figure S13), and reduced succinylation (log_2_FC = −2.04 and −2.29 at 0.5× and 2× IC_50_; Figure S13), consistent with PTM control of envelope matrix remodeling during glycopeptide stress. Deamidation increased at Asn299 of an acetyltransferase-3-domain-containing protein (Q2FZS9) at 1× and 2× IC_50_ (log_2_FC = +3.29 and +3.36; Figure S13), consistent with modulation of peptidoglycan O-acetylation-associated machinery (221, 222). Ciprofloxacin also affected envelope remodeling via LytN (*lytN*, Q9ZNI1), a protein involved in septal remodeling and cell separation (223, 224) that displayed altered succinylation at Lys90 (log_2_FC = − 1.74 and −2.26 at 1× and 2xIC_50_) and Lys97 (log_2_FC = 1.98 and 2.29; Figure S12).

Transport and nutrient acquisition were repeatedly targeted by PTMs. Under ampicillin, succinylation increased at Lys11 of an oligopeptide ABC transporter ATP-binding protein (Q2FZR0) at 0.5× and 1× IC_50_ (log_2_FC = +1.54 and +1.74; Figure S11) and at Lys10 of a predicted membrane protein (Q2FXB8) at 0.5× and 1× IC_50_ (log_2_FC = +3.09 and +2.63; Figure S11), with Q2FXB8 homologous to a MutE/EpiE-family permease (COV = 100 %, ID = 96.05 %) (225, 226). Under ciprofloxacin, an ABC transporter domain-containing protein (Q2G196) showed coordinated down-phosphorylation at T_215_YTYY_219_ across all concentrations (log_2_FC = −1.96 to −2.43; Figure S12), consistent with phosphorylation-linked regulation of ABC transport and resistance phenotypes (104, 227). Ciprofloxacin also altered a lactose-specific PTS component (Q2G2D4), with deamidation at Asn310 down-regulated at 0.5× and 2× IC_50_ (log_2_FC = −2.27 and −3.51; Figure S12) (228–230). Under vancomycin, deamidation increased at Asn200 of EcfT (Q2FZI4) at 1× and 2× IC_50_ (log_2_FC = +3.45 and +3.74; Figure S13), while deamidation decreased at Asn251 in Q2FVG1 (ABC permease; ID = 98.8 %, COV = 100 %) (log_2_FC = −2.56 to −3.13 at 0.5× to 2× IC_50_; Figure S13). High-dose vancomycin was further associated with induction of transport/ion-homeostasis proteins including HrtB (log_2_FC = +1.90) and a cation/H^+^ exchanger Q2FVI0 (log_2_FC = +1.95) (Figure S10).

Metabolic and redox/cofactor control was most apparent under vancomycin through broad dephosphorylation and coordinated succinylation loss. NnrD (Q2G2P8) showed loss of phosphorylation at Thr6/Thr9/Ser12 at 0.5× and 2× IC_50_ (log_2_FC = −4.59 and −5.35; Figure S13), and LacC (P0A0B9) was dephosphorylated at His232/Thr235/His234/Tyr237 (log_2_FC = −2.84 to −4.12; Figure S13). Succinylation was reduced in Q2FVG1, Q2FVG5, Q2G0R7, and Q2FXD3 (Figure S13), including Q2FVG5 (Fra/YdhG-like frataxin homolog) (log_2_FC = −2.18 and −2.43 at 0.5× and 2× IC_50_). Under ampicillin, phosphorylation increased at Ser3 of a MOSC-domain protein (Q2FVS9) at 0.5× and 2× IC_50_ (log_2_FC = +1.58 and +1.81; Figure S11), consistent with PTM-level modulation of sulfur/cofactor-linked functions.

Virulence- and mobile-element–linked proteins also carried PTM signals. Under ciprofloxacin, acetylation increased at Lys95 of Ssl7 (ssl7, Q2G2Y0) at 0.5× and 1× IC_50_ (log_2_FC = +2.25 and +2.81; Figure S12), and phosphorylation at Tyr94 was also increased (log_2_FC = +2.37 and +2.94; Figure S12). Under vancomycin, acetylation decreased in phage protein Q2FWQ8 at 1× and 2× IC_50_ (log_2_FC = −2.14 and −2.01; Figure S13), and succinylation decreased in DUF1433 protein Q2FXD3 (log_2_FC = −1.68 and −1.60 at 0.5× and 2× IC_50_; Figure S13), consistent with PTM involvement in mobile-element biology under stress.

Overall, PTM remodeling was antibiotic-specific but converged on genome maintenance, envelope remodeling, transport capacity, and metabolic/redox regulation. Ampicillin induced a limited set of recurrent PTMs centered on nuclease control, cofactor-linked regulation, and peptide transport (Figure S11). Ciprofloxacin produced structured transporter dephosphorylation with accompanying DNA repair and host-interface PTMs (Figure S12). Vancomycin showed the broadest multi-layer PTM remodeling across envelope-, transport-, and redox-linked proteins (Figures S10, S13), including repeated multi-PTM targeting of Q2G0R7, highlighting candidate regulatory nodes relevant to glycopeptide tolerance.

### 3.3 Metabolomics: Insights into metabolic adaptations

Metabolic profiling was performed to provide a direct view of the biochemical reprogramming of MRSA under antibiotic stress. Metabolites represent the final downstream products of gene and protein regulation and therefore can offer insight into rapid and adaptive changes in bacterial physiology that are not always captured using other omics approaches (25, 28, 29, 231, 232).

#### 3.3.1 Ampicillin

Untargeted metabolomics indicated a clear dose-dependent response to ampicillin in MRSA, with distinct pathway enrichments emerging as drug pressure increased (Figure S14). Across all three concentrations, guanylyl molybdenum cofactor (MoCo) biosynthesis was consistently enriched, supporting a recurrent perturbation of molybdoenzyme-linked redox and respiratory functions and suggesting that ampicillin impacts redox balance even at sub-inhibitory exposure (233, 234). In parallel, nucleotide metabolism was remodeled, with enrichment of purine salvage pathways (guanine/guanosine salvage at 0.5× IC_50_ and adenine/adenosine salvage at 2× IC_50_), consistent with maintaining nucleotide pools via lower-cost salvage routes during stress (235).

At 1× IC_50_, enrichment shifted toward cell-envelope homeostasis, including anhydromuropeptide recycling, heptaprenyl diphosphate biosynthesis, and the mevalonate pathway, which collectively support peptidoglycan fragment turnover and lipid-linked precursors required for cell wall integrity (236–238). At 2× IC_50_, perturbation extended to folate transformations and de novo pyrimidine biosynthesis, indicating broader disruption of biosynthetic capacity and processes required for growth under severe β-lactam stress (239, 240). Overall, ampicillin exposure was associated with early redox and nucleotide remodeling, followed by dose-escalation into envelope-associated pathway disruption and, at the highest concentration, impairment of nucleotide biosynthetic support for replication. Consistent with pathway-level enrichment, the MoO_2_-molybdopterin cofactor accumulated strongly and progressively across all ampicillin concentrations, with log_2_FC = +4.53, +6.09, and +6.80 at 0.5×, 1×, and 2× IC_50_, respectively (Figure 6A). This marked increase supports sustained engagement of MoCo metabolism during ampicillin stress and may reflect increased pathway flux and/or altered utilization of downstream molybdenum cofactors, including steps mediated by MoeA and MogA that catalyze molybdate insertion during cofactor biosynthesis (233).

**Figure 6.**
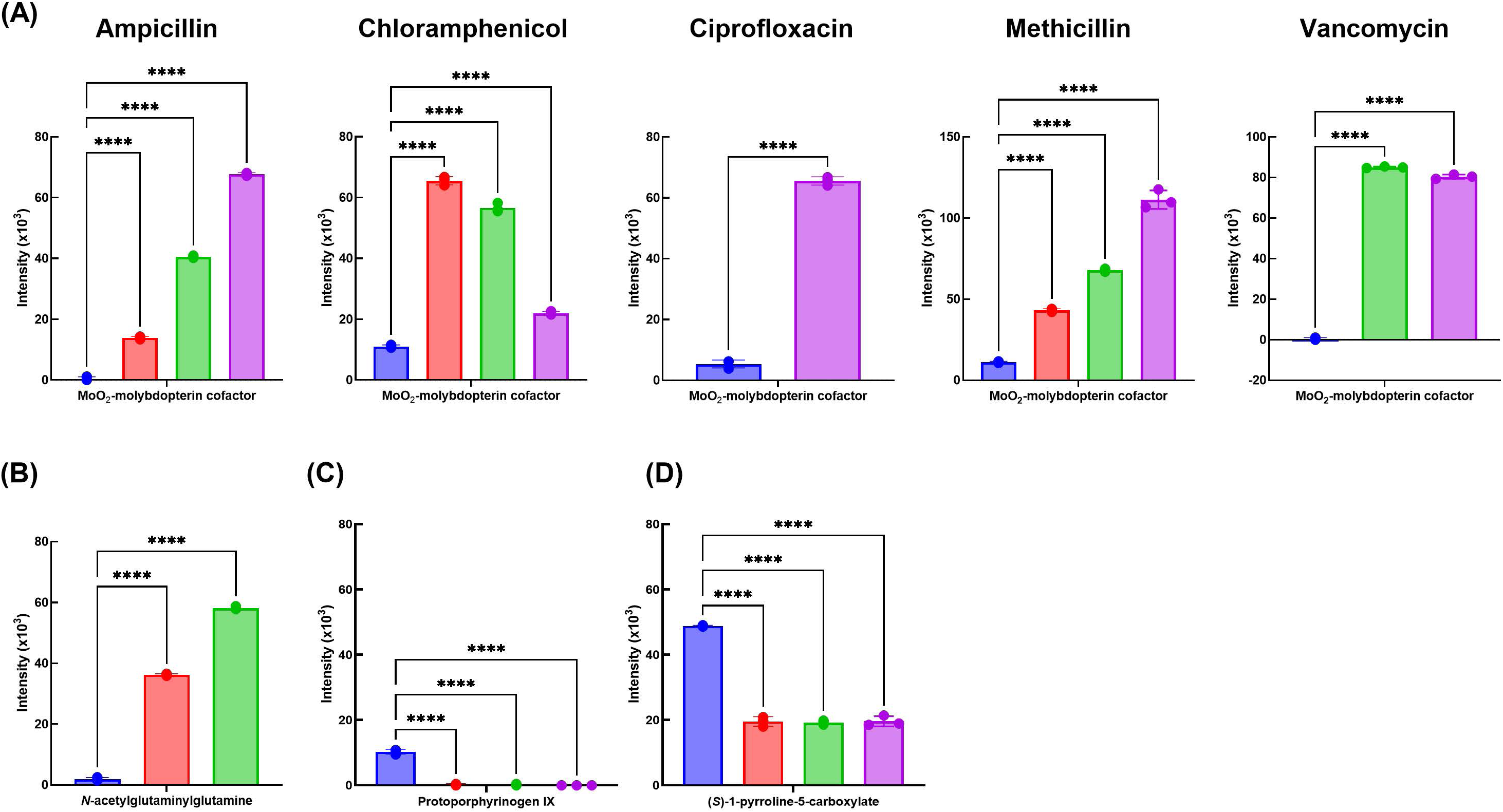
Relative intensities of selected metabolites across antibiotic treatments and concentrations. A: Mo_O2_-molybdopterin cofactor in control and antibiotic-treated samples for ampicillin, chloramphenicol, ciprofloxacin, methicillin, and vancomycin. B: *N*-acetylglutaminylglutamine under ampicillin treatment. C: Protoporphyrinogen IX under chloramphenicol treatment. D: (*S*)-1-pyrroline-5-carboxylate under ciprofloxacin treatment. Bars represent mean intensity, and dots represent technical replicate measurements. Colors indicate conditions: control (CTR, blue), 0.5× IC_50_ (red), 1× IC_50_ (green), and 2× IC_50_ (purple). Statistical comparisons were performed against the corresponding control using two-sided Welch’s *t*-tests. ****, *p* < 0.0001.

Collectively, these findings highlight the MoO_2_-molybdopeterin cofactor as a robust biomarker of adaptive responses to ampicillin, and an interesting therapeutic target to be explored. As molybdenum enzymes have been implicated in supporting bacterial survival under oxidative and antibiotic stress (234), targeting *moeA* or other enzymes of this pathway could represent a viable strategy to potentiate β-lactam efficacy against MRSA.

#### 3.3.2 Chloramphenicol

Chloramphenicol exposure induced a concentration-dependent metabolic response in MRSA, with pathway enrichment indicating early redox/nucleotide adjustments that broadened toward carbon and lipid remodeling as drug pressure increased (Figure S15). At 0.5× IC_50_, enriched pathways included *N*-acetylglutaminylglutamine (NAcGlnGln) amide biosynthesis, CMP phosphorylation, and guanylyl MoCo biosynthesis, consistent with rapid stress adaptation involving redox-associated metabolism and nucleotide/energy handling under translational inhibition (241, 242). The concomitant enrichment of teichoic acid (poly-glycerol) biosynthesis, cardiolipin biosynthesis, and related envelope-associated routes suggests early membrane/cell wall remodeling responses (165, 243, 244).

At 1× IC_50_, guanylyl MoCo biosynthesis remained enriched and additional changes appeared in alternative carbon utilization (e.g., lactose/galactose degradation I, androstenedione degradation) together with anhydromuropeptide recycling, indicating continued redox engagement alongside adjustments in carbon sourcing and cell wall turnover during sustained translation stress. At 2× IC_50_, the response became more focused, with persistent enrichment of MoCo-related pathways and additional perturbation of palmitate biosynthesis II, consistent with altered fatty acid metabolism and membrane lipid composition under high antibiotic stress (245–247). Collectively, these results support a trajectory in which chloramphenicol initially perturbs redox and nucleotide metabolism with modest envelope remodeling, then extends toward carbon and lipid pathway reconfiguration as exposure increases.

Consistent with the pathway enrichment, the MoO_2_-molybdopterin cofactor was found up-regulated across all concentrations (Figure 6A), with log_2_FC values of +0.98, +2.35, and +2.57 at 0.5×, 1×, and 2× IC_50_, respectively, supporting repeated engagement of molybdenum cofactor metabolism during antibiotic stress. In addition, NAcGlnGln, a conserved bacterial stress/osmoprotection-associated (248), was strongly increased at 0.5× IC_50_ (log_2_FC = +7.24). Its induction during chloramphenicol exposure suggests that translational inhibition may activate stress response pathways similar to those triggered by osmotic or redox imbalance. Given the role of NAcGlnGln in stress adaptation, disrupting its synthesis might sensitize bacteria to antibiotic induced stress. However, further work is needed to confirm this possibility as a viable target in *S. aureus*.

#### 3.3.3 Ciprofloxacin

Ciprofloxacin exposure produced a dose-dependent metabolomic response consistent with its genotoxic mechanism, with early enrichment dominated by nucleotide salvage/turnover and progressive expansion toward broader metabolic rewiring at higher drug pressure (Figure S16). At 0.5× IC_50_, multiple purine salvage and degradation pathways were enriched (guanine/guanosine salvage, guanine/guanosine salvage II, guanosine nucleotide degradation III, purine deoxyribonucleoside degradation, and purine ribonucleoside degradation), together with pyrimidine deoxyribonucleotide phosphorylation, indicating intensified nucleotide recycling and rebalancing likely linked to DNA damage management. NAcGlnGln amide biosynthesis was also enriched, consistent with activation of a broader stress-resilience program, and serine degradation suggested accompanying adjustments in amino acid utilization under repair-associated demand.

At 1×IC_50_, enrichment shifted toward combined amino acid remodeling, lipid remodeling, and continued nucleotide pressure, with ornithine degradation I (proline biosynthesis) consistent with amino-acid reconfiguration relevant to redox balance (249), palmitate biosynthesis II indicating membrane lipid adaptation, and enrichment of 5-aminoimidazole ribonucleotide biosynthesis pathways supporting sustained demand on de novo purine synthesis (250). At 2× IC_50_, the enrichment landscape broadened, with persistent purine-related pathways (purine deoxyribonucleoside degradation and salvage) alongside pathways indicative of alternative carbon utilization (ribose degradation, L-arabinose degradation, and androstenedione degradation), and enrichment of lipoate biosynthesis and incorporation I and anaerobic heme biosynthesis, consistent with remodeling of energy-generating systems under severe stress (251, 252). Notably, guanylyl MoCo biosynthesis appeared at 2× IC_50_, suggesting engagement of molybdoenzyme-linked redox metabolism primarily under the highest ciprofloxacin pressure.

At the metabolite level, NAcGlnGln was up-regulated at 0.5× and 1× IC_50_ (log_2_FC = +4.26 and +4.93, respectively; Figure 6B), consistent with the pathway enrichment and with its role as a conserved bacterial stress-associated metabolite. MoO_2_-molybdopterin cofactor was also increased at 2× IC_50_ (log_2_FC = +3.61) (Figure 6A), paralleling the induction of molybdenum cofactor-related pathways seen under other antibiotic exposures and supporting activation of redox-associated metabolism at high ciprofloxacin stress.

#### 3.3.4 Methicillin

Methicillin exposure induced a dose-dependent metabolic response consistent with β-lactam–associated envelope stress coupled to compensatory adjustments in energy and redox metabolism (Figure S17).

At 0.5× IC_50_, enriched pathways indicated combined remodeling of nucleotide turnover, central carbon metabolism, and cell-envelope homeostasis. Enrichment of pyrimidine deoxyribonucleosides degradation was consistent with increased nucleotide recycling, while perturbation of mannitol degradation, glycolysis, and thiamine diphosphate biosynthesis suggested reallocation of energy metabolism and cofactor supply under stress (253, 254). Of particular interest is the enrichment of teichoic acid biosynthesis, which is directly linked to MRSA cell wall structure and methicillin resistance. The enrichment of both aerobic and anaerobic heme biosynthesis suggests changes in respiratory activity, potentially to balance energy demand imposed by cell wall stress (255).

At 1× IC_50_, enrichment became more centered on respiratory adaptation and redox/energy homeostasis, including proline to cytochrome *b_0_* oxidase electron transfer and gluconeogenesis, consistent with carbon rerouting toward energy-generating routes under antibiotic pressure (256, 257). Continued enrichment of heme biosynthesis reinforced altered respiratory demand, while changes in fatty acid β-oxidation, lipoate salvage, and valine biosynthesis indicated coordinated adjustments in lipid and amino acid pathways supporting membrane integrity and metabolic capacity (258–260). At 2× IC_50_, enrichment shifted toward amino acid–based energy and stress pathways, including arginine degradation and ornithine degradation, which can support ATP production and redox balancing under strong stress (261, 262). Additional enrichment of queuosine biosynthesis suggested perturbation of tRNA modification systems (263), while recurrent nucleotide degradation pathways indicated sustained nucleotide recycling. Guanylyl MoCo biosynthesis again appeared at high dose, consistent with engagement of redox-associated molybdenum cofactor metabolism under antibiotic pressure, and enrichment of the mevalonate pathway pointed to perturbation of isoprenoid-linked precursors relevant to envelope maintenance (236).

Metabolite inspection supported disruption of respiratory-linked pathways. Protoporphyrinogen IX was down-regulated (Figure 6C), consistent with impaired heme-associated metabolism required for aerobic and anaerobic respiration (255). (*S*)-1-pyrroline-5-carboxylate, an intermediate in proline catabolism linked to electron transfer via cytochrome *b_0_*-associated routes, was also reduced (Figure 6D), supporting coordinated attenuation of proline-mediated respiratory inputs (256, 257). In addition, MoO_2_-molybdopterin accumulation increased progressively with methicillin concentration (Figure 6A), mirroring the recurring induction of MoCo metabolism observed across antibiotics and supporting a conserved redox-adaptive signature in MRSA under antimicrobial stress.

#### 3.3.5 Vancomycin

Vancomycin exposure induced a dose-dependent metabolic response consistent with glycopeptide-driven cell wall stress accompanied by adaptive reprogramming of redox/respiratory functions and envelope-precursor metabolism (Figure S18). Across concentrations, recurrent patterns included enrichment of MoCo biosynthetic routes, modulation of heme biosynthesis pathways, and high-dose perturbation of isoprenoid/mevalonate-derived intermediates relevant to envelope integrity. MoCo-related pathways (guanylyl MoCo biosynthesis and MoCo biosynthesis) were among the top enriched pathways at 1× IC_50_ and 2× IC_50_, paralleling the recurring enrichment observed under other antibiotic classes and supporting a conserved redox-associated response to antimicrobial pressure. Heme biosynthesis (aerobic and anaerobic routes) were enriched at 0.5× IC_50_ and again at 2× IC_50_, consistent with stress-linked remodeling of respiratory metabolism, given the role of heme intermediates in cytochrome-dependent electron transport (255, 264, 265).

At 0.5× IC_50_, enrichment of serine degradation and tRNA charging suggested early shifts in amino acid utilization and translational coupling under sub-inhibitory vancomycin exposure, consistent with growth-rate–linked regulation of charging pathways (266). At 1× IC_50_, several pyrimidine salvage/degradation pathways were enriched, supporting nucleotide supply under increasing stress (239, 240). At 1× IC_50_, several pyrimidine salvage/degradation pathways were enriched, supporting nucleotide supply under increasing stress.

At 2× IC_50_, enrichment extended to the mevalonate pathway, heptaprenyl diphosphate biosynthesis, and *trans,trans*-farnesyl diphosphate biosynthesis, indicating disruption of isoprenoid-related precursors that contribute to teichoic acids, membrane-associated isoprenoids, and respiratory cofactors, consistent with intensified demand for envelope remodeling under strong peptidoglycan inhibition. Although Men enzymes (MenA–MenG) were not dysregulated at the protein level, the upstream precursor pathways were altered, implicating menaquinone supply as a stress-sensitive point. As menaquinone biosynthesis is absent in humans (267), targeting key steps such as MenA or MenG could selectively weaken MRSA respiration and enhance vancomycin efficacy.

Metabolite-level inspection supported a robust molybdenum cofactor signature: MoO_2_-molybdopterin cofactor accumulated at 1× and 2× IC_50_ (log_2_FC = +7.23 and +7.12, respectively; Figure 6A). In contrast, although heme biosynthesis pathways were enriched, individual heme-related metabolites showed only modest changes (|log_2_FC| < 1.5), suggesting limited flux alteration detectable at the metabolite level despite pathway-level signals. Overall, vancomycin drove a coordinated metabolic response combining conserved redox-associated remodeling with dose-dependent shifts in nucleotide handling and high-dose disruption of isoprenoid/envelope precursor pathways required to sustain cell envelope integrity under glycopeptide stress.

## 4 Conserved adaptive themes in MRSA: therapeutic opportunities

Across all five antibiotics, MRSA responded through a set of recurring adaptive modules supported by concordant evidence across omics and enrichment analyses. Rather than simply reflecting drug-specific effects, these responses define a unified survival strategy in which MRSA preserves redox and information-processing functions, reallocates resources toward envelope maintenance and nutrient acquisition, and retains the capacity for rapid adaptation under antimicrobial pressure. The main conserved themes were metal/cofactor homeostasis, sustained pressure on nucleotide and folate metabolism, genetic plasticity through DNA repair and prophage-linked functions, and remodeling of the surfaceome and transport systems. Because these modules were repeatedly engaged across antibiotic classes (Figure 7), each also points to specific vulnerabilities that may be exploited therapeutically.

**Figure 7.**
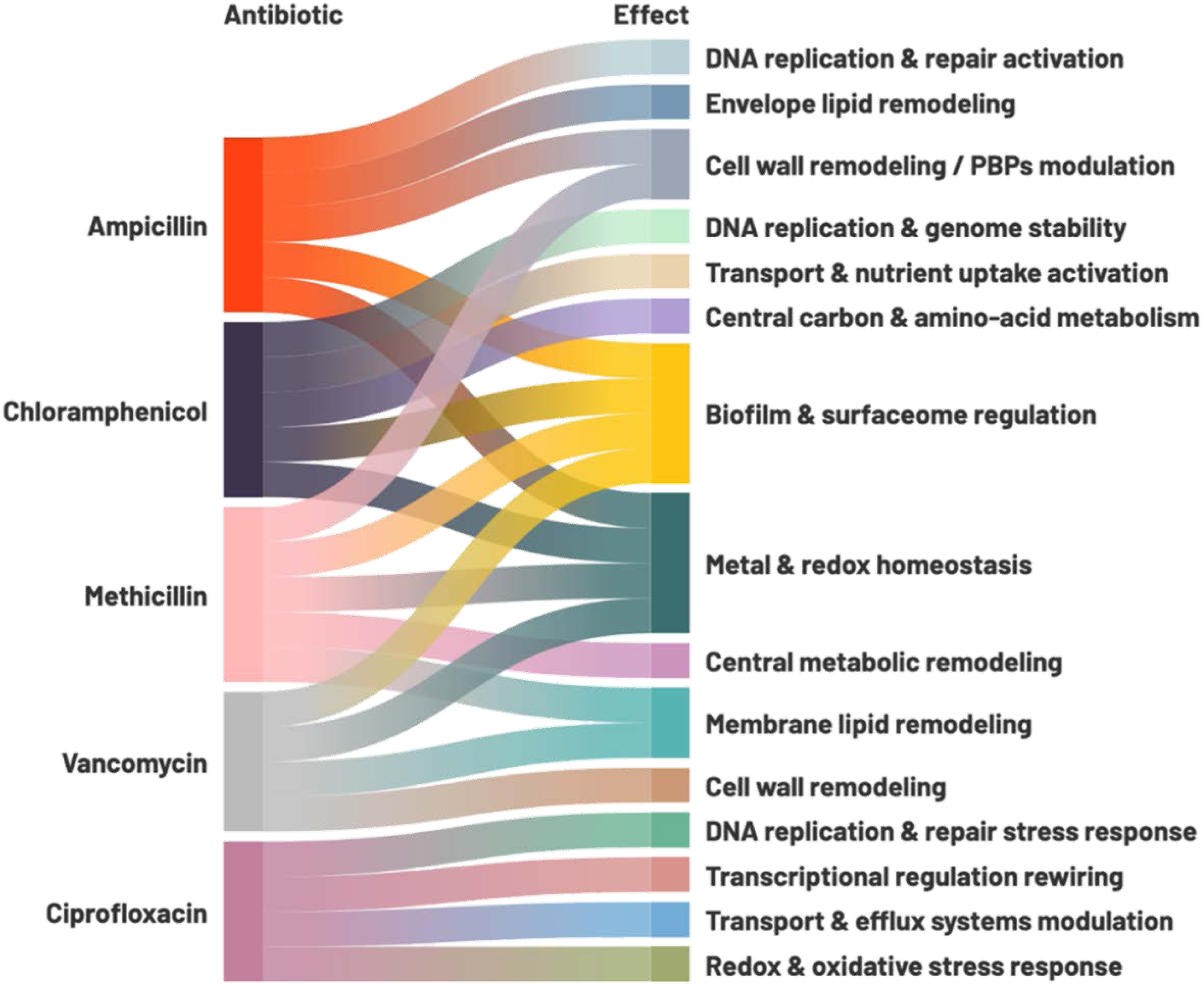
Cross-antibiotic convergence of responses identified in MRSA. Each antibiotic is linked to the major response modules supported by integrated proteomic, post-translational modification, metabolomic, lipidomic, and enrichment analyses. Flows connect ampicillin, chloramphenicol, ciprofloxacin, methicillin, and vancomycin to the principal adaptive effects identified in this study, including DNA replication and repair activation, genome stability responses, transcriptional regulation rewiring, metal/redox homeostasis, cell-wall and membrane remodeling, transport and nutrient uptake activation, central metabolic remodeling, and biofilm/surfaceome regulation. The figure provides a qualitative overview of recurrent and antibiotic-specific adaptive patterns rather than a quantitative weighting of pathway strength.

A first conserved layer was metal and cofactor homeostasis. Across antibiotics, multi-omics evidence repeatedly implicated remodeling of Fe-, Ni-, heme-, and Mo-linked pathways (summarized in Table S2). β-Lactams prominently affected iron acquisition, including altered siderophore-dependent iron import and changes in iron-binding or transport proteins such as Q2FWN6 and SirA/Q2G1N4, whereas vancomycin induced the heme transport permease HrtB/Q2G168 at 1× and 2× IC50 (log_2_FC = +2.64 and +1.90). Nickel-responsive features also recurred, with induction of the Ni(II)-dependent urease subunit γ ureA/Q2FVW5 under chloramphenicol ( 2× IC_50_: log_2_FC +1.60) and methicillin (2× IC_50_: log_2_FC +1.50), while high-dose vancomycin repressed the nickel-binding protein NikA/Q2G2P5. The strongest cross-antibiotic metabolite-level signature in the dataset was the consistent accumulation of MoO_2_-molybdopterin cofactor under all five antibiotics, with large effect sizes under ampicillin (log_2_FC = +4.53 to +6.80), chloramphenicol (+0.98 to +2.57), ciprofloxacin (+3.61 at 2× IC50), methicillin (+1.96 to +3.32), and vancomycin (+7.23 and +7.12). This axis was further supported by induction of the molybdenum ABC transporter Q2FVX4 under methicillin 0.5× IC50 (log_2_FC = +3.02) and phosphorylation of the MOSC-domain protein Q2FVS9 under ampicillin. Together, these data indicate that MRSA repeatedly rewires metal and cofactor handling to preserve redox balance and metabolic function under stress. The recurrent, high-magnitude regulation of MoCo metabolism, HrtB, and UreA supports these nodes as plausible adjuvant targets, as co-inhibition could weaken a conserved metal-dependent stress-buffering layer.

A second major module was sustained pressure on nucleotide and folate metabolism. Across antibiotics, the data converged on continued demand for nucleotide homeostasis, salvage, and folate-linked precursor supply, consistent with the need to maintain dNTP pools and repair capacity even when the primary drug target lies outside nucleic-acid metabolism (Table S3). Proteomics supported compensatory reinforcement of dNTP supply, including induction of ribonucleoside-diphosphate reductase Q2G078 under ampicillin (0.5xIC_50_: log_2_FC +3.32) and NrdI/Q2G079 under chloramphenicol (1× and 2×IC_50_: log_2_FC +2.74 and +2.57, respectively). PTMs added a further regulatory layer, including increased acetylation of the DinG exonuclease under ampicillin, dephosphorylation of UvrA/Q2G046 under ciprofloxacin (1× and 2×IC_50_, log_2_FC = −2.46 and −2.04), reduced acetylation of PyrG/Q2FWD1 under vancomycin, and recurrent repression of aspartokinase/Q2FYP1 under ciprofloxacin and vancomycin. These convergent signals indicate that MRSA prioritizes maintenance of nucleotide pools and repair competence under diverse antibiotic stresses.

A third conserved theme was genetic plasticity through DNA repair, replication control, and prophage-associated functions. Across antibiotics with distinct primary targets, MRSA repeatedly engaged genome-maintenance systems, supporting a general requirement to preserve chromosome integrity under antimicrobial pressure (Table S4). Under ampicillin, DNA ligase LigA/Q2G1Y0 was strongly induced (0.5× and 2× IC_50_: log_2_FC +2.80 and +3.04), while RadA increased at 0.5× IC_50_ (log2FC = +1.94), consistent with reinforcement of DNA repair capacity. Methicillin further strengthened this pattern through induction of AddB/Q2FZT6 (1–2× IC_50_: log_2_FC +4.02 and +3.98) and RecF/Q2G275 (0.5× IC_50_: log_2_FC +2.52), highlighting recombination-associated repair as a major survival axis under β-lactam stress. Ciprofloxacin, as expected for a genotoxic antibiotic, showed strong DNA-defense signatures, including restriction-modification enrichment, induction of an adenine-specific methyltransferase, dephosphorylation of UvrA (down-regulated at 1–2× IC_50_, log_2_FC −2.46 and −2.04), and induction of replication-initiation control through YabA. Prophage-linked responses were more heterogeneous, with ampicillin and methicillin inducing structural or mobilization-related proteins, whereas ciprofloxacin repressed a phage terminase and vancomycin down-regulated a hypothetical phage protein, indicating that prophage engagement is context dependent rather than universal. Overall, these data identify LigA, AddB/RecF-associated repair capacity, and PTM-regulated repair functions such as UvrA as plausible sensitization targets, because they mark core genome-stabilizing responses that buffer antibiotic stress.

A fourth conserved layer was remodeling of the surfaceome and transport systems. Integrated proteomics and PTM data showed that MRSA repeatedly reprogrammed its cell-environment interface by modulating nutrient uptake, transport regulation, surface-associated proteins, and envelope-remodeling functions (Table S5). A consistent feature under cell-wall stress was repression of major virulence-control systems, including down-regulation of *agrB* under methicillin and vancomycin, repression of YycH under vancomycin, and reduction of Sortase A under methicillin, suggesting that MRSA diverts resources away from energetically costly virulence programs during survival prioritization. In parallel, multiple antibiotics increased transport capacity, particularly through amino acid ABC transporters, MFS transporters, and PTS components.

Chloramphenicol induced the amino acid ABC transporter Q2FX87 (1–2× IC_50_: log_2_FC +1.57 and +2.66) and the PTS EIICB component Q2G2D4 (2× IC_50_: log_2_FC +2.36), while vancomycin induced the MFS transporter Q2G2B3 (1× IC_50_: log_2_FC +3.54) and showed strong PTM up-regulation on EcfT/Q2FZI4 (N200 deamidation, 1–2× IC_50_: log_2_FC +3.45 and +3.74). Ciprofloxacin contributed a complementary signature, with multi-site dephosphorylation of the ABC transporter domain protein Q2G196 (log_2_FC −1.96 to −2.43), indicating active transporter regulation under genotoxic stress. Surface-matrix and envelope-remodeling functions were also repeatedly affected, notably through recurrent down-regulation of the putative polysaccharide biosynthesis protein Q2G0R7 under ampicillin and vancomycin, with additional multi-site PTM remodeling under vancomycin. Ciprofloxacin further reinforced this theme through induction of PBP1/Q2FZ94 across all concentrations (log_2_FC = +2.88, +1.67, and +2.39), consistent with compensatory cell-wall maintenance. Together, these data support a conserved adaptive module in which MRSA increases nutrient acquisition and envelope maintenance while reducing investment in selected virulence-associated surface functions. Within this module, HrtB, PBP1, and selected ABC, MFS, PTS, and ECF transport nodes emerge as plausible adjuvant targets because they are repeatedly and strongly engaged across antibiotic conditions.

## 5 Conclusions

This study provides a multi-omics view of how MRSA ATCC 43300 responds to antibiotic stress across five mechanistically distinct antibiotics and graded inhibitory concentrations. Despite the diversity of primary drug targets, the responses did not resolve into fully separate drug-specific programs. Instead, antibiotic exposure repeatedly converged on a limited set of conserved adaptive modules detectable across proteomics, post-translational modifications, and metabolomics.

Overall, the data support four main conclusions. First, envelope stress and genome maintenance are closely linked in MRSA, indicating that preservation of chromosome integrity is a core requirement even during predominantly cell-wall-directed stress. Second, metal/cofactor and redox homeostasis constitute a recurrent stress-buffering layer, with consistent remodeling of Fe-, Ni-, heme-, and Mo-linked functions and a particularly strong cross-antibiotic MoCo signature. Third, nucleotide and folate metabolism represent a major adaptive pressure point, with recurring evidence of perturbed purine metabolism, reinforced dNTP supply, and repair-associated responses. Fourth, MRSA repeatedly reprograms its surface and transport interface, coupling nutrient acquisition and envelope maintenance with reduced investment in selected virulence-associated functions.

These findings support a unified model in which MRSA tolerates antibiotic stress not by deploying entirely distinct responses to each drug, but by repeatedly engaging a compact set of conserved stress-buffering functions. From a therapeutic perspective, this convergence highlights several candidate adjuvant vulnerabilities, particularly within metal/cofactor homeostasis, nucleotide supply and repair, and transport/envelope compensation programs.

This work should also be interpreted considering its design. The study was exploratory by design, and as such results define candidate adaptive patterns rather than population-level estimates of biological variability. In addition, the proposed vulnerabilities are inferred from molecular convergence and regulation, not from direct functional perturbation. Accordingly, the identified modules and targets now require validation in independently replicated cultures and targeted mechanistic experiments.

Even with these limitations, the consistency of the cross-omics and cross-antibiotic signals indicates that MRSA adaptation is organized around a relatively small number of conserved molecular solutions. This provides a framework for future work aimed at testing whether disruption of these recurrent adaptive layers can sensitize MRSA to antibiotic treatment and reduce stress tolerance or persistence.

## Supporting Information

Figures S1 to S18 and Tables S1 to S5 are available as Supporting Information.

## Author contributions: CRediT

**PCR** – Conceptualization, Investigation, Writing – original draft; **PFP** – Investigation, Resources; **MMM** – Resources, Writing – review and editing; **GCJ** – Methodology, Supervision, Writing – review and editing.

## Data Availability

Mass spectrometry datasets are available online at https://doi.org/10.5281/zenodo.19457185, https://doi.org/10.5281/zenodo.19457501, https://doi.org/10.5281/zenodo.19457546, https://doi.org/10.5281/zenodo.19457810, https://doi.org/10.5281/zenodo.19458205, https://doi.org/10.5281/zenodo.19459024, https://doi.org/10.5281/zenodo.19460939, https://doi.org/10.5281/zenodo.19461672, https://doi.org/10.5281/zenodo.19461883, https://doi.org/10.5281/zenodo.19462038, https://doi.org/10.5281/zenodo.19462199, https://doi.org/10.5281/zenodo.19462351, https://doi.org/10.5281/zenodo.19462432, https://doi.org/10.5281/zenodo.19462600, https://doi.org/10.5281/zenodo.19462710, and https://doi.org/10.5281/zenodo.19462885. Proteomic MaxQuant processing files are available online at http://doi.org/10.5281/zenodo.20179627.

