## Supporting Information for "Omics analysis of MRSA under antibiotic stress identifies conserved adaptive modules and candidate adjuvant targets"

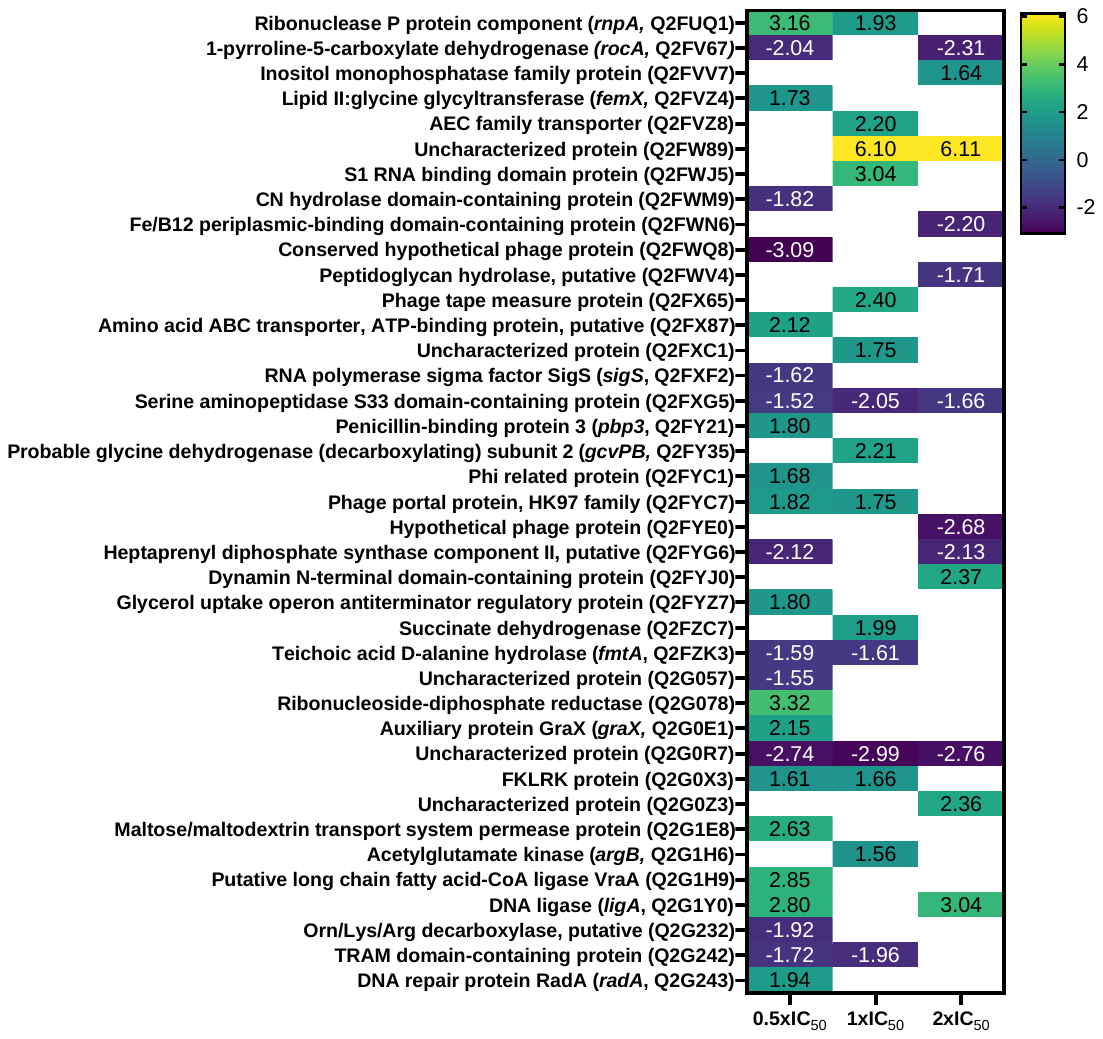

Figure S1 – Significantly dysregulated proteins in MRSA 43300 in the presence of different IC50 values of ampicillin.

Heatmap displays log_2_FC values for significantly dysregulated proteins across different ampicillin concentrations. Positive values indicate up-regulation; negative values indicate downregulation. Blank cells indicate proteins that were detected under that condition but did not meet the criteria for statistical significance (*p*-value < 0.05, and a |log_2_FC| ≥ 1.5). Protein names are listed with corresponding gene IDs (when available) and UniProt IDs. FC ‑ fold‑change.

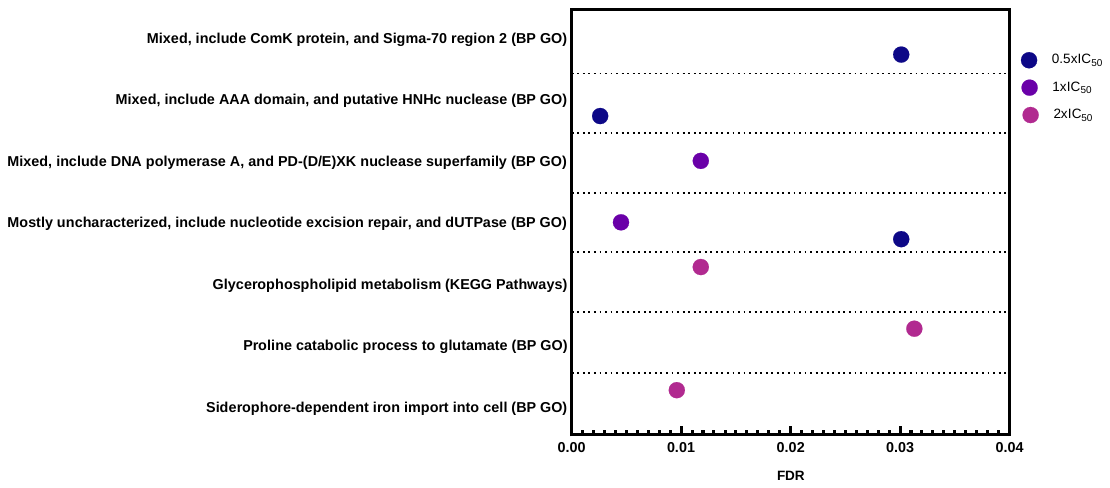

Figure S2 – STRING functional enrichments for MRSA response to ampicillin.

Dot plots show significantly enriched gene ontology (GO)-derived biological processes (BP) and KEGG pathways for proteins dysregulated at different ampicillin concentrations (0.5, 1, and 2xIC_50_). The x-axis represents the false discovery rate (FDR), with lower values indicating higher statistical significance. Circle colors distinguish between the three ampicillin concentrations. Only pathways with FDR < 0.05 were considered statistically significant.

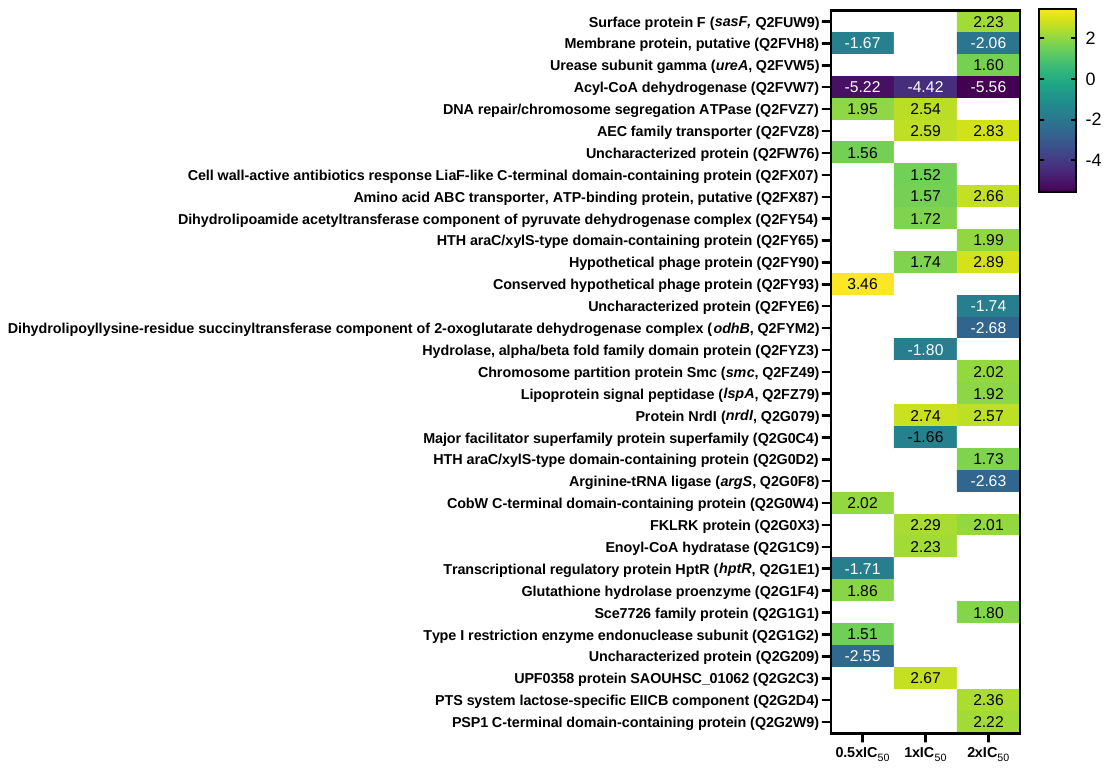

Figure S3 – Significantly dysregulated proteins in MRSA 43300 in the presence of different IC_50_ values of chloramphenicol.

Heatmap displays log_2_FC values for significantly dysregulated proteins across different chloramphenicol concentrations. Positive values indicate up-regulation; negative values indicate downregulation. Blank cells indicate proteins that were detected under that condition but did not meet the criteria for statistical significance (*p*‑value < 0.05, and a |log_2_FC| ≥ 1.5). Protein names are listed with corresponding gene IDs (when available) and UniProt IDs. FC ‑ fold‑change.

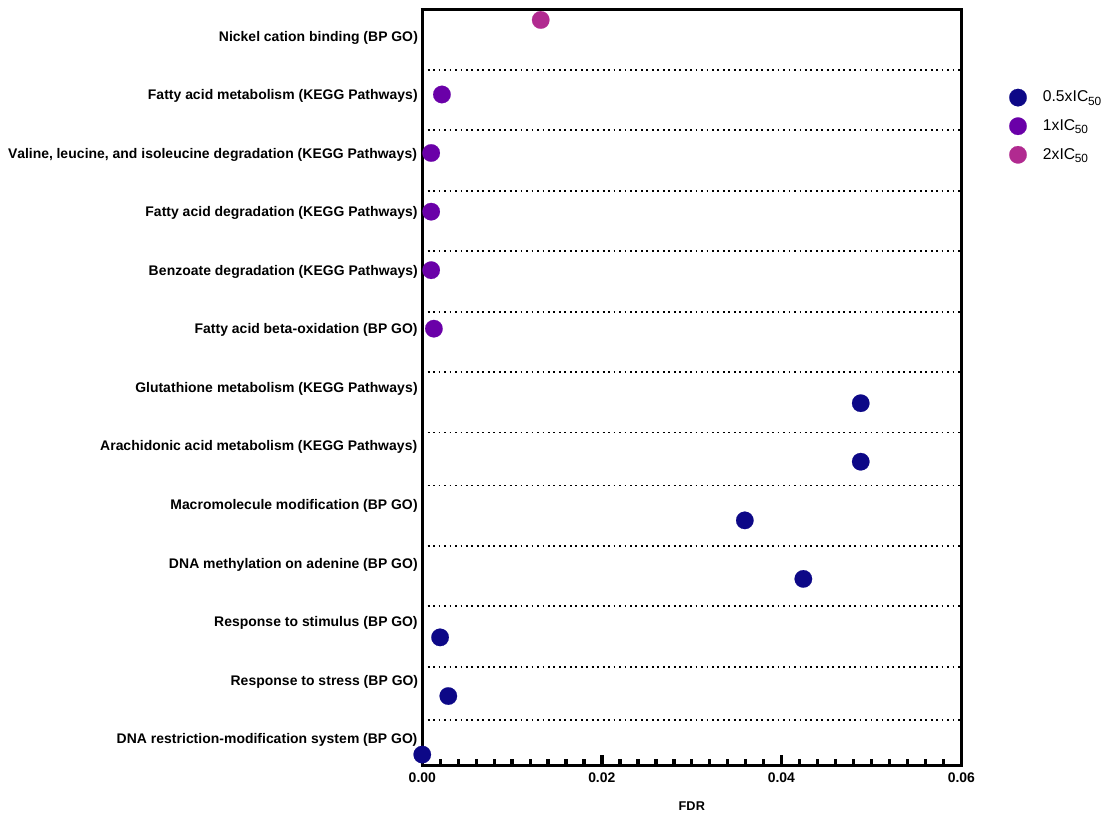

Figure S4 – STRING functional enrichments for MRSA response to chloramphenicol.

Dot plots show significantly enriched biological processes (BP) and KEGG pathways for proteins dysregulated at different chloramphenicol concentrations (0.5, 1, and 2xIC_50_). The x-axis represents the false discovery rate (FDR), with lower values indicating higher statistical significance. Circle colors distinguish between the three chloramphenicol concentrations. Only pathways with FDR < 0.05 were considered statistically significant.

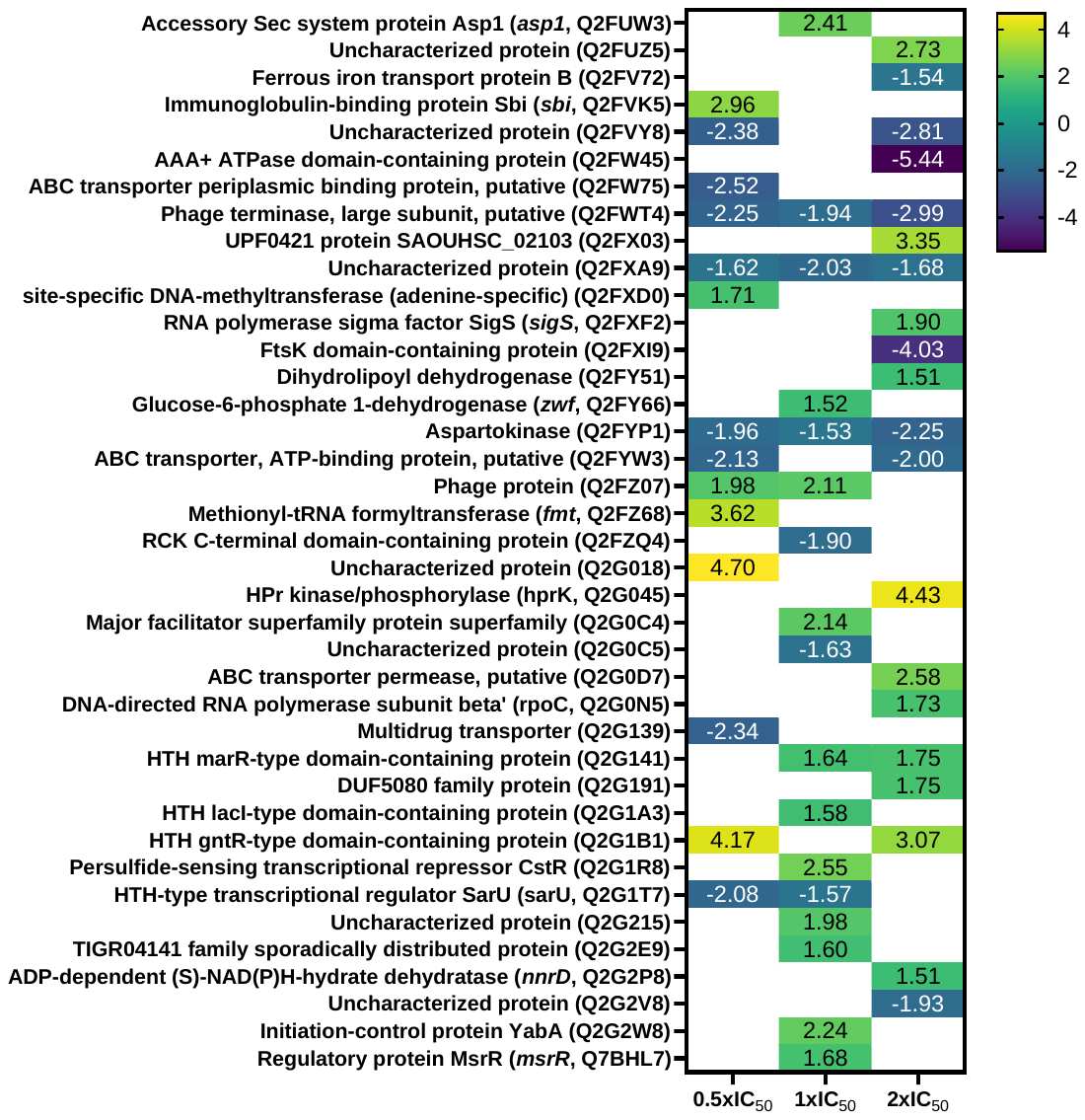

Figure S5 – Significantly dysregulated proteins in MRSA 43300 in the presence of different IC_50_ values of ciprofloxacin.

Heatmap displays log_2_FC values for significantly dysregulated proteins across different ciprofloxacin concentrations. Positive values indicate up-regulation; negative values indicate downregulation. Blank cells indicate proteins that were detected under that condition but did not meet the criteria for statistical significance (*p*-value < 0.05, and a |log_2_FC| ≥ 1.5). Protein names are listed with corresponding gene IDs (when available) and UniProt IDs. FC ‑ fold‑change.

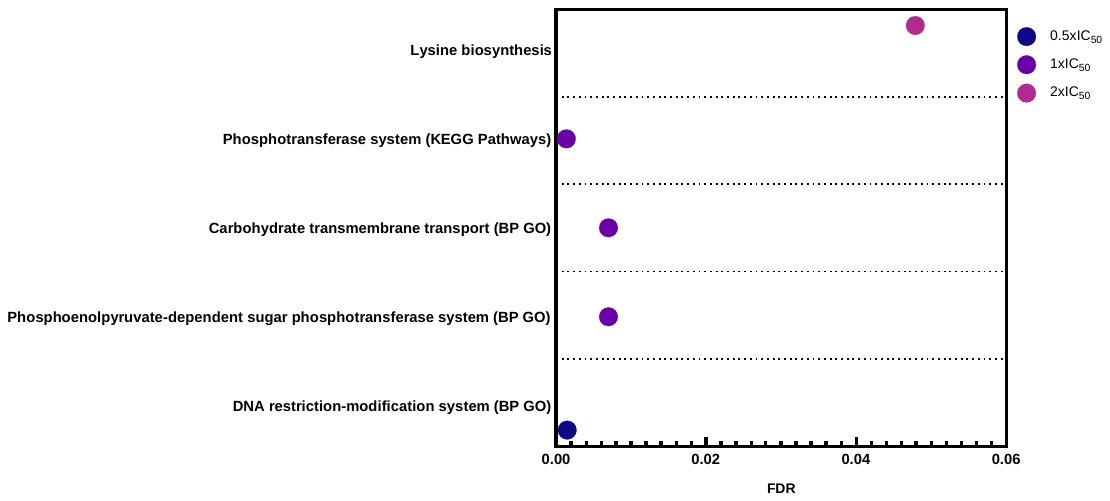

Figure S6 – STRING functional enrichments for MRSA response to ciprofloxacin.

Dot plots show significantly enriched biological processes (BP) and KEGG pathways for proteins dysregulated at different ciprofloxacin concentrations (0.5, 1, and 2xIC_50_). The x-axis represents the false discovery rate (FDR), with lower values indicating higher statistical significance. Circle colors distinguish between the three ciprofloxacin concentrations. Only pathways with FDR < 0.05 were considered statistically significant.

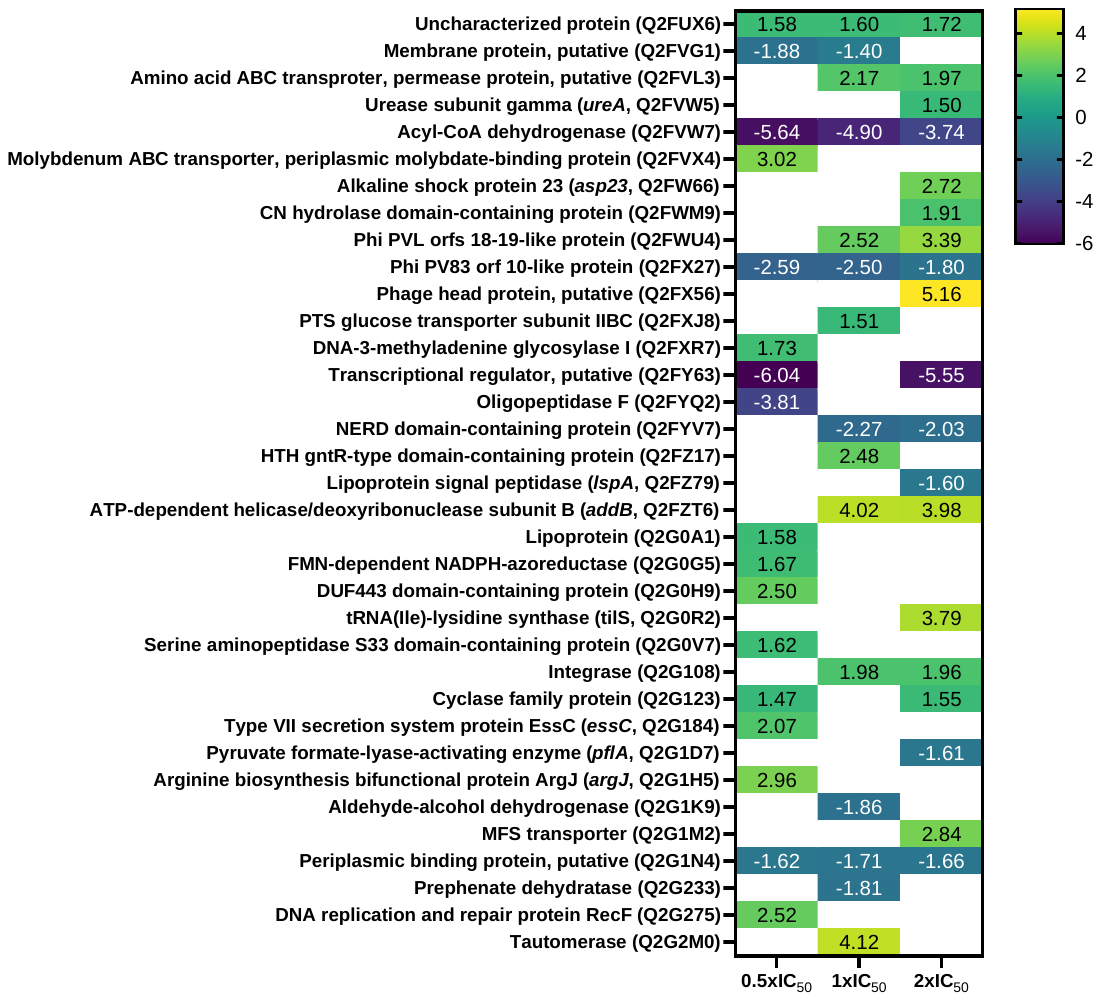

Figure S7 – Significantly dysregulated proteins in MRSA 43300 in the presence of different IC_50_ values of methicillin.

Heatmap displays log_2_FC values for significantly dysregulated proteins across different methicillin concentrations. Positive values indicate up-regulation; negative values indicate downregulation. Blank cells indicate proteins that were detected under that condition but did not meet the criteria for statistical significance (*p*-value < 0.05, and a |log_2_FC| ≥ 1.5). Protein names are listed with corresponding gene IDs (when available) and UniProt IDs. FC ‑ fold‑change.

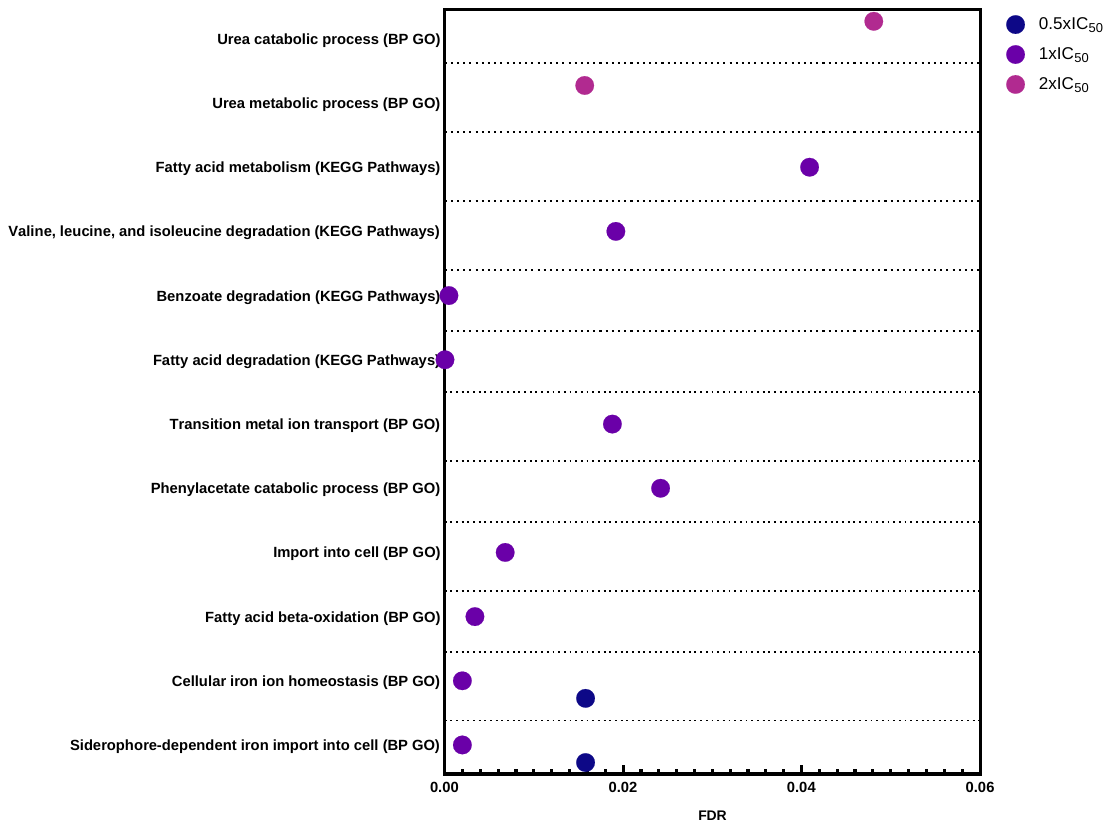

Figure S8 – STRING functional enrichments for MRSA response to methicillin.

Dot plots show significantly enriched biological processes (BP) and KEGG pathways for proteins dysregulated at different methicillin concentrations (0.5, 1, and 2xIC_50_). The x-axis represents the false discovery rate (FDR), with lower values indicating higher statistical significance. Circle colors distinguish between the three ciprofloxacin concentrations. Only pathways with FDR < 0.05 were considered statistically significant.

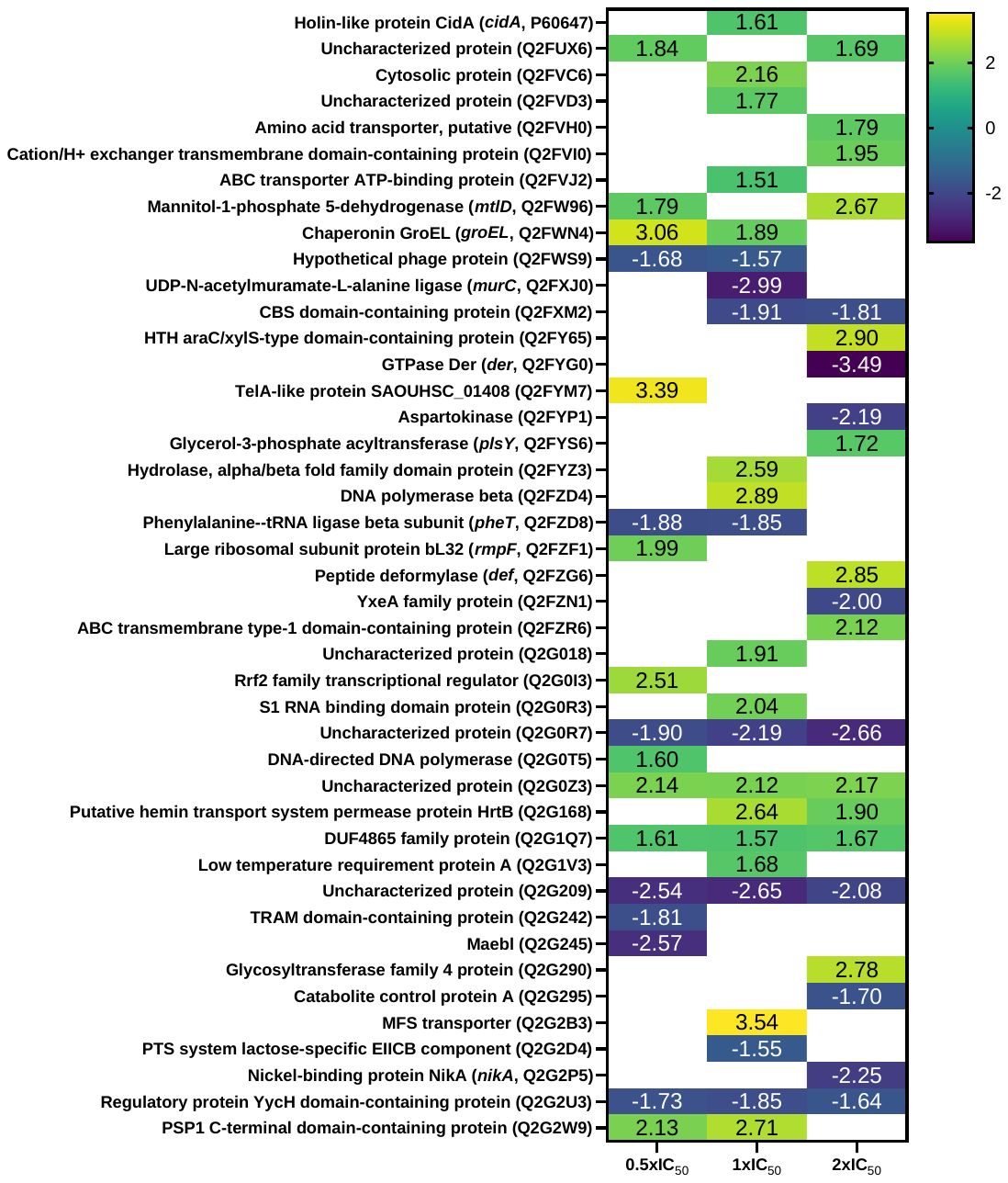

Figure S9  – Significantly dysregulated proteins in MRSA 43300 in the presence of different IC_50_ values of vancomycin.

Heatmap displays log_2_FC values for significantly dysregulated proteins across different vancomycin concentrations. Positive values indicate up-regulation; negative values indicate downregulation. Blank cells indicate proteins that were detected under that condition but did not meet the criteria for statistical significance (*p*-value < 0.05, and a |log_2_FC| ≥ 1.5). Protein names are listed with corresponding gene IDs (when available) and UniProt IDs. FC ‑ fold‑change.

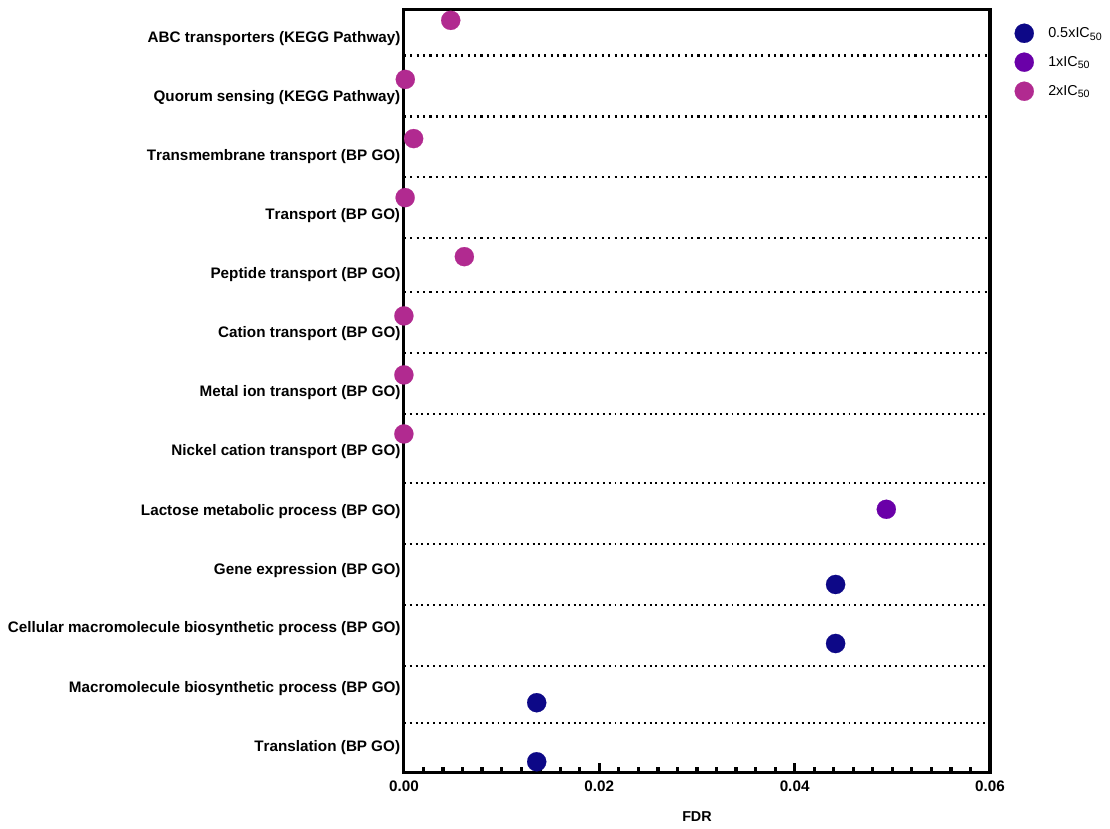

Figure S10 – STRING functional enrichments for MRSA response to vancomycin.

Dot plots show significantly enriched biological processes (BP) and KEGG pathways for proteins dysregulated at different vancomycin concentrations (0.5, 1, and 2xIC_50_). The x-axis represents the false discovery rate (FDR), with lower values indicating higher statistical significance. Circle colors distinguish between the three ciprofloxacin concentrations. Only pathways with FDR < 0.05 were considered statistically significant.

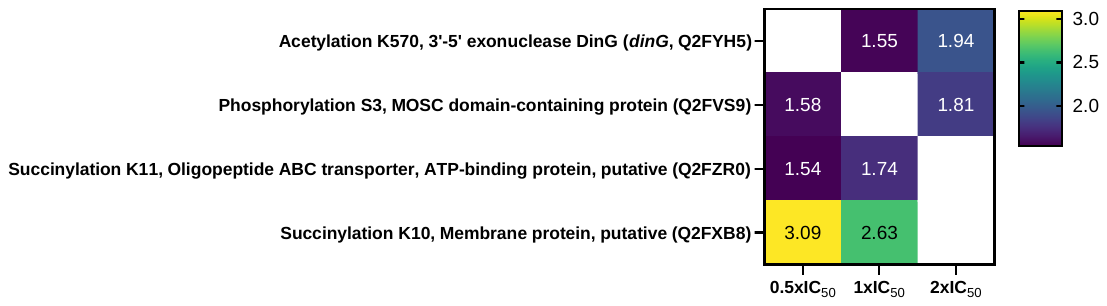

Figure S11 – Significantly dysregulated PTMs in MRSA 43300 in the presence of different IC_50_ values of ampicillin.

Heatmap displays log_2_FC values for significantly dysregulated PTMs across different ampicillin concentrations. Positive values indicate up-regulation; negative values indicate downregulation. Blank cells indicate proteins that were detected under that condition but did not meet the criteria for statistical significance (*p*-value < 0.05, and a |log_2_FC| ≥ 1.5). Protein names are listed with corresponding gene IDs (when available) and UniProt IDs. FC ‑ fold‑change.

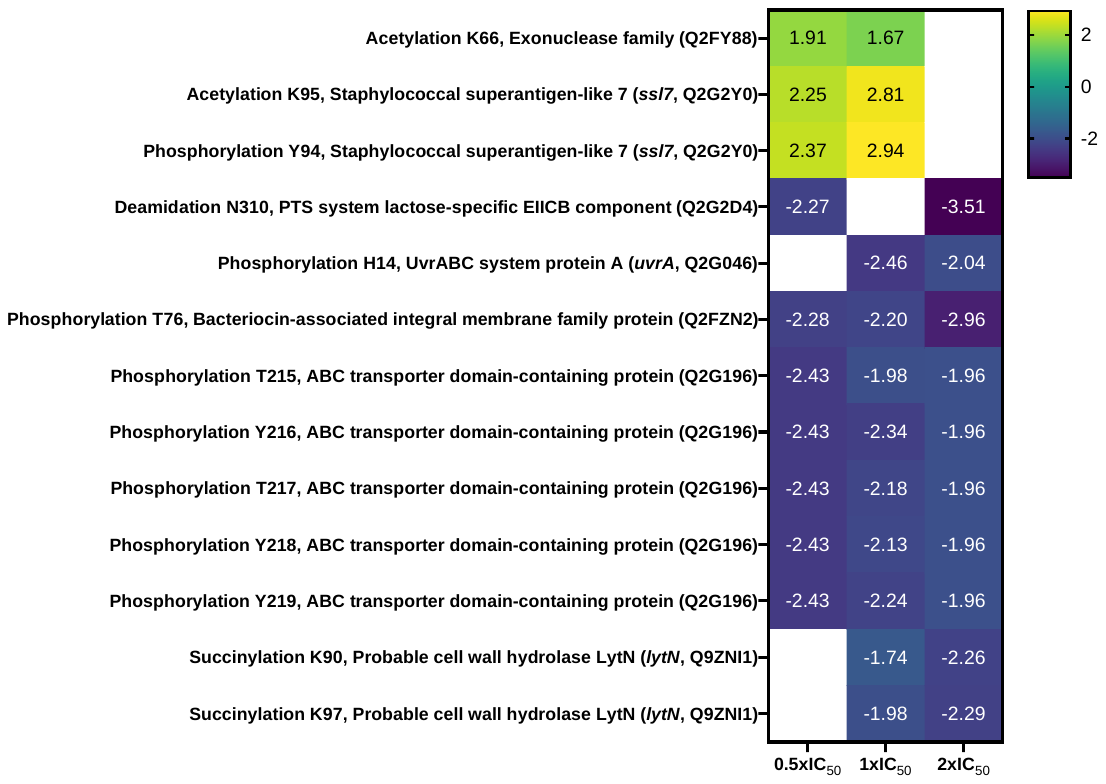

Figure S12 – Significantly dysregulated PTMs in MRSA 43300 in the presence of different IC_50_ values of ciprofloxacin.

Heatmap displays log_2_FC values for significantly dysregulated PTMs across different ciprofloxacin concentrations. Positive values indicate up-regulation; negative values indicate downregulation. Blank cells indicate proteins that were detected under that condition but did not meet the criteria for statistical significance (*p*-value < 0.05, and a |log_2_FC| ≥ 1.5). Protein names are listed with corresponding gene IDs (when available) and UniProt IDs. FC ‑ fold‑change.

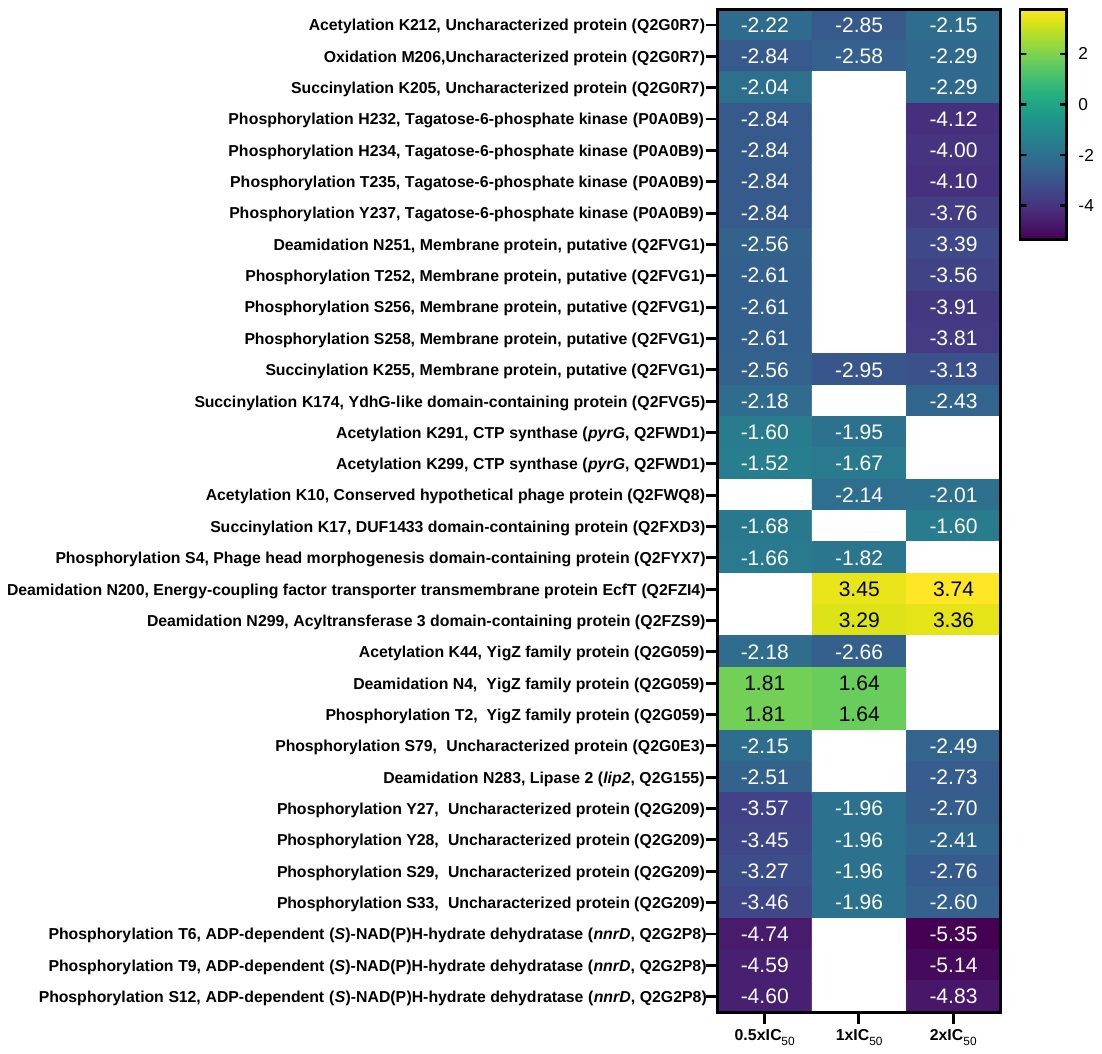

Figure S13 – Significantly dysregulated PTMs in MRSA 43300 in the presence of different IC_50_ values of vancomycin.

Heatmap displays log_2_FC values for significantly dysregulated PTMs across different vancomycin concentrations. Positive values indicate up-regulation; negative values indicate downregulation. Blank cells indicate proteins that were detected under that condition but did not meet the criteria for statistical significance (*p*-value < 0.05, and a |log_2_FC| ≥ 1.5). Protein names are listed with corresponding gene IDs (when available) and UniProt IDs. FC ‑ fold‑change.

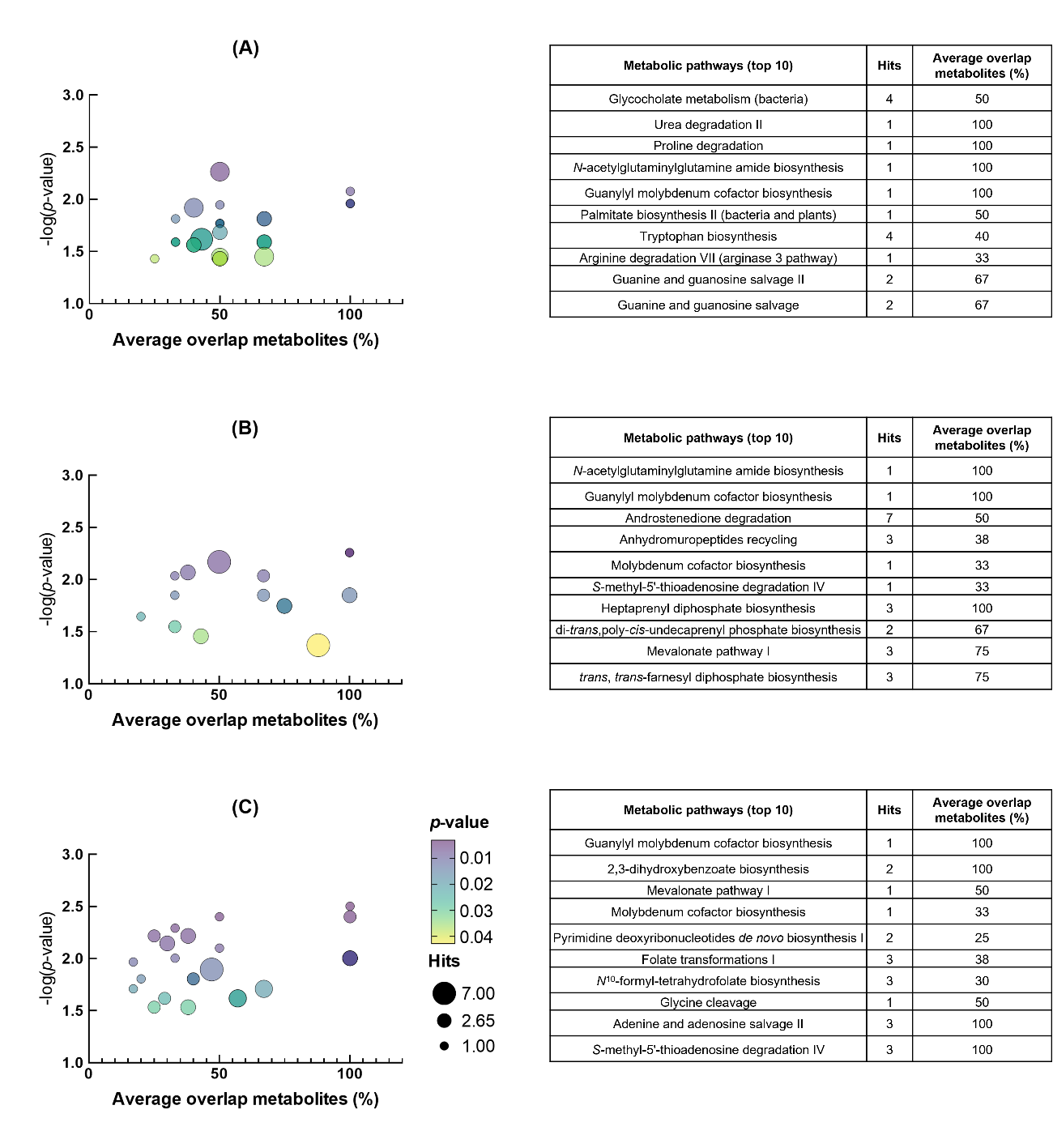

Figure S14 – Metabolic pathways significantly altered in MRSA 43300 in the presence of different IC_50_ values of ampicillin.

(A) 0.5xIC_50_, (B) 1xIC_50_, (C) 2xIC_50_ ampicillin. Cloud plots on the left show significantly altered pathways and their metabolite overlap (fraction of identified altered metabolites relative to total number of metabolites that constitute the pathway), as well as the number of hits per pathway. On the right, the top 10 altered pathways are listed. Statistical significance: A and B, *p* < 0.02 and C, *p* < 0.01.

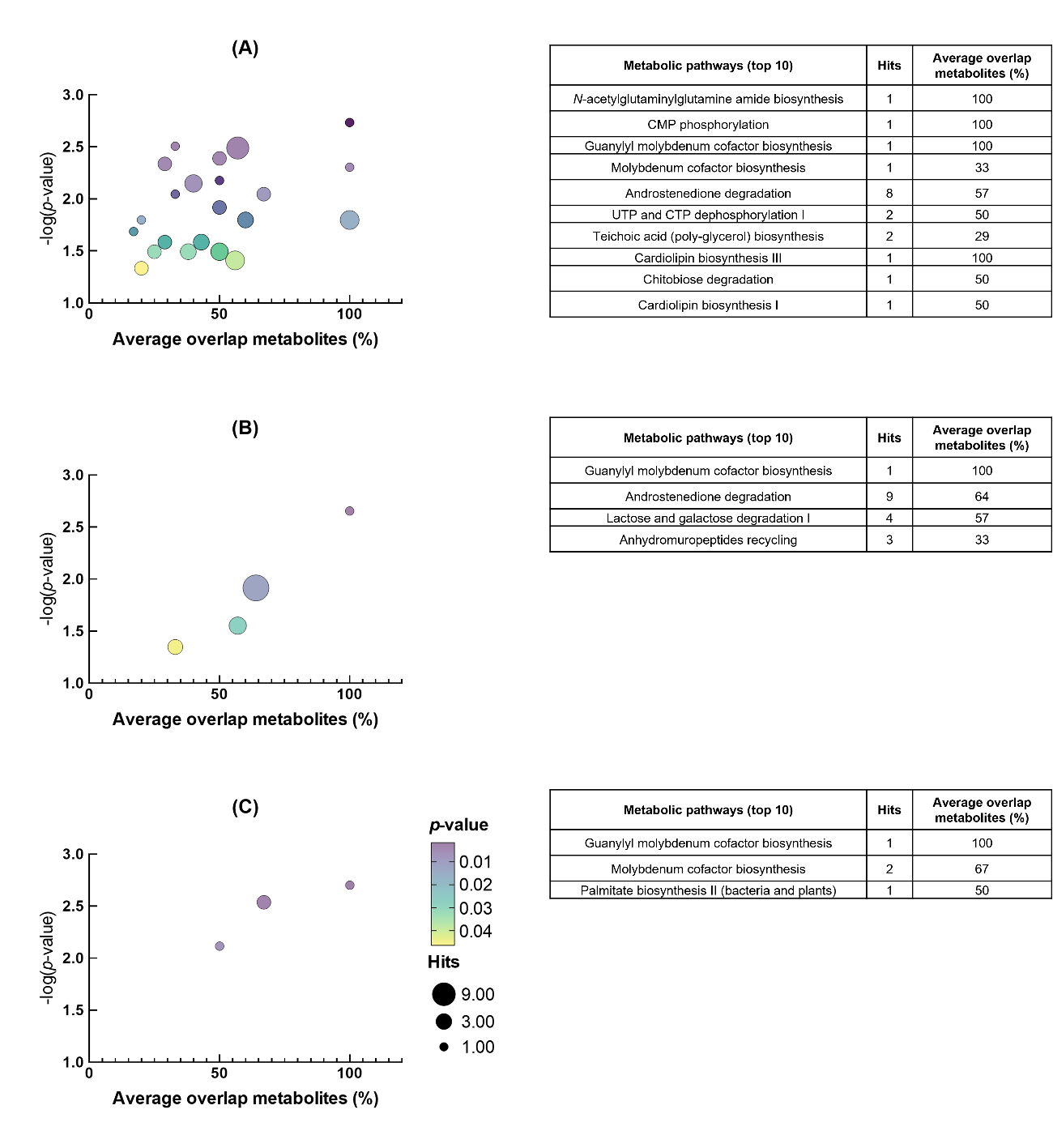

Figure S15 – Metabolic pathways significantly altered in MRSA 43300 in the presence of different IC_50_ values of chloramphenicol.

(A) 0.5xIC_50_, (B) 1xIC_50_, (C) 2xIC_50_ chloramphenicol. Cloud plots on the left show significantly altered pathways and their metabolite overlap (fraction of identified altered metabolites relative to total number of metabolites that constitute the pathway), as well as the number of hits per pathway. On the right, the top 10 altered pathways are listed. Statistical significance: B, *p* < 0.05 and A and C, *p* < 0.01.

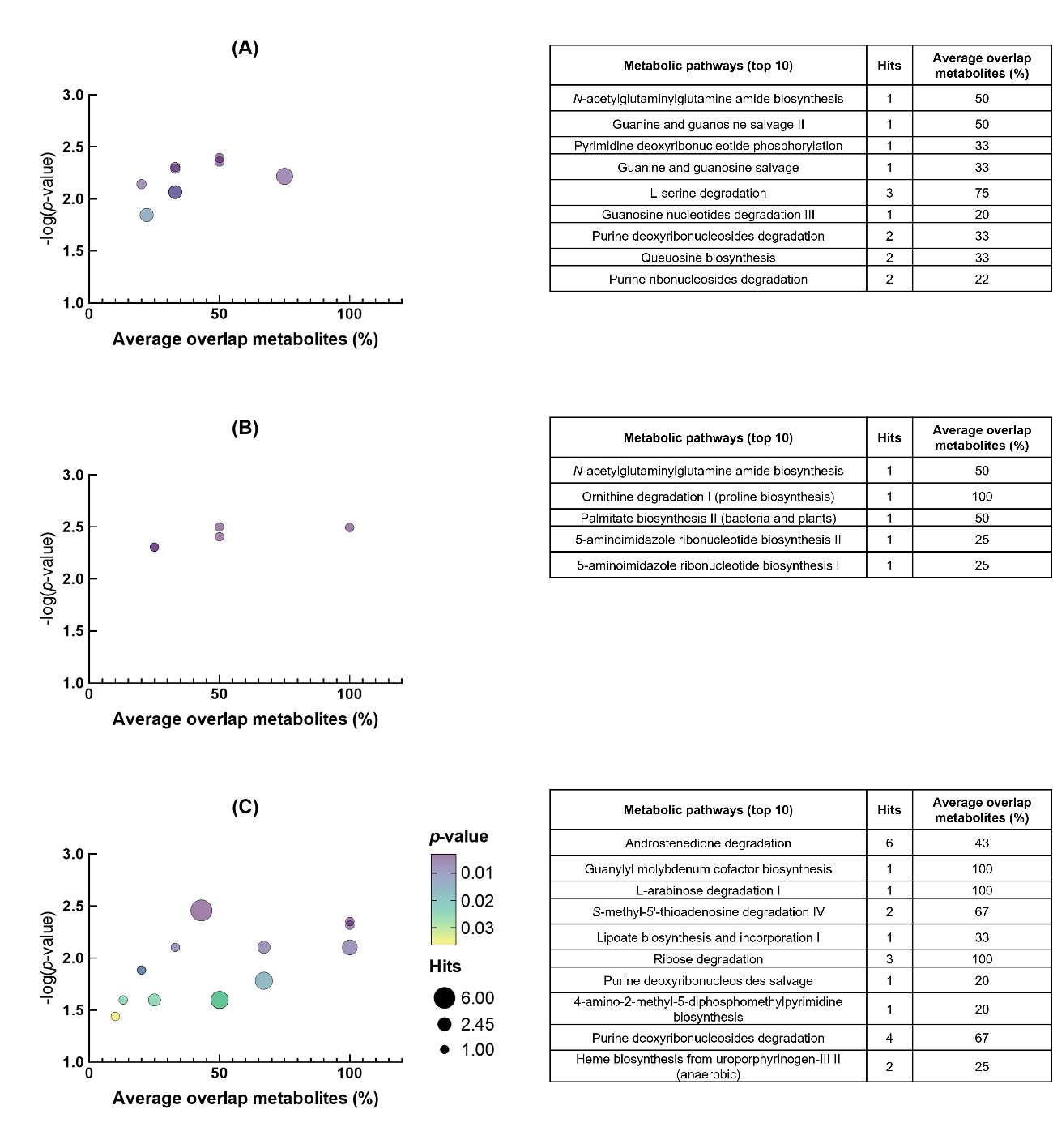

Figure S6 – Metabolic pathways significantly altered in MRSA 43300 in the presence of different IC_50_ values of ciprofloxacin.

(A) 0.5xIC_50_, (B) 1xIC_50_, (C) 2xIC_50_ ciprofloxacin. Cloud plots on the left show significantly altered pathways and their metabolite overlap (fraction of identified altered metabolites relative to total number of metabolites that constitute the pathway), as well as the number of hits per pathway. On the right, the top 10 altered pathways are listed. Statistical significance: C, *p* < 0.05, A, *p* < 0.02, and B, *p* < 0.01.

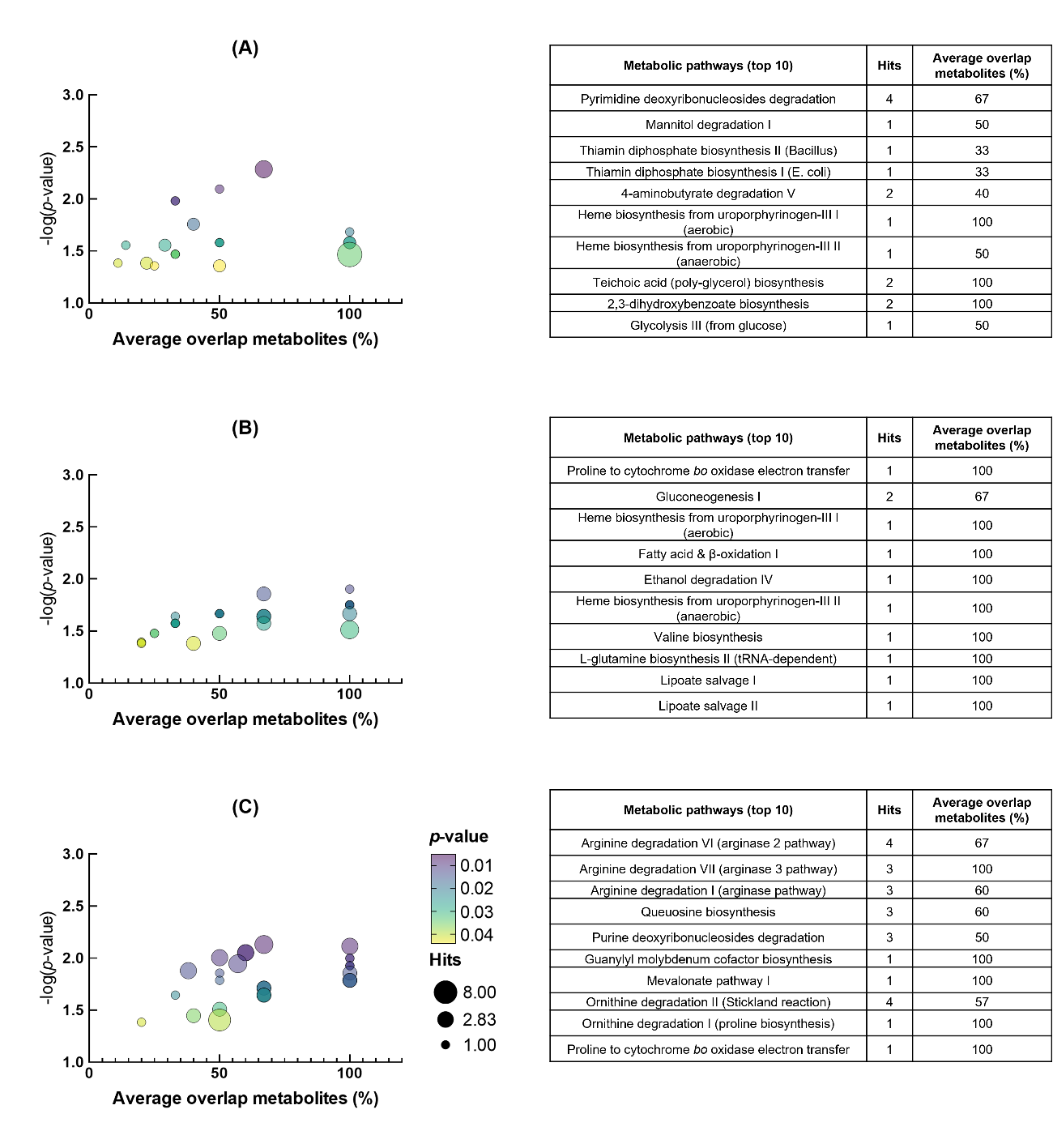

Figure S17 – Metabolic pathways significantly altered in MRSA 43300 in the presence of different IC_50_ values of methicillin.

(A) 0.5xIC_50_, (B) 1xIC_50_, (C) 2xIC_50_ methicillin. Cloud plots on the left show significantly altered pathways and their metabolite overlap (fraction of identified altered metabolites relative to total number of metabolites that constitute the pathway), as well as the number of hits per pathway. On the right, the top 10 altered pathways are listed. Statistical significance: C, *p* < 0.05, A and B, *p* < 0.02.

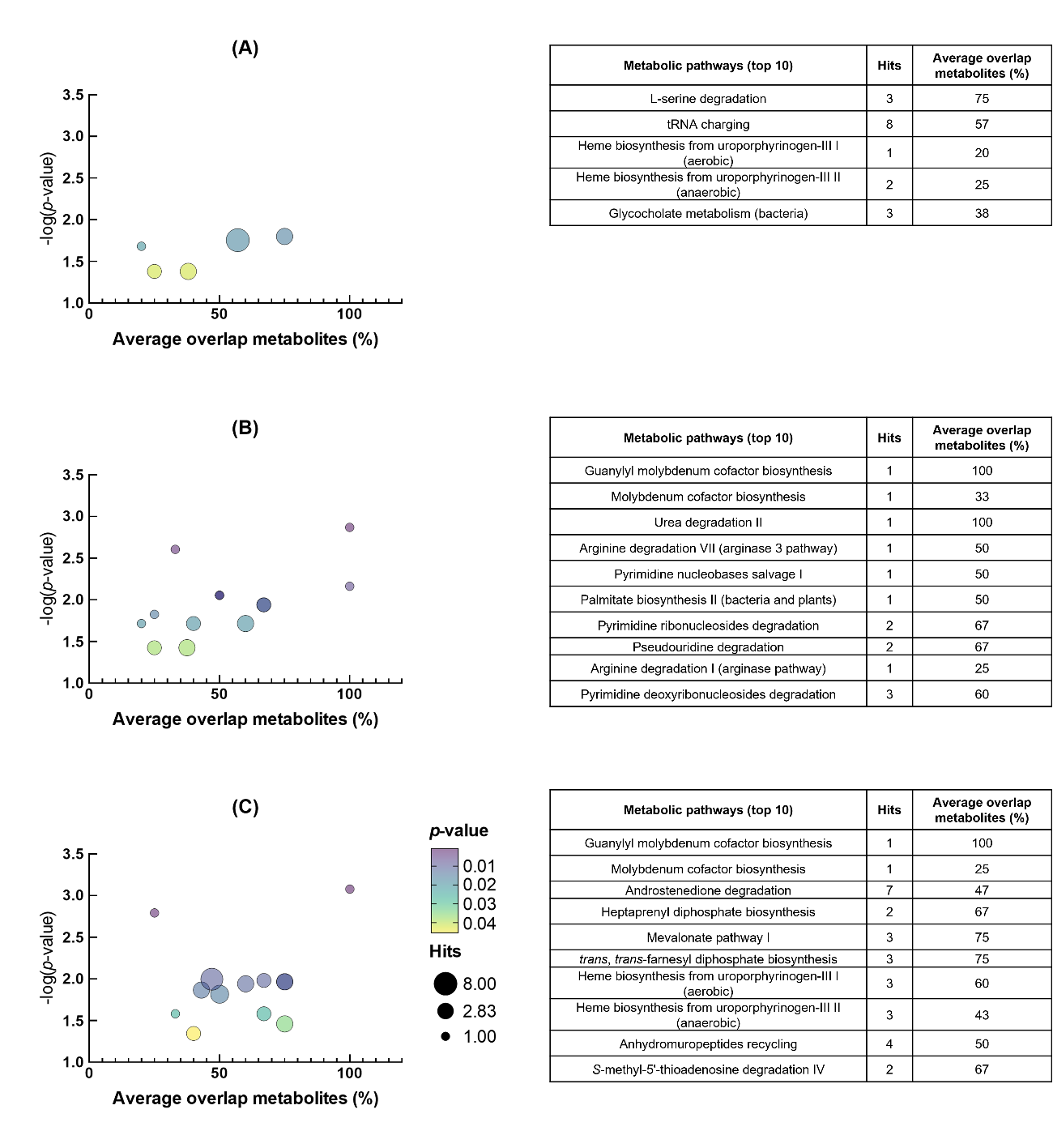

Figure S18– Metabolic pathways significantly altered in MRSA 43300 in the presence of different IC_50_ values of vancomycin.

(A) 0.5xIC_50_, (B) 1xIC_50_, (C) 2xIC_50_ vancomycin. Cloud plots on the left show significantly altered pathways and their metabolite overlap (fraction of identified altered metabolites relative to total number of metabolites that constitute the pathway), as well as the number of hits per pathway. On the right, the top 10 altered pathways are listed. Statistical significance: A and C, *p* < 0.05, and B, *p* < 0.02.

Table S1. Parameters used for the analysis of pairwise jobs processing in XCMS.

| Step | Option | Value |
| --- | --- | --- |
| Feature detection | method | centWave |
|  | ppm | 10 |
|  | minimum peak width | 5 |
|  | maximum peal width | 20 |
|  | mzidff | 0.01 |
|  | signal/noise threshold | 6 |
|  | integration method | 1 |
|  | prefilter peaks | 3 |
|  | prefilter intensity | 100 |
|  | noise filter | 100 |
| Retention time correction | method | obiwarp |
|  | profstep | 1 |
| Alignment | bw | 5 |
|  | minfrac | 0.5 |
|  | mzwid | 0.015 |
|  | minsamp | 1 |
|  | max | 100 |
| Statistics | statistical test | Unpaired parametric t-test (Welch t-test) |
|  | perform post-hoc analysis | True |
|  | *p*-value threshold (highly significant features) | 0.001 |
|  | fold-change threshold (highly significant features) | 1.5 |
|  | *p*-value threshold (significant features) | 0.05 |
|  | value | into |
| Annotation | ppm | 5 |
|  | *m/z* absolute error | 0.015 |
|  | search for | Isotope + adducts |
| Identification | adducts | all |
|  | ppm | 10 |
|  | biosource | SAUR282458 |
|  | pathway ppm deviation | 5 |
|  | significant list *p*-value cutoff | AUTO |
| Visualization | Value | 200 |

Table S2. Multi-omics evidence for metal/cofactor-related responses across antibiotic treatments in MRSA**.**

Summary of omics and enrichment-based evidence related to metal homeostasis and metal-dependent pathways after exposure to ampicillin, chloramphenicol, ciprofloxacin, methicillin, and vancomycin at 0.5, 1, and 2× IC_50_. For each condition, the table reports the direction of change and corresponding log_2_(FC) values together with relevant functional annotations and enrichment terms. ↑, up-regulated; ↓, down-regulated; P, proteomics, E, enrichment; M, metabolomics; AMP, ampicillin; CHL, chloramphenicol; CIP, ciprofloxacin; MET, methicillin; VAN, vancomycin; MoCo, molybdenum cofactor; MOSC, MoCo sulfurase C-terminal; p, phosphorylation; succ, succinylation.

| Antibiotic | Experimental Conditions | Molecular Change (log_2_ FC) | Feature |
| --- | --- | --- | --- |
| AMP | P, 2× IC_50_ | ↑ (+1.64) | Inositol monophosphatase family protein (Q2FVV7) |
|  | P, 2× IC_50_ | ↓ (-2.20) | Fe periplasmic-binding domain-containing protein (Q2FWN6) |
|  | E, 2× IC_50_ | Enriched | Siderophore-dependent iron import into cell |
|  | PTM, 0.5, 2× IC_50_ | ↑ pSer3 (+1.58, +1.81) | MOSC domain-containing protein (Q2FVS9) |
|  | M, all | ↑ (+4.53 to +6.80) | MoO_2_-molybdopterin cofactor |
|  | M,E, all | Enriched | Guanylyl MoCo biosynthesis |
| CHL | P, 0.5× IC_50_ | ↑ (+1.56) | FecCD-family ABC transporter permease (Q2FW76) |
|  | P, 0.5× IC_50_ | ↑ (+2.02) | CobW C-terminal domain-containing protein (Q2G0W4) |
|  | P, 1× IC_50_ | ↑ (+2.67) | UPF0358 protein SAOUHSC_01062 (Q2G2C3) / Fpa (YlaN) |
|  | P, 2× IC_50_ | ↑ (+1.60) | Urease subunit gamma (ureA, Q2FVW5) |
|  | E, 2× IC_50_ | Enriched | Nickel cation binding |
|  | M, all | ↑ (+0.98 to +2.57) | MoO_2_-molybdopterin cofactor |
|  | M, E, all | Enriched | Guanylyl MoCo & MoCo biosynthesis |
| CIP | P, 0.5× IC_50_ | ↓ (-1.54) | Ferrous iron transport protein B (Q2FV72) |
|  | P, all | ↓ (-1.62 to -2.03) | Uncharacterized protein (Q2FXA9) / Putative metal binding site |
|  | M, 2× IC_50_ | ↑ (+3.61) | MoO_2_-molybdopterin cofactor |
|  | M, E, 2× IC_50_ | Enrichment | Guanylyl MoCo biosynthesis |
|  | M, E, 2× IC_50_ | Enrichment | Anaerobic heme biosynthesis |
| MET | P, 2× IC_50_ | ↑ (+1.50) | Urease subunit gamma (ureA, Q2FVW5) |
|  | P, 0.5× IC_50_ | ↑ (+3.02) | Molybdenum ABC transporter (Q2FVX4) |
|  | P, all | ↓ (-1.62 to -1.71) | Periplasmic binding protein (Q2G1N4) / SirA |
|  | E, 0.5× IC_50_ | Enrichment | Siderophore iron import into cell, Cellular iron homeostasis |
|  | E, 1× IC_50_ | Enrichment | Siderophore iron import, Cellular iron homeostasis, Transition metal ion transport |
|  | M, all | ↑ (+1.96 to +3.32) | MoO_2_-molybdopterin cofactor |
|  | M, all | ↓ (-4.74 to -5.02) | Protoporphyrinogen IX |
|  | M, E, 0.5-1× IC_50_ | Enrichment | Aerobic and anaerobic heme biosynthesis |
|  | M, E, 2× IC_50_ | Enrichment | Guanylyl MoCo biosynthesis |
| VAN | P, 1-2× IC_50_ | ↑ (+2.64, +1.90) | Putative hemin transport system permease protein HrtB (Q2G168) |
|  | P, 2× IC_50_ | ↓ (-2.25) | Nickel-binding protein NikA (nikA, Q2G2P5) |
|  | E, 2× IC_50_ | Enrichment | Nickel cation transport, Metal ion transport, Cation transport |
|  | PTMs, 0.5, 2× IC_50_ | ↓ succLys174 (-2.18, -2.43) | YdhG-like domain-containing protein (Q2FVG5) |
|  | M, 1-2× IC_50_ | ↑ (+7.23, +7.12) | MoO_2_-molybdopterin cofactor |
|  | M, E, 1-2× IC_50_ | Enrichment | Guanylyl MoCo & MoCo biosynthesis |
|  | M, E, 0.5, 2× IC_50_ | Enrichment | Heme biosynthesis |

Table S3. Multi-omics evidence for convergent perturbation of nucleotide and folate metabolism across antibiotic treatments in MRSA**.**

Summary of omics and enrichment-based evidence related to nucleotide metabolism, folate-dependent precursor supply, and associated stress-response functions after exposure to ampicillin, chloramphenicol, ciprofloxacin, methicillin, and vancomycin at 0.5, 1, and 2× IC_50_. For each condition, the table reports the direction of change and corresponding log_2_(FC) values together with relevant functional annotations and enrichment terms. ↑, up-regulated; ↓, down-regulated; P, proteomics, E, enrichment; M, metabolomics; L, lipidomics; AMP, ampicillin; CHL, chloramphenicol; CIP, ciprofloxacin; MET, methicillin; VAN, vancomycin; MoCo, molybdenum cofactor**;** f-GAR, *N*^2^-formyl-*N*^1^-(5-phospho-β-d-ribosyl)glycinamide; p, phosphorylation; ac, acetylation.

| Antibiotic | Experimental Conditions | Molecular Change (log_2_ FC) | Feature |
| --- | --- | --- | --- |
| AMP | M, L, 0.5-1× IC_50_ | ↓ (-3.71, -4.46) | f-GAR |
|  | M, L, 2× IC_50_ | ↑ (+6.28) | Adenylosuccinate |
|  | P, 0.5 IC_50_ | ↑(+3.32) | Ribonucleoside-diphosphate reductase (Q2G078) |
|  | L, E, all | Enrichment | Tetrahydrofolate salvage |
|  | M, L, 0.5, 1× IC_50_ | ↓ (-3.71, -4.46) | f-GAR |
| CHL | L, M, all | ↓ ( -3.60 to ‑4.25) | f-GAR |
|  | P, 1, 2× IC_50_ | ↑ (+2.74, +2.57) | Protein NrdI (nrdI, Q2G079) |
|  | L, E, all | Enrichment | Tetrahydrofolate salvage |
|  | M, E, 0.5× IC_50_ | Enrichment | UTP/CTP dephosphorylation I pathway |
| CIP | L, E, 0.5, 2× IC_50_ | Enrichment | Tetrahydrofolate salvage |
|  | M, L, 0.5, 2× IC_50_ | ↓ (-3.18, -5.06) | f-GAR |
|  | P, all | ↓ (-1.53 to -2.25) | Aspartokinase (Q2FYP1) |
|  | M, E, all | Enrichment | Purine degradation/salvage, de novo purine synthesis |
|  | PTM, 1, 2× IC_50_ | ↓ p H14 (-2.46, -2.04) | UvrABC system protein A (*uvrA*, Q2G046) |
| MET | L, E, 0.5, 2× IC_50_ | Enrichment | Tetrahydrofolate salvage |
|  | L, M, 0.5, 2× IC_50_ | ↓ (-3.56, -3.71) | f-GAR |
|  | L, M, 2×IC_50_ | ↑ (+5.14) | Adenylosuccinate |
|  | M, E, 0.5, 2× IC_50_ | Enrichment | Pyrimidine/Purine deoxyribonucleosides degradation |
| VAN | P, 2× IC_50_ | ↓ (-2.19) | Aspartokinase (Q2FYP1) |
|  | PTMS, 1, 2× IC_50_ | ↓ acK291, acK299 (-1.52 to ‑1.95) | CTP synthase (pyrG, Q2FWD1) |
|  | M, E, 1× IC_50_ | Enrichment | Pyrimidine salvage/degradation pathways |
|  | PTMS, 0.5, 2× IC_50_ | ↓ p (-4.59 to ‑5.35) | ADP-dependent (S)-NAD(P)H-hydrate dehydratase (*nnrD*, Q2G2P8) |

****Table S4. Multi-omics evidence for coordinated DNA repair, replication control, and prophage-associated responses across antibiotic treatments in MRSA.****

Summary of omics and enrichment-based evidence related to genome maintenance, replication control, and prophage-associated functions after exposure to ampicillin, chloramphenicol, ciprofloxacin, methicillin, and vancomycin at 0.5, 1, and 2× IC50. For each condition, the table reports the direction of change and corresponding log2(FC) values together with relevant functional annotations and enrichment terms. ↑, up-regulated; ↓, down-regulated; P, proteomics, E, enrichment; M, metabolomics; L, lipidomics; AMP, ampicillin; CHL, chloramphenicol; CIP, ciprofloxacin; MET, methicillin; VAN, vancomycin; p, phosphorylation; ac, acetylation.

| Antibiotic | Experimental Conditions | Molecular Change (log_2_ FC) | Feature |
| --- | --- | --- | --- |
| AMP | P, 0.5, 2× IC_50_ | ↑ (+2.80, +3.04) | DNA ligase (*ligA*, Q2G1Y0) |
|  | P, 0.5× IC_50_ | ↑ (+1.94) | DNA repair protein RadA (*radA*, Q2G243) |
|  | P, 0.5, 1× IC_50_ | ↑ (+1.82, +1.75) | Phage portal protein, HK97 family (Q2FYC7) |
|  | PTM, 1, 2× IC_50_ | ↑ acK570 (+1.55, +1.94) | 3'-5' exonuclease DinG (*dinG*, Q2FYH5) |
|  | E, 0.5, 1× IC_50_ | Enrichment | Nucleotide excision repair components, DNA metabolism |
| CHL | P, 0.5, 1x IC_50_ | ↑ (+1.95, +2.54) | DNA repair/chromosome segregation ATPase (Q2FVZ7) |
|  | P, 2× IC_50_ | ↑ (+2.02) | Chromosome partition protein Smc (*smc*, Q2FZ49) |
|  | P, all | ↑ (+3.46 to +2.89) | Conserved / Hypothetical phage proteins (Q2FY93, Q2FY90) |
|  | E, 0.5× IC_50_ | Enrichment | DNA restriction-modification system, DNA methylation on adenine |
| CIP | P, all | ↓ (-1.94 to -2.99) | Phage terminase, large subunit, putative (Q2FWT4) |
|  | P, 1× IC_50_ | ↑ (+2.24) | Initiation-control protein YabA (Q2G2W8) |
|  | P, 0.5× IC_50_ | ↑ (+1.71) | Site-specific DNA-methyltransferase (adenine-specific) (Q2FXD0) |
|  | PTM, 1, 2× IC_50_ | ↓ pH14 (-2.46, -2.04) | UvrABC system protein A (uvrA, Q2G046) |
|  | E, 0.5× IC_50_ | Enrichment | DNA restriction-modification system |
| MET | P, 1, 2× IC_50_ | ↑ (+4.02, +3.98) | ATP-dependent helicase/deoxyribonuclease subunit B (addB, Q2FZT6) |
|  | P, 0.5× IC_50_ | ↑ (+2.52) | DNA replication and repair protein RecF (Q2G275) |
|  | P, 2× IC_50_ | ↑ (+5.16) | Phage head protein, putative (Q2FX56) |
|  | P, 1, 2× IC_50_ | ↑ (+1.98, +1.96) | Integrase (Q2G108) |
| VAN | P, 1× IC_50_ | ↑ (+2.89) | DNA polymerase beta (Q2FZD4) |
|  | P, 0.5× IC_50_ | ↑ (+1.60) | DNA-directed DNA polymerase (Q2G0T5) |
|  | P, 0.5, 1× IC_50_ | ↓ (-1.68, -1.57) | Hypothetical phage protein (Q2FWS9) |

Table S5. Multi-omics evidence for coordinated remodeling of transport systems, surface-associated functions, and virulence regulation across antibiotic treatments in MRSA**.**

Summary of omics and enrichment-based evidence related to transport, surface-associated functions, envelope remodeling, and virulence regulation after exposure to ampicillin, chloramphenicol, ciprofloxacin, methicillin, and vancomycin at 0.5, 1, and 2× IC_50_. For each condition, the table reports the direction of change and corresponding log_2_(FC) values together with relevant functional annotations and enrichment terms. ↑, up-regulated; ↓, down-regulated; P, proteomics, E, enrichment; AMP, ampicillin; CHL, chloramphenicol; CIP, ciprofloxacin; MET, methicillin; VAN, vancomycin; MoCo, molybdenum cofactor**;** p, phosphorylation; succ, succinylation.

| Antibiotic | Experimental Conditions | Molecular Change (log_2_ FC) | Feature |
| --- | --- | --- | --- |
| AMP | P, 1, 2× IC_50_ | ↑ (+6.10, +6.11) | Uncharacterized protein (Q2FW89) / TcaR-like |
|  | P, all | ↓ (-2.74 to -2.99) | Polysaccharide biosynthesis protein (Q2G0R7) |
|  | PTM, 0.5, 1× IC_50_ | ↑ succK11 (+1.54, +1.74) | Oligopeptide ABC transporter (Q2FZR0) |
|  | P, 0.5, 1× IC_50_ | ↑ (+1.61, +1.66) | FKLRK protein (Q2G0X3) |
| CHL | P, 1, 2× IC_50_ | ↑ (+1.57, +2.66) | Amino acid ABC transporter (Q2FX87) |
|  | P, 2× IC_50_ | ↑ (+2.36) | PTS system lactose-specific EIICB component (Q2G2D4) |
|  | P, 2× IC_50_ | ↑ (+2.23) | Surface protein F (sasF, Q2FUW9) |
|  | P, 2× IC_50_ | ↑ (+1.99, +1.73) | HTH araC/xylS-type domain-containing protein (Q2FY65, Q2G0D2) |
| CIP | P, all | ↑ (+2.88, +1.67, +2.39) | Penicillin-binding protein 1 (pbp1, Q2FZ94) |
|  | PTM, all | ↓ p (-1.96 to -2.43); multi-site | ABC transporter domain-containing protein (Q2G196) |
|  | PTM 1, 2× IC_50_ | ↓ succ K90/K97 (-1.74 to  -2.29) | Probable cell wall hydrolase LytN (lytN, Q9ZNI1) |
|  | E, 1× IC_50_ | Enrichment | Phosphotransferase system, Carbohydrate transmembrane transport |
| MET | P, all | ↓ (-2.05 to -2.23) | Accessory gene regulator protein B (agrB, Q2FWM7) |
|  | P, all | ↓ (-1.81 to -1.90) | Sortase A (Q2FVN4) |
|  | P, 1, 2× IC_50_ | ↑ (+2.17, +1.97) | Amino acid ABC transproter, permease protein (Q2FVL3) |
|  | P, 0.5× IC_50_ | ↑ (+2.07) | Type VII secretion system protein EssC (essC, Q2G184) |
| VAN | P, all | ↓ (-1.64 to -1.85) | Regulatory protein YycH domain-containing protein (Q2G2U3) |
|  | P, all | ↓ (-1.93 to -2.31) | Accessory gene regulator protein B (agrB, Q2FWM7) |
|  | P, PTM, all | ↓ (-1.90 to -2.66); multi-site | Polysaccharide biosynthesis protein (Q2G0R7) |
|  | PTM, 1, 2× IC_50_ | ↑ deamidationN200 (+3.45, +3.74) | Energy-coupling factor transporter EcfT (Q2FZI4) |
|  | P, 1× IC_50_ | ↑ (+3.54) | MFS transporter (Q2G2B3) |
